# Diversity and ecological roles of RNA viral communities in the cryosphere of the Tibetan Plateau

**DOI:** 10.1101/2024.08.30.610425

**Authors:** Yongqin Liu, Zhihao Zhang, Nianzhi Jiao, Yongguan Zhu, Rui Zhang, Guillermo Dominguez-Huerta, Haina Wang, Meiling Feng, Rong Wen, Mukan Ji, Qiang Zheng, Pengfei Liu, Tandong Yao

## Abstract

RNA viruses play crucial roles in modulating host communities, but their diversity and ecological roles in the cryosphere are poorly understood. We investigated the RNA virome from the cryosphere of the Tibetan Plateau (TPC), the largest area of the cryosphere in the mid-low latitude regions, which includes 79 metatranscriptomes from ecosystems including glaciers, proglacial lake sediments, permafrost wetland and upland soils. We obtained 8,799 RNA-dependent RNA polymerase encoding contigs, which clustered into 5,333 viral species, adding more than 4,900 new RNA viral species to the global RNA virome database. The recovered RNA viruses primarily infect prokaryotes, distinctively different from the dominance of eukaryotic RNA viruses in oceanic ecosystems. In addition to the known taxa such as *Leviviricetes*, we identified novel clades of putative prokaryotic RNA viruses, which greatly extended our understanding of their diversity. We further expanded the host of RNA virus by 20 additional prokaryotic phylum- or class-level taxonomic groups. Moreover, the TPC RNA viruses encoded dozens of auxiliary metabolic genes that could reprogram the metabolisms of diverse hosts, including photosynthesis in glacier eukaryotic algae and cyanobacteria, which may exert great impacts on glacier dynamics through modulating host adaptation. Furthermore, we identified potential pathogenic RNA viruses in various cryosphere environments, though their overall risks are low. This study provides the first comprehensive view of cryosphere RNA virome, highlighting their high diversity, vital ecological functions, and potential health impacts.

## Introduction

RNA viruses are a type of virus that uses ribonucleic acid (RNA) as their genetic material instead of deoxyribonucleic acid (DNA)(*1, 2*). Recent metatranscriptomic studies have revealed a vast diversity of RNA viruses in natural environments, with their hosts spanning bacteria, fungi, plants, and animals(*3–13*). Environmental RNA viruses play important roles in global biogeochemical cycling and ecosystem function maintenance(*14–16*) beyond their impacts on public health(*17*). They are recognized as a crucial factor in controlling microbial community dynamics by selectively infecting their hosts, impacting host abundance, diversity, metabolism, and genetic composition(*6, 8, 18, 19*). Therefore, understanding the biogeography and genetic characteristics of environmental RNA viruses can provide novel insights into the dynamics and the ecological roles of microbial communities in natural ecosystems(*8, 20, 21*).

With enhanced sequencing efforts and expanded ecosystems studied, the diversity of RNA viruses is rapidly increasing(*4–6, 12*). Currently, over 10^5^ species of RNA viruses that could be assigned to over 180 phyla- or class-level RNA viral superclades(*4–6*) have been revealed. A recent analysis revealed that oceanic RNA viruses can alter various aspects of host metabolism such as photosynthesis and carbon cycling. By using specific auxiliary metabolic genes (AMGs)(*8*), these viruses might enhance their hosts’ environmental adaptability, directly influencing ecosystem carbon dynamics(*8*). Additionally, certain RNA viruses that infect marine green algae were proved to be predictors of ocean carbon fluxes(*8*). The size of the RNA virosphere has been speculated to be near 2 trillion (2×10^12^) RNA virus species(*20*). Therefore, further exploration of diverse natural environments through advanced sequencing techniques may unveil additional hidden RNA viruses, offering new insights into their evolutionary processes and ecological functions.

The Tibetan Plateau (TP) owns the largest area of cryosphere in the mid-low latitude region(*22*). Microbes are the main forms of life that have adapted to harsh conditions like intense UV radiation, extreme temperature fluctuations, and low atmospheric pressure(*23*) of the TP cryosphere (TPC). However, except for our recent work on one alkaline saline lake of the TPC(*11*), the RNA viral communities of the TPC environment have rarely been explored. Hence, the diversity, biogeography, and functions of TPC RNA viruses are largely unknown. Moreover, the cryosphere may serve as a reservoir for pathogenic viruses that could be released after it thaws due to global warming(*24, 25*). The TP and its surrounding regions store the largest amount of frozen water and is known as the “Asian Water Tower”, which provides a reliable water supply to nearly two billion people(*26*). The TP is also highly sensitive to climate change, experiencing rapid warming at twice the global average rate(*26*). Therefore, whether the TPC contains pathogenic RNA viruses have received wide public attention(*24, 25, 27*). Thus, a comprehensive examination of RNA viruses in the TPC is crucial to improving our ability to develop prevention strategies against the transmission of potential pathogenic RNA virus.

Here, we sought to explore the diversity, ecological functions and potential pathogenic risks of TPC RNA viruses. We performed metatranscriptome sequencing of 79 samples from the TPC (Fig. 1A&B; table S1), covering habitats of glacial environments (including snow, ice, and cryoconite-a granular, dark-colored sediment on the glacial surface), sediments of a proglacial lake, and soil from permafrost regions (including wetland and upland). We identified RNA viruses through mining RNA-dependent RNA polymerases (RdRps), the single hallmark protein that is shared by all orthornavirans, *i.e*., all RNA viruses belonging to the riboviriad kingdom *Orthornavirae*(*16, 28–31*), and required for replication. We also inferred the RNA viruses encoded AMGs through functional annotation, and assessed their pathogenic risks through host assignments and genome feature-based prediction. We hypothesize that the diversity of RNA viruses in the TPC would be distinct from those in other environments due to the unique environmental conditions such as intense UV radiation, extreme temperature fluctuations, and low atmospheric pressure, and this unique RNA viral community may play a crucial role in shaping microbial dynamics and influencing biogeochemical processes in the TPC ecosystem.

**Fig. 1.**
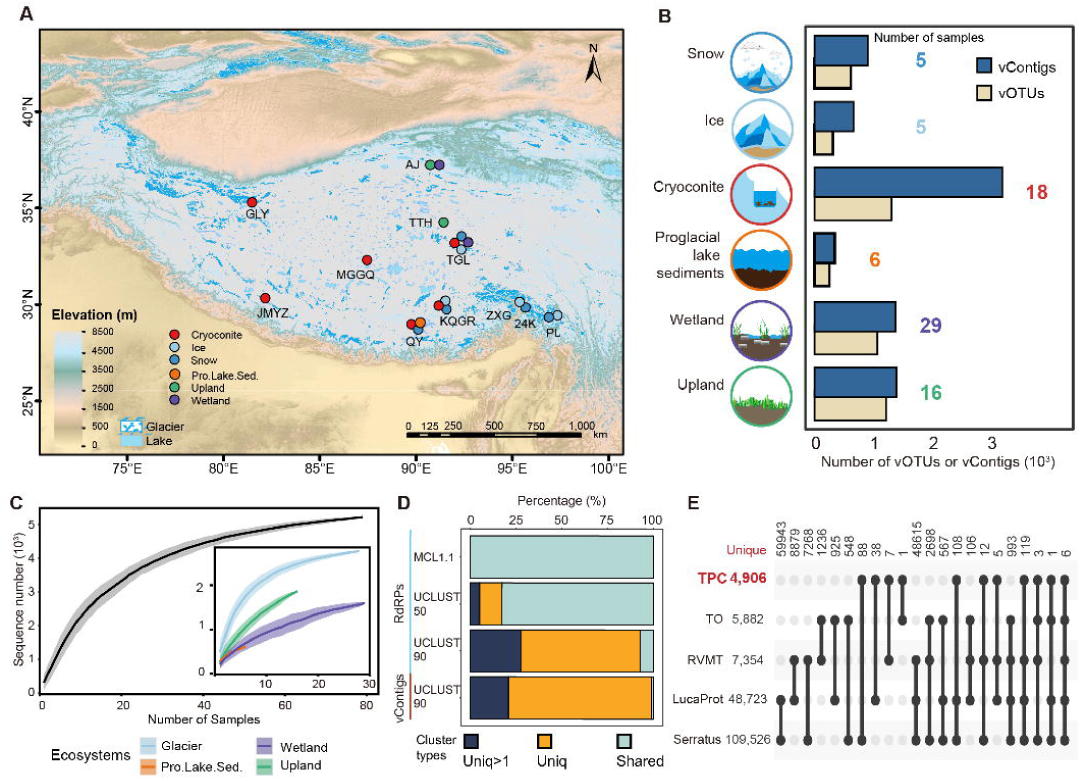
Overview of the TPC RNA virome. (**A**) Overview of metatranscriptome sampling sites across the Tibetan Plateau. (**B**) Sample sources and the number of vContigs and vOTUs obtained from each habitat. (**C**) RdRp90 accumulation curve, the accumulation of unique clusters as a function of the number of samples analyzed. (**D**) Number of vContigs shared with reference datasets or unique to the TPC RNA virome. Shared means that the TPC vContigs were members of clusters with vContigs from reference datasets including the *Tara* Oceans, the RVMT, the Serratus, and the LucaProt according to different clustering strategies. (**E**) Upset plot showing the number of vOTUs shared between the TPC dataset with the *Tara* Oceans, the RVMT, the Serratus, and the LucaProt datasets. TPC only, vContigs were members of clusters with vContigs only from the TPC metatranscriptomes.

## Results

### RNA viral communities of the TPC

We obtained 8,799 RNA viral contigs (vContigs) with length ≥1 kb (Fig. 1A, fig. S1; table S2). Based on an RdRp amino acid sequence identity cutoff of 90%, these vContigs were clustered into 5,333 species-level virus operational taxonomic units (vOTUs; Fig. 1B; table S2). The diversity of the TPC RNA viral species was well represented by our current sampling efforts as evaluated by the accumulation curve (Fig. 1C). Compared to global RNA virome datasets (Fig. 1D), the TPC RNA viruses with full-length RdRps (3,045, 34.6% of total TPC RdRps) all belonged to established phyla (Fig. 1D). At family-to subfamily-level(*8*), TPC RNA vContigs were grouped into 3,176 clusters, and 549 clusters of them (17.3%) contained only the TPC RdRps, while the rest 2,627 clusters (82.7%) were also identified in the global RNA virome datasets (e.g., the RVMT(*6*); Fig. 1D, fig. S3 and S4). However, at the species level, only 425 (8.0% of 5,333) vOTUs were shared with viral genomes from the global RNA virome datasets, with 4,908 vOTUs (92.0%) being TPC unique (Figs. 1D and 1E, fig. S2).

Consensus taxonomic classification of RNA viral genomes based on both RdRps and vContigs showed that 6,893 vContigs (78.3% of 8,799) could be assigned to 6 established viral phyla, with the remaining 1,906 (21.7%) vContigs could not be classified (Fig. 2A, figs. S5-S8; table S3). Classified vOTUs are taxonomically affiliated with *Lenarviricota* (3,146 vOTUs, 59.0% of 5,333), *Kitrinoviricota* (481, 9.0%), *Pisuviricota* (380, 7.1%), *Duplornaviricota* (103, 1.9%), *Negarnaviricota* (60, 1.1%), and p.0002.base-Kitrino(*6*) (2, 0.038%) (Fig. 2A, fig. S5). These vContigs could be further assigned to 130 established families according to the RVMT framework(*6*) (Fig. 2A; table S3). Families with the top most vContigs were mainly from the phylum *Lenarviricota*, including the families of *Steitzviridae* (994 vOTUs, 18.6% of total vOTUs), *Fiersviridae* (725, 13.6%), *Atkinsviridae* (130, 2.4%), *Botourmiaviridae* (426, 8.0%) and *Mitoviridae* (338, 6.3%). Other families with more than 80 vOTUs included *Tombusviridae* (160, 3.0%), *Narnaviridae* (144, 2.7%) and *Partitiviridae* (83, 1.6%) (Fig. 2A, fig. S6C; table S3).

**Fig. 2.**
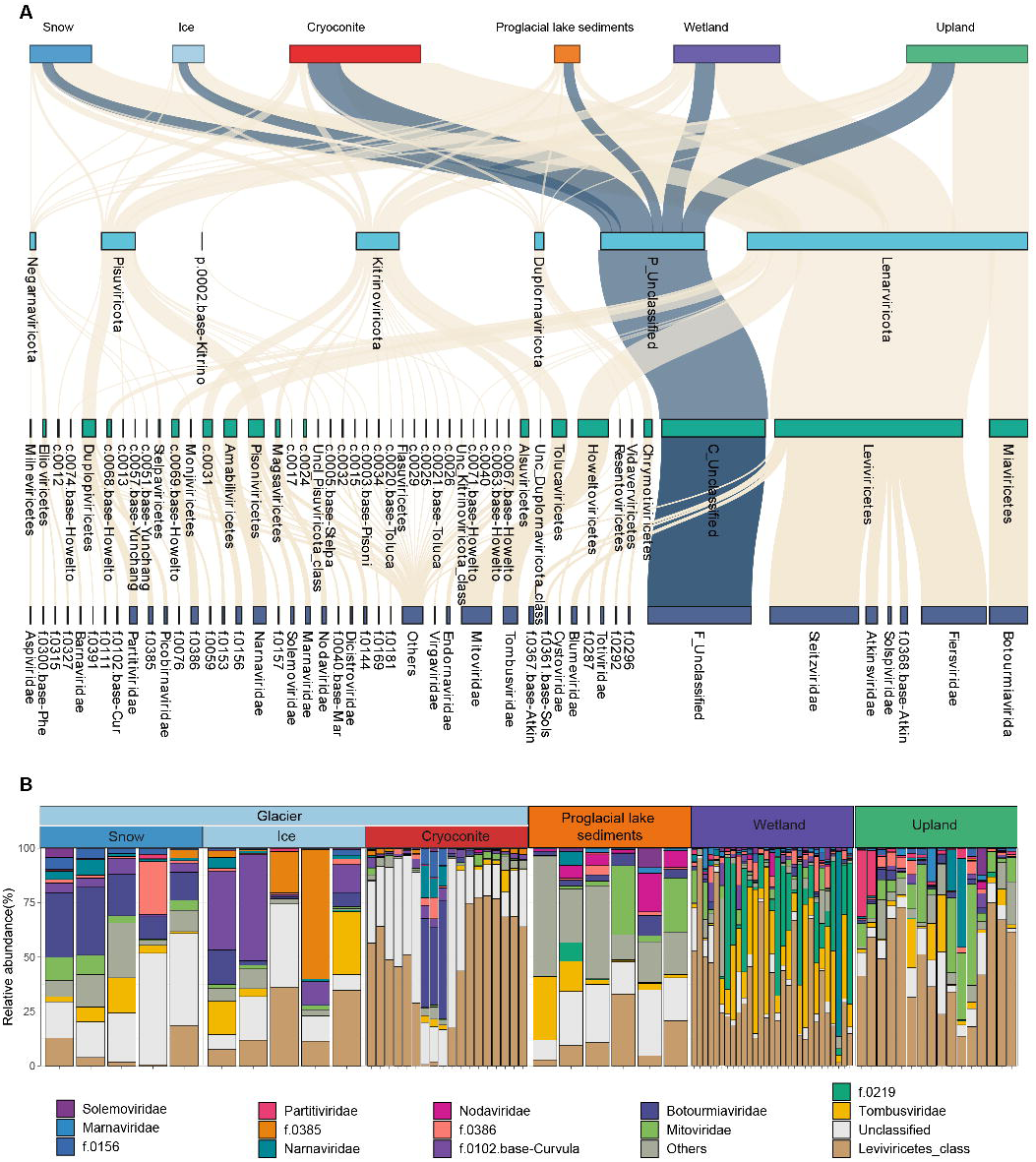
RNA viral community structure of the TPC. (**A**) Sanky plot showing the distribution of vOTUs in different RNA viral taxonomic ranks. Un., unclassified, C_ or c., class; f., family. (**B**) Relative abundance of RNA viral families across different habitats. All families belonging to the *Leviviricetes* class are grouped and the relative abundance of each *Leviviricetes* family (e.g., *Steitzviridae*) is shown in fig. S7. Pro. Lake Sed., proglacial lake sediments.

Dominant TPC RNA viral families and their relative abundance varied across the ecosystems and habitats (Fig. 2B). The *Leviviricetes* class is the most abundant taxonomic group (Fig. 2B), ranging from 0.1% (minimum in cryoconite) to 95.0% (maximum in wetland). The *Leviviricetes* were dominant (relative abundance >30%) in 53.2% of all samples (Fig. 2B), especially in cryoconite (72.2% of all cryoconite samples), wetland (44.8%), and upland (85.7%). In addition, viruses of the families *Steitzviridae*, *Fiersviridae*, *Atkinsviridae*, and f.0368.base-Atkins are the most abundant within the *Leviviricetes* class (fig. S8). The relative abundance of these families reached up to 95.0% (in wetland), 77.7% (cryoconite), 75.1% (upland), and 18.6% (snow), respectively. Other *Leviviricetes* (e.g., *Solspiviridae*) are less abundant, reaching only a maximum of 10% across all samples (fig. S8). Dominant RNA viral groups (with maximum relative abundance >10% across all samples) outside *Leviviricetes* mainly included *Tombusviridae*, *Botourmiaviridae*, *Mitoviridae*, *Flaviviridae*, and *Nodaviridae* (Fig. 2B). Finally, we found unclassified RNA viral families also accounted for a large proportion of viral communities, especially in glacier habitats: cryoconites (maximum 77.6%; mean 30.7%), ice (38.2%; 16.8%) and snow (51.2%; 29.8%), respectively (Fig. 2B).

### Diversity of the TPC RNA viral communities

Comparison of taxa across habitats revealed the high diversity of TPC RNA viruses. To explore local diversity (i.e., per-sample) of the TPC RNA communities, the alpha-diversity was accessed by richness and Shannon’s H indices (Fig. 3A). The observed viral richness was between 66 and 998 (mean ranged from 244 in wetland to 534 in cryoconite) and the Shannon’s H was between 1.2 and 7.3 (mean ranged from 4.1 in cryoconite to 5.8 in snow). The richness of the RNA viral communities of the glacier habitats was slightly higher than other ecosystems while the Shannon’s H was similar among all ecosystems and habitats (Fig. 3A). The TPC RNA viruses showed remarkable ecosystem specificity and varied significantly across different ecosystems and habitats (PERMANOVA, Habitat *R^2^*=0.23; Ecosystem *R^2^*=0.174, both *P<*0.001; Fig. 3B). The specificity of the TPC RNA viral communities was validated by the pairwise PERMANOVA analysis (all pairwise *P*<0.001; fig. S9).

**Fig. 3.**
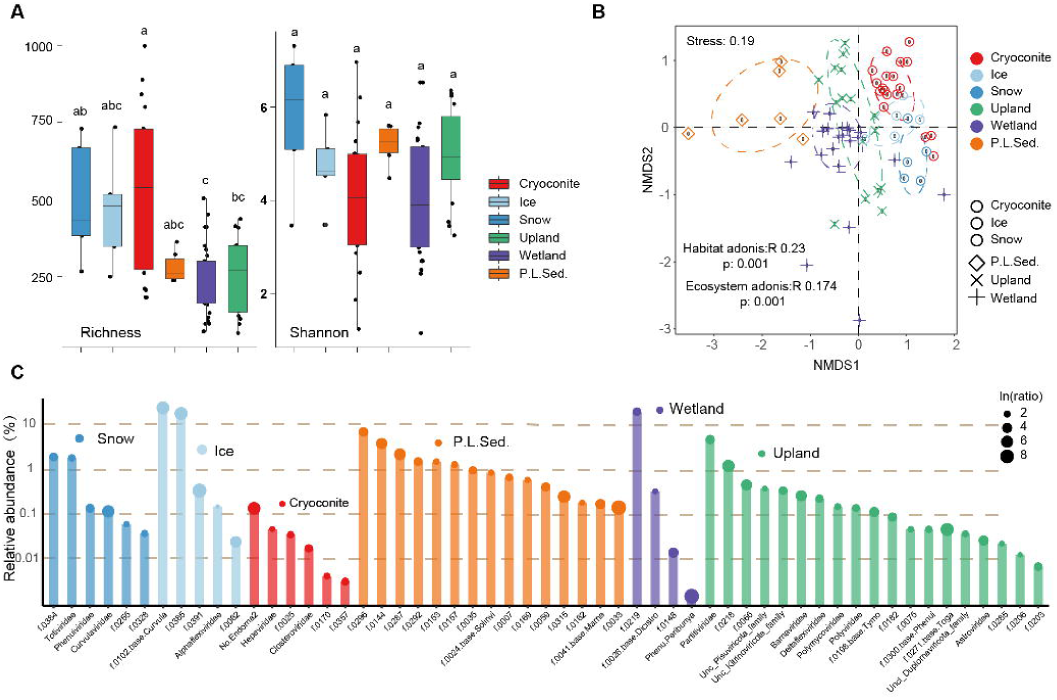
Diversity of The TPC RNA viral communities. (**A**) Alpha-diversity of the TPC RNA viral communities. (**B**) Beta-diversity of the TPC RNA viral communities. (**C**) Lollipop plot showing the average relative abundance of habitats enriched viral families and the environments in which they were enriched. The size of the dots on top of the bar indicates the folds of the average relative abundance of the corresponding family in the enriched environments compared to that of the environment with the second highest values.

Since RNA viral communities showed a high habitat dissimilarity, we investigated the habitat-enriched RNA viral families (Fig. 3C). We found that 55 (41.3% of all 130 families) RNA viral families were enriched in certain habitats, including 6, 5, 6, 15, 4, and 19 for snow, ice, cryoconites, proglacial lake sediments, wetland, and upland, respectively (Fig. 3C). However, most of these enriched RNA viral families were rare groups (relative abundance <1%), with 42 viral families (76.4%) having a relative abundance <1%, while only 13 of them (23.6% of all 55) having a relative abundance >1% in the enriched environments (Fig. 3C).

### Potential hosts of TPC RNA viruses

Screening for the hosts of RNA viruses is challenging due to the lack of cultivated virus-host interaction systems for most environmental RNA viruses. Hence, we first assessed the hosts of RNA viruses by using the host information available for viruses of established taxa. For the vOTUs with a family-rank taxonomic assignment (4,160), 3,762 (70.5% of 5,333 vOTUs) could be linked to the putative hosts, including eukaryotes (29.8%; 1,587 of 5,333) and prokaryotes (40.8%; 2,175 of 5,333) according to the virus-host databases, ICTV RNA meta-information, the RVMT(*6*), and the *Tara* Oceans datasets(*5*) (Fig. 4A, fig. S10; table S3). Remarkably, for those with host assignments, 57.8% of vOTUs (2,175/3,762) were classified as bacteriophages, while only 9.15%, 1.08%, 0.7%, 0.05%, and 1.19% were assigned to the eukaryotic viruses of plants/fungi, protists, invertebrates, vertebrates, and animals, respectively (Fig. 4B, fig. S10). The prediction of the TPC RNA viral hosts by the taxonomic assignment was further supported by other *in-silico* methods including the search of (i) endogenous virus elements (EVEs) across reference cellular genomes, (ii) movement protein of plant RNA viruses, and (iii) hallmark genes of prokaryotic RNA viruses (table S4; see below for more details).

**Fig. 4.**
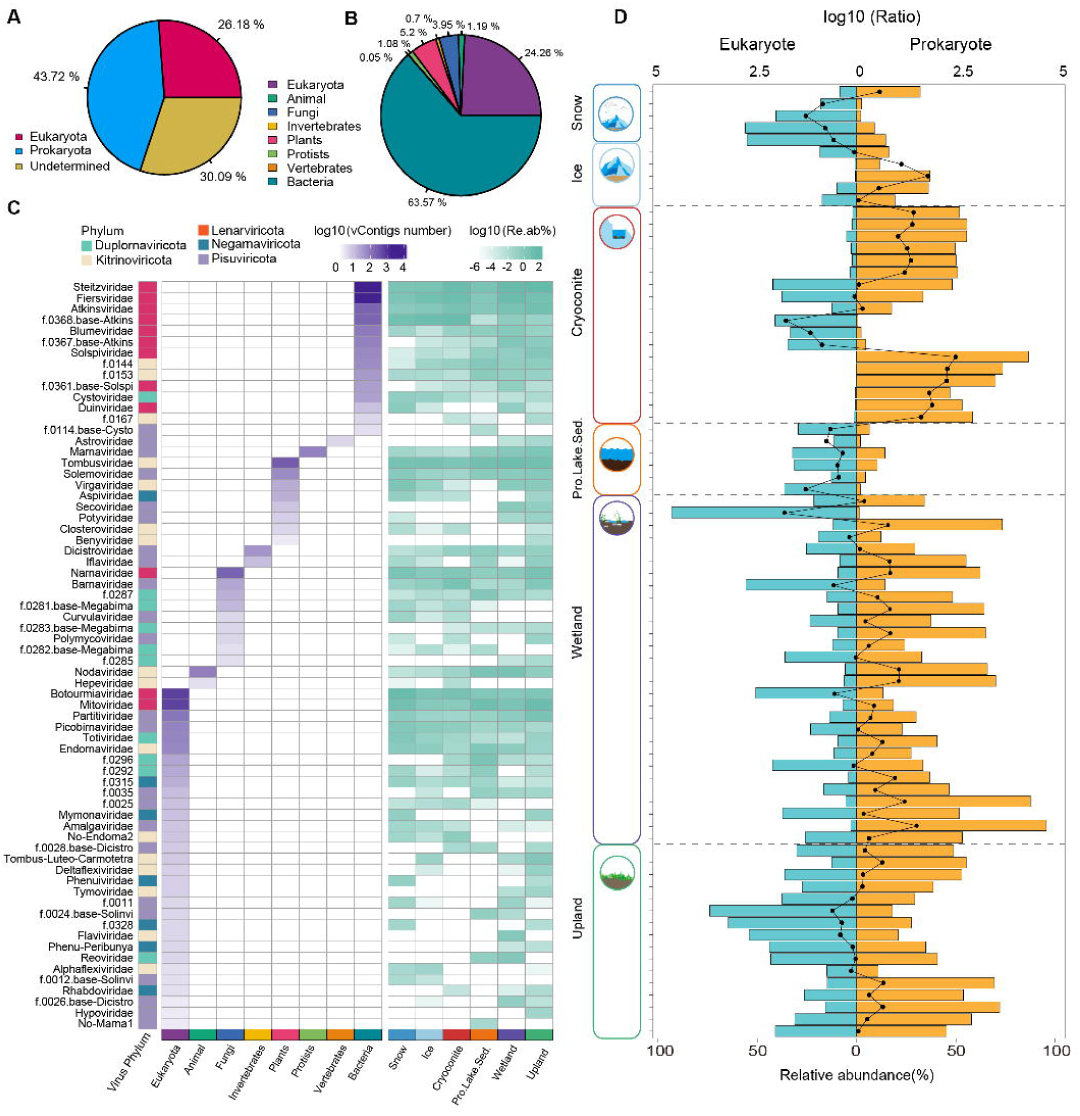
Viral-host linkage of the TPC RNA viral communities. **(A)** and (**B**) Composition of host assigned to the TPC vContigs based on vContigs taxonomic classification. (**C**) Numbers (left heatmap) and the relative abundance (right heatmap) of canonical eukaryotic and prokaryotic vContigs belonging to different RNA viral families. (**D**) Comparison of the relative abundance of eukaryotic and prokaryotic RNA viral families across samples.

The vContigs distribution and their relative abundance among different host families are summarized in Fig. 4C. Among all the 68 established RNA viral families with host assignments, 14 and 54 of them were predicted to infect prokaryotes and eukaryotes (table S3), respectively. The dominant groups of eukaryotic RNA viruses included *Botourmiaviridae* (maximum mean relative abundance 17.4% in snow), *Tombusviridae* (9.6%, wetland), and *Mitoviridae* (14.3%, upland), all of which had a mean relative abundance >5% in at least one type of habitats (Fig. 4C). The dominant groups of prokaryotic RNA viruses were from *Leviviricetes* (e.g., *Steitzviridae*) as described above (Fig. 2B, fig. S8). In addition, dsRNA phages of *Cystoviridae* were also detected across all the habitats, with mean relative abundances ranging from 0.0005% (proglacial lake sediments) to 0.2% (snow) (Fig. 4C).

Ecologically, the TPC RNA viral communities were dominated by the prokaryotic RNA viruses in ice, cryoconite, wetland, and upland while those were dominated by eukaryotic RNA viruses in snow and proglacial lake sediments (Fig. 4D). The relative abundance ratio of the prokaryotic to eukaryotic RNA viruses were 311 (in cryoconite), 62 (ice), 4.7 (upland), and 33 (wetland), respectively, while the ratio of eukaryotic to prokaryotic RNA viruses was 19 in both snow and the proglacial lake sediments. The cryoconite RNA viral communities of JMYZ glacier was an exception, which was also dominated by eukaryotic RNA viruses, and the ratio of eukaryotic to prokaryotic viruses varied between 7.5 and 60.1 (Fig. 4D).

### Expansion of the bacterial and archaeal RNA viruses and their hosts

To identify prokaryotic RNA viruses beyond the viral taxonomic assignment, we searched the nucleotide sequences of TPC vContigs against CRISPR-spacers from the prokaryotic genome of our TP datasets(*23*) and the iPHoP database(*32*). We detected 304 unique linkages between 208 prokaryotic genomes (201 from the iPHoP database and 7 from TPC) and 280 TPC vContigs (Figs. 5A to 5C). Metagenome assembled genomes (MAGs) with at least one potential CRISPR-spacer linked to the RNA viruses belonged to 23 prokaryotic phyla (Fig. 5C; table S5). The top 8 phyla with the highest number of MAGs included Firmicutes (58 MAGs), Bacteroidota (54), Proteobacteria (48), Actinobacteriota (16), Cyanobacteria (6), Fusobacteriota (6), Verrucomicrobiota (3), and Myxococcota (3) (Fig. 5C). For these 280 RNA vContigs with putative host linkages, 215 (76.8%) belonged to 38 families mainly including typically “bacterial” [*Fiersviridae* (57), *Steitzviridae* (45)], and “eukaryotic” ones [*Mitoviridae* (19), *Botourmiaviridae* (12), f.0156 (5), and *Endornaviridae* (8)] (Fig. 5A; table S5). Besides, 65 unclassified vContigs can also be linked to prokaryotic MAGs, which accounted for 23.2% of total vContigs with CRIPSR-spacer links (Fig. 5A; table S5). Remarkably, we found five vContigs linked to four archaeal MAGs (Fig. 5B). The five RNA viruses belonging to *Fiersviridae*, *Steitzviridae*, f.0144, *Endornaviridae*, and *Picobirnaviridae* were putatively linked to two Halobacteriota (*Halobellus* sp. and *Methanoregulaceae* UBA467 sp.), one Thermoproteota (*Thermoproteia* sp.) and one Methanobacteriota (*Methanobacteria* sp., with two RNA viruses) (Fig. 5B, fig. S11).

**Fig. 5:**
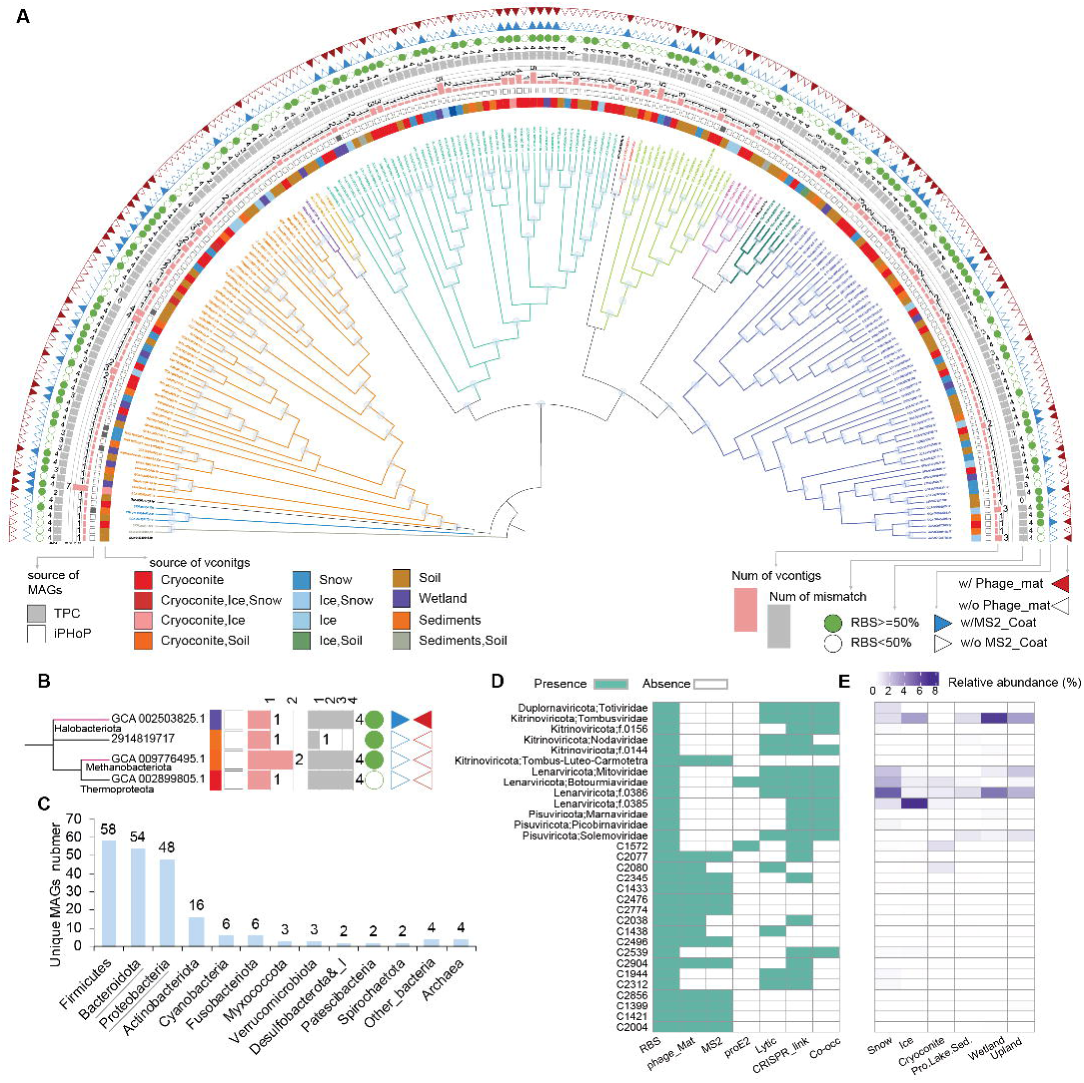
Extended prokaryotic hosts for RNA viruses. (**A**) Phylogenomic tree of Bacteria with CRISPR-spacer linkages to vContigs. Branch colors indicate the phylum (e.g., Cyanobacteria) of MAGs. The tree annotations from the inside ring to the outside are: 1, color pallets show the sources of vContigs (e.g., cryoconite); 2, rectangles show whether the MAGs were from our TPC genome datasets (filled grey) or from iPHoP database (empty); 3, bars show the number of unique vContigs linked to the corresponding MAG; 4, bars show the minimum number of mismatches (including inserted gaps) between vContigs and the corresponding MAGs; 5, dots show whether vContigs with RBS ratio ≥50% (filled green) or not (empty); 6, outwards triangles show whether vContigs coding phage MS2 coat gene (blue filled) or not (empty) and inwards triangles show whether vContigs coding phage maturation gene (blue filled) or not (empty). For MAGs with multiple vContig links, if one of them met the criteria (e.g., carry phage maturation genes), then an annotation was assigned (e.g., filled red triangles). The minimum number of mismatches (including inserted gaps) between vContigs and the corresponding MAGs are shown for all vContigs linked to one MAG. (**B**) Phylogenomic tree of Archaea with CRISPR-spacer linkages to vContigs. Tree annotations are shown as described for panel (A). (**C**) The distribution of Bacteria and Archaea MAGs with CRISPR-spacer linkages to RNA viruses. Taxonomic groups that have been identified to be infected by RNA viruses in previous studies are highlighted with underlines. (**D**) Summary of extended prokaryotic RNA viruses signals for established RNA viral families and unclassified RNA viral clades. (**E**) Heatmap showing the relative abundance of extended prokaryotic RNA viruses. Panel (**E**) shared the same row label as (**D**).

We further evaluated the confidence of the clades containing RNA phages or archaeal RNA viruses in addition to the CRISPR-spacer linkages, and explored other potential clades as prokaryotic RNA viruses by using RNA phage hall marker genes (including phage maturation protein, lytic protein, pro_E2 and pro_ring proteins; fig. S12), RBS motifs, and potential virus-host abundance correlation (table S4, see Materials and Methods for details). Virus-host linkages were deemed credible if they were detected by at least two of the above analyses. We discovered 31 clades (with vContigs number ≥5) assigned to the established families and 13 clades (with vContigs number ≥5) containing RNA viruses of putative new families that could infect prokaryotes (Fig. 5D). Some members of *Tombusviridae* harbor lytic proteins, had CRISPR-spacer linkages, prok_E2/ring proteins, or had RBS ratio ≥50% (Fig. 5D; tables S2 and S4). The lytic proteins and pro_E2/ring proteins are detected in f.0144 vContigs, and 9 vContigs of f.0144 also had an RBS ratio 50% (Fig. 5D).

The relative abundance of extended prokaryotic RNA viral clades is shown in Fig. 5E. Most extended prokaryotic RNA viruses with relative abundance >0.5% belonged to *Lenarviricota*, including *Mitoviridae* in snow (1.63%) and upland (1.55%), *Botourmiaviridae* in snow (2.18%), f.0386 in snow (5.36%), wetland (4.36%), and upland (2.17%), and f.0385 in ice (7.23%). Besides the phylum *Lenarviricota*, only *Tombusviridae* of *Kitrinoviricota* showed a relative abundance >2% in ice (2.77%), wetland (6.76%), and upland (2.76%). While for the unclassified clades, only cluster C1572 (0.81%) and C2080 (0.58%) had a relative abundance >0.5% in cryoconite (Fig. 5E). For the rest of these clades, the mean relative abundances within each habitat were generally less than 0.2% (Fig. 5E).

### AMGs encoded by TPC RNA viruses

Here, we detected 39 AMGs in 15 vContigs (0.2% of the total vContigs; fig. S13 and S14; table S6), which mainly belonged to the functional categories of translation (18), carbohydrate metabolism (4), transportation (5), energy metabolism (5), chaperones (1) and peptidases (1) (fig. S13B). vContigs with AMGs were from diverse taxonomic groups, including 4 vContigs from unclassified groups and 11 from established families, e.g., *Fiersviridae* (3), f.0368.base-Atkins (2), and f.0102.base-Curvula (1). Aside from the vContigs whose hosts were undetermined (9), the remaining RNA viruses carrying AMGs putatively infect prokaryotes (4; Cyanobacteria and undetermined bacteria) and eukaryotes (2; algae, fungi, and undetermined eukaryotes) (fig. S13; table S6).

In addition, these AMG-carrying vContigs were not only detected in the environmental samples of which they were assembled from but also found in widespread environments (Fig. 6A). Among all 15 vContigs with AMGs, 6 (40%) and 6 (40%) of them had a TPM >1,000 and a mean reads coverage >80% in at least 3 samples, respectively (Fig. 6A), indicating high confidence of occurrence of these viruses in widespread environments.

**Fig. 6:**
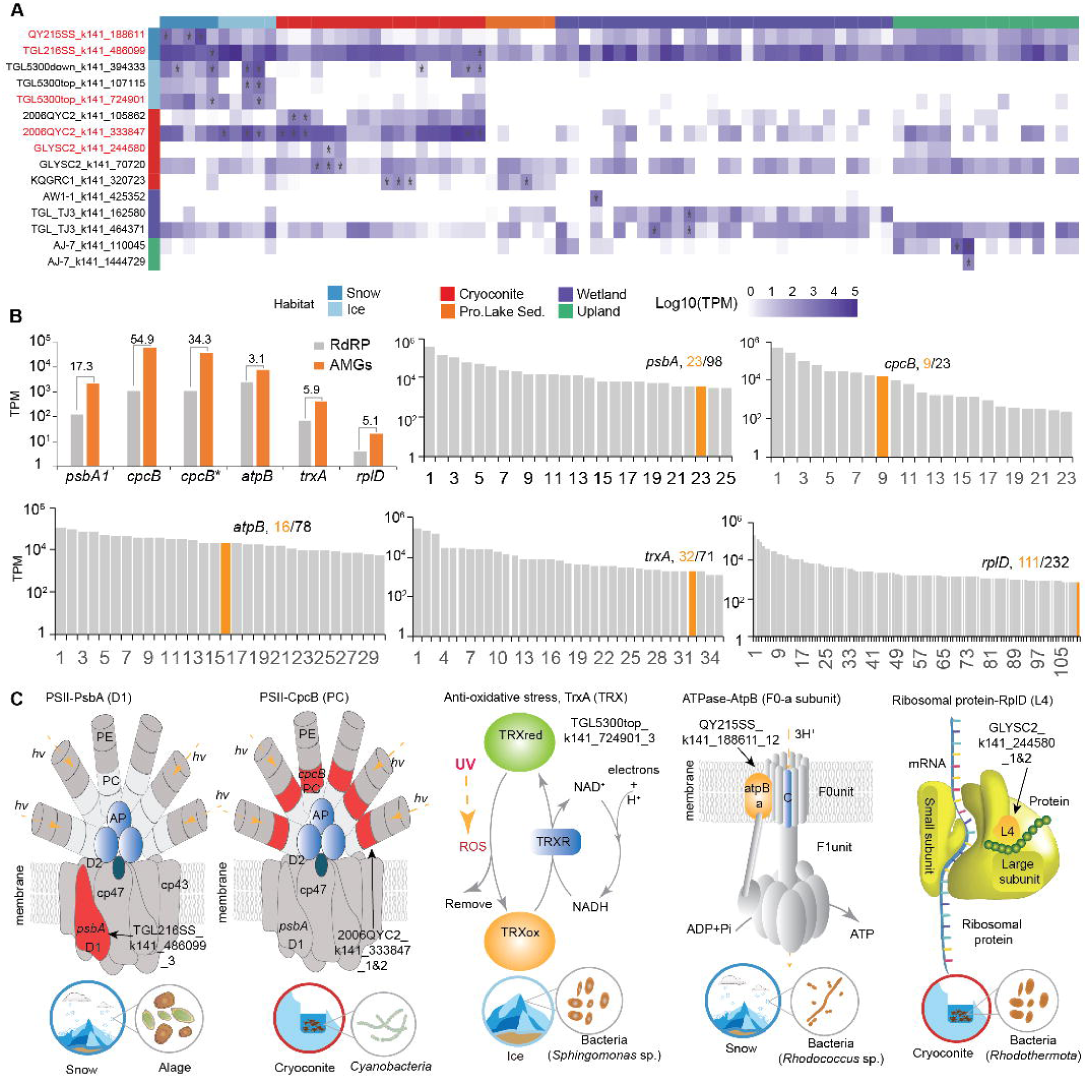
AMGs of the TPC RNA viruses. (**A**) The heatmap showing the TPM and reads coverage of RNA viruses with AMGs across different samples. Row labels are vConitgs names, with those harbouring 5 selected AMGs highlighted in red. (**B**) TPM of RdRps coding genes, AMGs, and comparison of transcripts abundances of AMGs to non-viral encoded functional genes in corresponding samples of selected viral genomes. (**C**) Carton showing the cellular location and function of selected AMGs, and the source and potential host of vContigs with the corresponding AMGs. Asterisks indicate vContigs with horizontal coverage >80% in corresponding samples.

We compared the transcripts abundance of AMGs with the RdRp-coding gene from the same vContig (Fig. 6B; table S6). The abundance of most AMGs (table S6) was markedly lower than RdRp-coding genes. However, the abundance of 5 AMGs from glacier environments were profoundly higher than that of RdRp genes, including the *psbA* (17.3), *cpcB* (54.9 and 34.3 folds higher, respectively), *atpB* (3.1), *trxA* (5.9), and *rplD* (5.1) (Fig. 6B). Moreover, we compared the abundance of the 5 AMGs to the same type of genes encoded by prokaryotes or eukaryotes within the same sample of which the AMGs were retrieved. The abundance of these genes was among the top 9 to 111 among all obtained genes within each sample, indicating a substantial contribution of RNA viral AMGs to the community level gene transcription (Fig. 6B).

AMGs with high abundance included key functional genes for photosynthesis, anti-oxidative stress, energy conservation, and protein synthesis (Fig. 6C). The *psbA* encodes the photosystem II core reaction center protein D1. The *psbA* gene encoded by the snow RNA vContig (TGL216SS_k141_486099) of f.0102.base-Curvula (*Durnavirales*) showed a high similarity (>94%) to and was clustered with the algal (e.g., *Xanthophyceae*) D1 protein in the phylogenetic tree (fig. S15). Therefore, the host of this snow RNA virus is most likely a kind of snow algae. The *cpcB* encodes the phycocyanin protein beta chain, which is a key component of the light-harvesting antenna system in red algae and cyanobacteria. The *cpcB* gene encoded PC protein in the cryoconite RNA vContig (2006QYC2_k141_333847) is identical to that of Cyanobacteria (e.g., WP_193972088.1, unclassified *Microcoleaceae* sp.) (table S7). This vContig belongs to the RNA phage family of *Fiersviridae*, and thus we could speculate that the host of this *cpcB*-harboring vContig is most likely a photoautotrophic cyanobacteria species. The *trxA* encodes the thioredoxin protein, which plays a key role in maintaining intracellular redox status and coping with oxidative stress(*33*). The *trxA* gene encoded thioredoxin protein in the ice RNA vContig (TGL5300top_k141_724901) showed a high similarity to that of *Sphingomonas* sp. (e.g., WP_056058473.1, 97.4%) (table S7). Such contig belongs to the RNA phage family of f.0368.base-Atkins (*Leviviricetes*). Therefore, the host of this *trxA* harboring vContig is most likely a species of heterotrophic *Sphingomonas*. Interestingly, the *atpB* gene encoded ATPase F0a protein in the snow RNA vContig (QY215SS_k141_188611) showed a high similarity to that of bacteria *Rhodococcus antarcticus* sp. (e.g., WP_265383704.1, 90.2%) (table S7). The RNA vContig belongs to f.0386 of *Lenarviricota*. Since f.0386 also posed several lines of RNA phage signals (Fig. 4), we proposed heterotrophic bacteria *Rhodococcus* sp. as the potential host for this vContig. The *rplD* encodes the large subunit ribosomal protein L4, which plays a key role in protein synthesis. The L4 protein coded by the *rplD* gene in the cryoconite RNA vContig (GLYSC2_k141_244580) showed the highest similarity to that of Rhodothermota bacteria species (e.g., *Rubrivirga marina* WP_056058473.1, 86.2%). However, the vContig belongs to unclassified RNA viruses (C1921), therefore, we could only deduce that the putative host is probably a Rhodothermota-associated bacteria (Fig. 6C).

## Discussion

In this study, we obtained the first RNA viral genome dataset for the TPC. Most of those RNA viral species are novel compared to the known RNA virus repertoire, which contributes with >4,900 new species to expand the limits of the known RNA virosphere (Fig. 1). The Shannon diversity indices of TPC RNA viral communities were comparable to those of other environments on Earth(*4, 8, 34*), despite the ecosystems being considered extreme environments. In particular, prokaryotic RNA viruses dominate the TPC RNA virus communities, contrasting to RVMT(*6*) and *Tara* Oceans datasets(*5, 8*), which are dominated by eukaryotic RNA viruses. The genomes of RNA phages account for more than 59% of all classified TPC RNA viruses, which is notably the highest compared to all published global RNA virome datasets(*5, 6, 8*). This could be explained by the availability of hosts, as prokaryotes dominate TPC ecosystems, which is different from the eukaryotic algae-dominated marine ecosystems(*5, 8*). Additionally, we further expanded the range of RNA viruses that could infect prokaryotes. Recent studies have revealed a great diversity of RNA phages(*6, 10, 35–37*). The number of canonical RNA phage species (e.g., *Leviviricetes*) and the taxonomic groups of RNA viruses proposed to be capable of infecting prokaryotic hosts have both substantially increased(*6, 35, 36*). However, the diversity of prokaryotic RNA viruses is still largely unexplored as speculated by Dominguez-Huerta et al(*38*). The results of the TPC RNA viral community composition, structure, and virus-host association well supported this proposition. Our results revealed 31 new viral families and unclassified viral clades that can potentially infect prokaryotic hosts (Fig. 5D). Therefore, this represents a further expansion of RNA phage after Neri et al.(*6*) and Urayama et al.(*35*).

For the host of RNA viruses, we identified 20 new host taxa of prokaryotic RNA viruses, much greater than the previously recognized Bacteroidota(*6*), Chloroflexota(*6*), Protobacteria(*6*), Candidatus *Accumulibacter* sp. SK-02(*9*), and thermoacidophilic bacteria (e.g., *Hydrogenobaculum* sp. of *Aquificota*) from hot springs in Japan(*35*). Most strikingly, we found RNA viruses associated with MAGs of three archaeal phyla, including Halobacteriota, Thermoproteota, and Methanobacteriota (Fig. 5). Till now, no RNA viruses have been identified with a definite association with archaea(*39, 40*), the only domain of life that appeared to be untouched by RNA viruses. This could be the first archaeal RNA virus identified, but further evidence is required for the validation of the archaeal RNA virus proposed here. Therefore, by constructing the largest RNA virus dataset of the TPC, our results substantially expanded the types of prokaryotic RNA viruses and the number of prokaryotes that could serve as hosts for RNA viruses, which highlights their overlooked significant ecological footprints on the food webs and biogeochemical cycling of the TPC environments.

Diverse AMGs were identified in RNA viruses that may enhance the survival of their host in the cryosphere. AMGs with ROS response (TrxA), energy conservation (ATPase), and protein synthesis (ribosomal proteins) were identified in viruses infecting glacier heterotrophic bacteria (Fig. 6C). These AMGs might play a crucial role in removing ROS indirectly induced by high levels of UV radiation(*41, 42*) to avoid cellular damage(*33*), or enhance the metabolism and resistant to the low temperature and nutrient condition on the glacier(*23, 43, 44*). Most excitingly, we found RNA viruses encode two key proteins (PsbA and CpcB) of the core photosynthesis system for the first time. The *psbA* gene codes for the D1 polypeptide of the photosystem II reaction center complex that is found in all photosynthetic organisms(*45, 46*). Viruses carrying this gene have been detected in marine cyanophage (the DNA virus that infects Cyanobacteria), which helps the host cope with radiation damage(*47–49*). In our dataset, the RNA virus with the *psbA* gene was identified from snow and was predicted to infect eukaryotic algae. Similarly, the *cpcB* that encodes a light-harvesting pigment-protein was identified in Cyanobacteria-infecting RNA virus from cryoconite (Fig. 6C). Hence, we propose that the *psbA* and *cpcB* genes encoded by RNA viruses may have a similar function as that encoded by cyanophage(*47–49*), which is to cope with radiation-caused photosynthesis-related protein damage. These genes demonstrated a higher transcription level compared to the cellular genes (Fig. 6b), indicating the active expression of these AMGs. In addition, the hosts of those vContigs with these AMGs were prevalent in global glaciers(*23*), i.e. Xanthophyceae and Chlamydomonadales algae(*50*) dominated in snow, Cyanobacteria predominant in cryoconite(*23, 51*) and potential host (*Sphingomonas* sp.) of *trxA*-coding RNA viruses are prevalent groups in ice(*52, 53*). All these indicated that RNA viruses would facilitate the adaptation and proliferation of their hosts on the glacier surface through the activity of carried AMGs, which may accelerate the melting of glaciers.

The release of viruses with potential public health risks from the cryosphere is growing social concern(*24*) in light of rapid climate change. We revealed 56 RNA viral families associated with eukaryotes in the TPC RNA virome dataset, with *Botourmiaviridae*(*54*), *Solemoviridae*(*55*), and *Benyviridae*(*56*) that might infect plants having high abundance in the glaciers and proglacial lake sediments, respectively. In addition, we also detected *Reoviridae*(*57*) (prevalent in proglacial lake sediments and wetland), *Flaviviridae*(*58*) (wetland), *Hepeviridae*(*59*) (ice), and *Astroviridae*(*60*) (wetland and upland) that might infect animals and vertebrates in various habitats, although with much lower relative abundances (<1.1%) (Fig. 4C). Those families are also frequently detected but with low relative abundance in Arctic permafrost(*18*), grassland soil from the USA(*19, 61*) and Wales(*34*), and various habitats (e.g., Cornfield, Old-growth forest) of terrestrial ecosystems of China(*12*). Hence, those RNA virus families are also widespread in natural environments.

Explicit identification and prediction of pathogenic viruses is still a great challenge(*62*). In order to further identify potential zoonotic RNA viruses in the TPC, we applied Zoonotic_rank(*63*), a machine learning and genome feature(*63*) based approach in addition to the putative host assignment. This analysis showed that only 0.4% of the 8,799 vContigs were categorized as zoonotic viruses with a rank of “very high” (e.g., *Iflaviridae* that infect insects and other arthropods; table S8). However, biological and clinical characterizations are still needed to verify their true infectivity in humans(*62*) and the potential public health risks of TPC RNA virus communities deserve further evaluations. In conclusion, we obtained the first RNA viral genome dataset recovered from the metatranscriptomes of the TPC, which fills a unique geographic and ecosystem gap of global RNA virome atlas. The dataset contains 5,333 vOTUs, from which 92% represent novel species compared to the up-to-date global RNA virome database. We found that diverse RNA phages dominated in major TPC habitats, revealing a large number of previously unidentified viral clades and bacterial hosts associated with diverse metabolisms, challenging the established view of RNA viruses as mostly eukaryotic, and highlighting an overlooked role of RNA viruses in mediating biogeochemical cycling in the cryosphere. Moreover, we revealed diverse AMGs encoded by the RNA viruses that may enhance the survival of hosts in the cryosphere environment. Specifically, the identification of AMGs associated with photosynthesis highlights the roles of RNA viruses in sustaining the growth of hosts on glacier surface and impact glaciers melting. Finally, this study reveals a few of potential pathogenic RNA viruses, which could serve as targets for the RNA virus surveillance with public health risks in the future. Our work provides a wealth of resources for global RNA virosphere study from the cryosphere and greatly expands our understanding of the diversity and ecological roles of environmental RNA viruses.

## Materials and Methods

### Samples collection and preparation

The 79 samples were collected from habitats of glaciers (28 samples), proglacial lake (6), upland soil (16) and wetland soil (29) in permafrost regions from November 2020 to July 2023 (Fig. 1; table S1). Snow (5), ice (5), and cryoconite (18) samples were collected from nine glaciers including Guliya glacier (GLY), Jiemayangzong glacier (JMYZ), Mugagangqiong glacier (MGGQ), Tanggula glacier (TGL), Qiangyong glacier (QY), Kuoqionggangri glacier (KQGR), Parlung No.4 glacier (PL), 24K glacier (24K), Zhuxigou glacier (ZXG). Proglacial lake sediments (6) were collected from Qiangyong Lake that was fed by QY. Upland soils were collected (16, with 13 from surface and 3 from subsurface) from Aerjin (AJ), the Altun Mountains National Nature Reserve and in permafrost regions of the Tuotuohe (TTH), the source region of the Yangtze River(*64*). Wetland soils (29) from the shore of one lake of AJ and permafrost regions of Longxiazai. See Supplementary Materials for details of filed sampling. All samples were stored at –80 °C or liquid nitrogen until total RNA extraction.

### Total RNA extraction, quality checking, sequencing, and reads processing

Total RNA extraction was carried out for samples within half year after sampling. Total RNA was extracted from all samples with the RNAeasy PowerSoil Total RNA Kit (QIAGEN, Hilden, Germany) according to the manufacturer’s instructions. Residual DNA removal and RNA cleaning was carried out using DNase I and the RNA Clean&Concentrator-5 (Zymo, Irvine, USA), respectively. The quantity and quality of total RNA was measured using RNA HS Assay Kit (Thermo Fisher Scientific, Waltham, USA) with Qubit 3 (Invitrogen, Carlsbad, USA) and Bioanalyzer 2100 (Agilent, Santa Clara, USA). Ribosomal RNA (rRNA) was removed with Ribo-off rRNA Depletion Kit V2 kits (Vazyme, Nanjing, China). Library was prepared with VAHTS Universal V6 RNA-seq Library Prep Kit (Vazyme, Nanjing, China) for Illumina following the manufacturer’s recommendations. Sequencing was carried out using the Illumina NovaSeq 6000 platform with pair-end 2×150 mode by Guangdong Magigene Biotechnology Co. Ltd. (Guangzhou, China) company.

In total, over 5 TB raw reads were obtained, with an average of 313.8 million reads per sample. Raw reads were trimmed with Trimmomatic v0.39(*65*) with default parameters. The remaining rRNA reads were removed from total RNA with SortMeRNA v4.3.6(*66*), which yielded 1.5 TB non-rRNA reads, with an average of 108.2 million mRNA reads per sample.

### Metatranscriptomic reads assembly, RNA virus identification and verification

mRNA reads were *de novo* assembled into contigs using MEGAHIT v1.29(*67*) with default parameters as previously described(*5*). All RNA viruses of the kingdom *Orthornavirae* [hence, excluding satellite RNAs (realm *Ribozyviria*) and reverse-transcribing RNA viruses (kingdom *Pararnavirae*)] share a single hallmark protein, the RNA-dependent RNA polymerase (RdRp)(*28*). Therefore, we searched for RNA viral genomes through mining assembled contigs encoding RdRps. Contigs with length ≥1 kb were kept for gene prediction with Prodigal v2.6.3(*68*) with the “-p meta” option setting and the default translation table. The yield protein sequences were subject to RdRp domain identification similar to previous studies by using hidden Markov model (HMMs) and hmmsearch (HMMER v3.3.2)(*69*) approaches. To increase the detection of divergent target domain sequences, HMMs from the *Tara* Oceans RNA virus dataset(*5, 8*), NeoRdRp pipeline(*70*), and the RVMT global RNA virome dataset(*6*) were used for RdRp domain search. The protein sequence containing domains with an E-value ≤0.01 was kept for RdRp verification through the Palmscan algorithm(*31*)and metagenomic reads mapping strategy as described by Hou et al.(*4, 6*) to remove DNA remnants (table S9). See Supplementary Materials for more details on RdRp verification. In total, we obtained 8,799 RNA viral genomes (vContigs) from all samples, with an average of ∼111 vContigs per sample.

### vContigs quality assessment

The completeness of the vContigs was assessed by using CheckV v1.01(*71*) with default parameters. To evaluate the RdRp domain completeness, vContigs were translated to amino acid (AA) sequences by using transeq in EMBOSS v6.5.7.0 with six possible frames and all translational codes with transeq to resolve difficulties associated with alternative genetic code usage and non-canonical translation events. All translated AA sequences were searched against the 65 RdRp HMM profiles from Wolf et al.(*16*). The AA sequences containing RdRp domains with an E-value ≤ 0.01 were kept. The aligning regions of each AA sequence with the highest score and longest length were extracted as the best RdRp domain. The frame and code combinations from the AA sequence with the best RdRp domain were kept for genome annotation of the corresponding RNA virus (see below). The AA sequences with aligning regions of ≥90% of the average length of the best-matching HMMs were considered to contain “complete” RdRp domains.

### The generation of viral operational taxonomic units (vOTUs) of RNA viruses

Generally, virus operational taxonomic units (vOTUs) represent the “species” rank of RNA viruses (*5, 6, 8*). In this study, 8,799 RdRp domain sequences were clustered into 5,712 vOTUs using CD-HIT v4.8.1(*72*), with a clustering threshold of 90% amino acid identity (AAI) and default parameters as previously described(*4–6, 8*). In addition, 8,799 vContigs nucleotide sequences were clustered into the 6,620 clusters (C90) by using CD-HIT v4.8.1, with a clustering threshold of 90% average nucleotide identity (ANI) and 80% alignment fraction (AF)(*5, 6, 8*).

### Taxonomy classification of TPC RNA viruses

To assign taxonomic information for RNA viruses, we applied pipelines including Markov Cluster Algorithm (MCL) clustering and network analysis as described by Zayed et al.(*5*), RdRp AAI and vContigs ANI comparison as described by Neri et al(*6*)., and RdRps phylogenetic tree construction(*5, 6*). Public reference databases, including the global RNA virome dataset (RVMT)(*6*), the *Tara* Oceans datasets (TO)(*5*), the Serratus palmprint database (Palmdb)(*31*) and IMG/VR4 database(*73*) were used along with the pipelines for taxonomy classification of TPC RNA viruses. We constructed phylogenetic trees for all RdRp AA sequences classified into 6 established RNA viral phyla. Reference sequences were obtained from RVMT C90, TO, IMG/VR-V4 during the classification step (see above). In brief, RdRp AA sequences from each dataset or database that affiliated or clustered with our RdRps at species or family ranks were kept as references. See Supplementary Materials for details on taxonomy classification.

### Novelty assessment of TPC RNA viruses

To assess the novelty of the TPC RNA viruses, we compared the RNA viral genomes and RdRp domains of this study with the ones of known RNA viral genome and four reference databases, including the TO(*8*), the RVMT(*6*), the LucaProt(*4*), and the Serratus(*7*). The novelty was estimated through two approaches:

i. RdRp based. To evaluate the novelty at phylum/class level, “complete” RdRps from TPC, RVMT and TO were clustered by using USEARCH v10.0.240 (-id 0.5, -qcov 0.5) and MCL clustering (inflation values of 1.1). Meanwhile, to evaluate the novelty at family and species level, we clustered our RdRps with the ones from RVMT, LucaProt, TO and Serratus database by using USEARCH v10.0.240 with UCLUST algorithm as described above. RdRps AAIs of 50% and 90% were considered as convenient similarity cutoff for RNA viruses “family-genus” and “species” ranks, respectively.
ii. RNA virus genome based. vContigs from the TPC also were clustered with the vContigs from RVMT and TO database by using USEARCH v10.0.240 with parameters: -id 0.9 and -qcov 0.5.

### Calculation of the relative abundance and diversity of vOTUs

mRNA reads were mapped to the nucleotide sequences of vOTUs genome with Bowtie2 v2.2.5 using the sensitive mode. The mapped reads were further filtered using Samtools v1.15.1 with -q 30 option to keep only high-quality mappings. TPM were used to represent the relative abundance of the vOTUs. The TPM value of each vOTU within each sample was calculated by using the quality filtered mapping file (sorted bam file) and CoverM v0.6.1 with -m tpm option (https://github.com/wwood/CoverM). The coverage percentage of each vOTU was estimated by using CoverM v0.6.1 with -m covered_fraction option.

Alpha-diversity indices of each sample including Shannon and Richness were calculated using vegan package in R v4.0.3 with the TPM data matrix. The dissimilarity of RNA viral communities (beta-diversity) was estimated by vegan package in R v4.0.3 with Bray-Curtis distance calculated from the TPM matrix. The clustering of viral communities was visualized with non-metric multi-dimensional scaling (NMDS) with the TPM-transformed Bray-Curtis distance.

### Enrichment analysis of RNA virus family and vOTUs

Enriched RNA virus families were identified by their relative abundance difference among habitats. If the maximum relative abundance of a family in a habitat was four times that of the second largest relative abundance, we considered that family to be enriched.

### Metagenome quality control and mapping to the viral genome

The metagenomic sequences mapping approach was used to verify that the identified RdRps were from RNA viruses rather than from the DNA of cellular organisms. Thirty-two metagenomic samples that accompanied with the metatranscriptomes from the same sample (table S9) were used. The sampling methods of metagenome samples were similar as that of the metatranscriptome samples (see above). In addition, for metatranscriptome without accompanied metagenome, sequences from the metagenome of similar samples were used (table S9). Metagenome reads were mapped against the RdRp nucleotide sequences from Prodigal with Bowtie2 v2.2.5 with the “end-to-end” setting to check whether there was a DNA counterpart. The mapped reads were further filtered using Samtools v1.15.1 with parament: -q 30 -F 0×08 -b -f 0×2. RdRps with horizontal metagenome reads coverage ≥75% and average vertical coverage depths ≥1× were removed.

### The transcript abundances of MAGs and functional genes

We collected 15,640 medium-high quality MAGs (with completeness >50 and contamination <10) from the glacier, lake sediment, wetland, and upland of the TP cryosphere(*23*). All these MAGs were clustered into strain level units via dRep v3.4.5 with ANI 99% threshold. A total of 6,063 strain-level MAGs were obtained and used to estimate transcript abundances. BBmap v39.01 and coverM with ‘genome’ module with default parameters were used to calculate the transcript abundance of MAGs. Briefly, clean mRNA reads were mapped to host MAG sequences with default parameters, and yielded bam files were sorted using Samtools v1.15.1. TPM was generated based on the sorted bam files using coverM with ‘--min-covered-fraction 10 -m tpm’ parameters.

To find the host marker genes (including both prokaryotic and eukaryotic *cox1* and *rps3*), we used an HMM profile from Pfam Cox1 (PF00115), and HMMs for (https://github.com/AJProbst/rpS3_trckr) rps3 and searched using hmmsearch (-E 0.00001) from the HMMER suite as previously described(*19*). The Cox1 protein sequences were dereplicated and clustered at 98% AAI representing an estimated species-level designation. The Rps3 protein sequences were clustered at 99% AAI, representing species-level differences using CD-HIT. The identified proteins were classified using the NCBI protein blast. Clean mRNA reads were mapped to host the nucleotide sequences of representing *rps3* and *cox1* with default parameters, and yielded bam files were sorted using Samtools v1.15.1. TPM were generated based on the sorted bam files as described above (fig. S16).

### Inference of virus-host relationship and risk assessment of RNA viruses

To infer the putative host of RNA viruses, we used four approaches:

i. Taxonomy based. The known RNA virus taxa (phyla, classes, and families) that were assigned to the vOTUs after clustering the RdRp domain protein sequence similarity network (see RdRp-based taxonomy annotation of RNA viruses) were used to retrieve information on putative hosts.
ii. CRISPR-spacer screening. We retrieved 52,134 CRISPR spacer sequences from 15,640 medium-high quality MAGs of the TPC. Identification of CRISPR spacer was performed by MinCED v0.4.2 with default parameters. In addition, a CRISPR-Cas spacer database of iPHoP_db_Sept21_rw compiled from GTDB, IMG, and GEM v1 database with 1,398,130 CRISPR-Cas spacers was also included for virus-host prediction. All spacers were queried for matches against viral contigs using the BLASTn-short function implemented in the NCBI BLAST v2.13 package with parameters “-evalue 1e-10, -perc_identity 95, -dust no -word_size 7”, allowing only <=4 mismatches (≥86% identity, table S5) across the entire length of the spacer(*4, 6*).
iii. EVE based. EVEs are nucleotide sequences derived from DNA or RNA viruses that have integrated into host genomes. For this third approach, all nucleotide sequences from cellular organisms available in NCBI GenBank release 243 were used as a nucleotide database. To avoid including exogenous RNA virus genomes in the database, we excluded sequences shorter than 45 kb since the longest RNA virus genome reported so far is 45 kb(*4*). To assess the evolutionary relationship of OTUs of exogenous viruses to EVEs, the near-complete RdRp protein sequences were searched against the nucleotide database by using tblastn algorithm. As previously described(*74*), the thresholds were set to 100 amino acids for alignment length and 1 × 10^−20^ for e-value.
iv. Co-occurrence network based on abundance(*19*). Co-occurrence network analysis was used to reveal putative virus-host linkages. First, vConitgs, MAGs, *rps3* and *cox1* gene were filtered based on the threshold: with a max TPM>1000 and detected in at least 10 samples with TPM>5. Then a co-occurrence network was built based on the abundance of filtered sequences with Spearman correlation algorithms. Only correlations with thresholds: r>0.7 and p <0.001 were kept as previously described(*19*) (table S4). Retaliative abundance datasets were deposited in figshare (https://figshare.com/s/949e8151580d648951a5).

### Zoonotic risks prediction

In addition, zoonotic_rank(*63*) (https://github.com/Nardus/zoonotic_rank.git) with default parameters was applied to identify and prioritize potential human-infecting viruses in the TPC RNA virus communities. Metadata files were generated for all CDS regions from 8,799 vContigs using Prodigal, followed by the execution of the PredictNovel.R script. A total of 1,043 vContigs met the default calibrated_score threshold of 0.293, which were classified as potential zoonotic viruses with priority_category ranks of “high” (1,007, 11.4% of 8,799) and “very high” (36, 0.4%). To control the false positive rate, we retained only those vContigs with a priority_category rank of “very high” as putative zoonotic viruses (table S8).

### Genomic features and function annotation of TPC RNA viruses

Functional annotation of the TPC RNA virus was combined with three approaches:

i. ORFs based on default code. A total 8,799 nucleotide sequences of vContigs were annotated by InterProScan v5.62-94.0(*75*) against MobiDBLite (v2.0)(*76*), PRINTS (v.42.0)(*77*), Phobius (v.1.0.1)(*78*) and TMHMM (v.2.0c) database(*79*).
ii. ORFs based on optimized code. Optimize code was determined by the longest RdRp and corresponding code table as described in the vContigs quality assessment section. For RNA viral genome domain annotation, ORFs were identified by using Prodigal v2.6.3 with the optimized genetic code. One iteration of hhblits (-n 1 -e 0.001) by HHsuite v3.3.0(*80*) and UniRef30_2020_03 database(*81*) was used to generate profiles for amino-acid sequences. Generated profiles were searched against Pfam 35 to identify domains in the amino-acid sequences, and hits with >95% probability score were used for domain annotation(*5*).
iii. Based on amino acid sequences of functional domains. Since the determination of code usage is still a challenge, we also applied a functional domain annotation on the translated virus genome sequences with the longest RdRp domain detected without ORF calling. AA sequences from contigs translated by EMBOSS transeq of each vConitgs were annotated by hmmsearch (from the HMMER V3.3.2 suite)(*82*) to match these proteins to hidden Markov models (HMMs) gathered from multiple protein profile databases using a maximal E-value of 0.001 (PFam 35(*83*), CDD v.3.19(*84*), Gene3D v4.3(*85*), ECOD 2017 release(*86*), LysDB(*6*)). LysDB is from a previous study(*6*), which contains a custom collection of HMMs profiles for proteins with bacteriolytic functions.

### Identification of AMGs in vContigs

Auxiliary metabolic genes (AMGs) encoded by vContigs were mainly identified using the functional annotation and Manual checking as previously described(*8*). All annotations from RNA viruses contigs annotation were screened for putative cellular metabolic functions. To verify the AMGs encoded by RNA viruses were not from transcripts of cellular genes, we applied a similar strategy as RdRps through metagenome reads mapping against the nucleotide sequences of putative AMGs. Metagenomic clean reads were mapped against the nucleotide sequences of AMGs with Bowtie2 v2.2.5 with the “end-to-end” setting. The mapped reads were further filtered using Samtools v1.15.1 with parameter: -q 30 -F 0×08 -b -f 0×2. AMGs with horizontal coverage greater than 75% were considered as “contamination” from host transcripts and were removed from downstream analysis. Thirty-nine AMGs from 15 vContigs were kept (Fig. 6a; Table 6). In addition, the functions of these AMGs were further checked with Phyre^2^ as previously described(*87*). For genomic architecture, the annotation files of vContigs with putative AMGs were visualized in Geneious v2021.2.2 (fig. S14). In addition, a phylogenetic analysis of PsbA amino acid sequences was carried out to check its origins (fig. S15). See Supplementary Materials for more details.

### Functional annotation of MAGs with RNA virus associations

All 208 MAGs with RNA viruses association were annotated using the Distilled and Refined Annotation of Metabolism (DRAM)(*88*) pipeline with default parameter as previously described(*89*). DRAM is a tool for annotating genomes, and KOfam, UniRef-90, PFAM-A, dbCAN, RefSeq viral, VOGDB, and the MEROPS peptidase databases were used for homolog search and genome annotations in the current study (fig. S17).

### Identification of Shine-Dalgarno (SD) sequences of RNA viruses

Two pipelines were used to identify the ribosome binding site (RBS) motif (Shine-Dalgarno sequences, SD) within each vContigs (fig. S18), including OSTIR(*90*) and Prodigal(*68*). The Shine-Dalgarno (SD) sequence is an RBS in bacterial and archaeal mRNA. The prevalence of SD in an RNA virus indicates it is more likely a prokaryotic RNA virus(*6, 91, 92*). OSTIR v1.1.0 was used to identify SD sequences of the virus with anti-Shine-Dalgarno sequences (aSD: 5’-ACCTCCTTT/A-3’). For Prodigal, default parameters were used. In addition, vContigs from the RVMT and TO were also included for SD sequence identification.

### Statistics analysis and figure generation

Statistics analyses like PERMANOVA were conducted in R using the ‘vegan’ package (v.2.5-7). Significant difference evaluation (e.g., DunnTest *post-hoc* test) was also performed using ‘FSA’ package (v0.9.5; https://fishr-core-team.github.io/FSA/). Manuscript figures were mainly generated in R sing ggplot2 and pheatmap (https://github.com/raivokolde/pheatmap) packages and custom R scripts. The figure format was adjusted using Adobe Illustrator if needed.

## Supporting information

Supplementary Materialss

## Acknowledgments

We thank Yang Liu, Hongfei Chi and Wenqiang Wang for help on field sampling, Xuefeng Zhang for help on data analysis, and Dr. Kevin Xu Zhong and Qicheng Bei for help on RNA extraction procedure setup. We also thank “Chayu integrated observation and research station of the Tibet Autonomous Region for the changes in multisphere processes of the monsoon passage” for supporting this work.

## Author contributions

YQ.L., PF.L., TD.Y., and NZ.J. initiated the study; YQ.L. and PF.L. designed the study; R.W. and ML.F. carried out sample processes and total RNA extraction. ZH.Z. and PF.L. performed all the data analysis, including metagenomes, metatranscriptomes and RNA viral community analysis, with results interpretations from also G.D-H, R.Z., HN.W., YG.Z. PF.L., YQ.L. and ZH.Z. led the manuscript writing with the help from R.Z, G.D-H, HN.W, MK.J., and Q.Z. All authors contributed to the editing of the text and approved the final version.

## Data and materials availability

Metatranscriptome datasets from current study were deposited at NCBI with accession number of PRJNA1105542. TPC RNA virus genome datasets, including the RdRps amino acid sequences, vContigs sequences, full functional annotations of the 8,799 RNA virus genomes were available at figshare (https://figshare.com/s/949e8151580d648951a5). The Codes and programs used for analyses are described in Materials and Methods, and are also available at https://github.com/Hame1N/TPC-RNA-virus.

## References

1. E. V. Koonin, J. H. Kuhn, V. V. Dolja, M. Krupovic, Megataxonomy and global ecology of the virosphere. ISME J 18, wrad042 (2024).

2. E. V. Koonin, M. Krupovic, V. I. Agol, The Baltimore classification of viruses 50 years later: How does It stand in the light of virus evolution? Microbiol. Mol. Biol. Rev. 85, e00053–00021 (2021).

3. Y.-F. Pan, H. Zhao, Q.-Y. Gou, P.-B. Shi, J.-H. Tian, Y. Feng, K. Li, W.-H. Yang, D. Wu, G. Tang, B. Zhang, Z. Ren, S. Peng, G.-Y. Luo, S.-J. Le, G.-Y. Xin, J. Wang, X. Hou, M.-W. Peng, J.-B. Kong, X.-X. Chen, C.-H. Yang, S.-Q. Mei, Y.-Q. Liao, J.-X. Cheng, J. Wang, Chaolemen, Y.-H. Wu, J.-B. Wang, T. An, X. Huang, J.-S. Eden, J. Li, D. Guo, G. Liang, X. Jin, E. C. Holmes, B. Li, D. Wang, J. Li, W.-C. Wu, M. Shi, Metagenomic analysis of individual mosquito viromes reveals the geographical patterns and drivers of viral diversity. *Nat*. Ecol. Evol. 8, 947–959 (2024).

4. X. Hou, Y. He, P. Fang, S.-Q. Mei, Z. Xu, W.-C. Wu, J.-H. Tian, S. Zhang, Z.-Y. Zeng, Q.-Y. Gou, G.-Y. Xin, S.-J. Le, Y.-Y. Xia, Y.-L. Zhou, F.-M. Hui, Y.-F. Pan, J.-S. Eden, Z.-H. Yang, C. Han, Y.-L. Shu, D. Guo, J. Li, E. C. Holmes, Z.-R. Li, M. Shi, Artificial intelligence redefines RNA virus discovery. bioRxiv, 2023.2004.2018.537342 (2023).

5. A. A. Zayed, J. M. Wainaina, G. Dominguez-Huerta, E. Pelletier, J. Guo, M. Mohssen, F. Tian, A. A. Pratama, B. Bolduc, O. Zablocki, D. Cronin, L. Solden, E. Delage, A. Alberti, J.-M. Aury, Q. Carradec, C. d. Silva, K. Labadie, J. Poulain, H.-J. Ruscheweyh, G. Salazar, E. Shatoff, R. Bundschuh, K. Fredrick, L. S. Kubatko, S. Chaffron, A. I. Culley, S. Sunagawa, J. H. Kuhn, P. Wincker, M. B. Sullivan, S. G. Acinas, M. Babin, P. Bork, E. Boss, C. Bowler, G. Cochrane, C. d. Vargas, G. Gorsky, L. Guidi, N. Grimsley, P. Hingamp, D. Iudicone, O. Jaillon, S. Kandels, L. Karp-Boss, E. Karsenti, F. Not, H. Ogata, N. Poulton, S. Pesant, C. Sardet, S. Speich, L. Stemmann, M. B. Sullivan, S. Sungawa, P. Wincker, Cryptic and abundant marine viruses at the evolutionary origins of Earth’s RNA virome. Science 376, 156–162 (2022).

6. U. Neri, Y. I. Wolf, S. Roux, A. P. Camargo, B. Lee, D. Kazlauskas, I. M. Chen, N. Ivanova, L. Zeigler Allen, D. Paez-Espino, D. A. Bryant, D. Bhaya, A. B. Narrowe, A. J. Probst, A. Sczyrba, A. Kohler, A. Séguin, A. Shade, B. J. Campbell, B. D. Lindahl, B. K. Reese, B. M. Roque, C. DeRito, C. Averill, D. Cullen, D. A. C. Beck, D. A. Walsh, D. M. Ward, D. Wu, E. Eloe-Fadrosh, E. L. Brodie, E. B. Young, E. A. Lilleskov, F. J. Castillo, F. M. Martin, G. R. LeCleir, G. T. Attwood, H. Cadillo-Quiroz, H. M. Simon, I. Hewson, I. V. Grigoriev, J. M. Tiedje, J. K. Jansson, J. Lee, J. S. VanderGheynst, J. Dangl, J. S. Bowman, J. L. Blanchard, J. L. Bowen, J. Xu, J. F. Banfield, J. W. Deming, J. E. Kostka, J. M. Gladden, J. Z. Rapp, J. Sharpe, K. D. McMahon, K. K. Treseder, K. D. Bidle, K. C. Wrighton, K. Thamatrakoln, K. Nusslein, L. K. Meredith, L. Ramirez, M. Buee, M. Huntemann, M. G. Kalyuzhnaya, M. P. Waldrop, M. B. Sullivan, M. O. Schrenk, M. Hess, M. A. Vega, M. A. O’Malley, M. Medina, N. E. Gilbert, N. Delherbe, O. U. Mason, P. Dijkstra, P. F. Chuckran, P. Baldrian, P. Constant, R. Stepanauskas, R. A. Daly, R. Lamendella, R. J. Gruninger, R. M. McKay, S. Hylander, S. L. Lebeis, S. P. Esser, S. G. Acinas, S. S. Wilhelm, S. W. Singer, S. S. Tringe, T. Woyke, T. B. K. Reddy, T. H. Bell, T. Mock, T. McAllister, V. Thiel, V. J. Denef, W.-T. Liu, W. Martens-Habbena, X.-J. Allen Liu, Z. S. Cooper, Z. Wang, M. Krupovic, V. V. Dolja, N. C. Kyrpides, E. V. Koonin, U. Gophna, Expansion of the global RNA virome reveals diverse clades of bacteriophages. Cell 185, 4023–4037 (2022).

7. R. C. Edgar, J. Taylor, V. Lin, T. Altman, P. Barbera, D. Meleshko, D. Lohr, G. Novakovsky, B. Buchfink, B. Al-Shayeb, J. F. Banfield, M. de la Peña, A. Korobeynikov, R. Chikhi, A. Babaian, Petabase-scale sequence alignment catalyses viral discovery. Nature 602, 142–147 (2022).

8. G. Dominguez-Huerta, A. A. Zayed, J. M. Wainaina, J. Guo, F. Tian, A. A. Pratama, B. Bolduc, M. Mohssen, O. Zablocki, E. Pelletier, E. Delage, A. Alberti, J.-M. Aury, Q. Carradec, C. d. Silva, K. Labadie, J. Poulain, C. Bowler, D. Eveillard, L. Guidi, E. Karsenti, J. H. Kuhn, H. Ogata, P. Wincker, A. Culley, S. Chaffron, M. B. Sullivan, Diversity and ecological footprint of Global Ocean RNA viruses. Science 376, 1202–1208 (2022).

9. Y. I. Wolf, S. Silas, Y. Wang, S. Wu, M. Bocek, D. Kazlauskas, M. Krupovic, A. Fire, V. V. Dolja, E. V. Koonin, Doubling of the known set of RNA viruses by metagenomic analysis of an aquatic virome. Nat. Microbiol. 5, 1262–1270 (2020).

10. J. Callanan, S. R. Stockdale, A. Shkoporov, L. A. Draper, R. P. Ross, C. Hill, Expansion of known ssRNA phage genomes: From tens to over a thousand. Sci. Adv. 6, eaay5981 (2020).

11. L. Wu, Y. Liu, W. Shi, T. Chang, P. Liu, K. Liu, Y. He, Z. Li, M. Shi, N. Jiao, X. Dong, Q. Zheng, Alpine lakes harbor diverse, endemic and structurally unique RNA viruses. bioRxiv, 2024.2007.2004.601995 (2024).

12. Y.-M. Chen, S. Sadiq, J.-H. Tian, X. Chen, X.-D. Lin, J.-J. Shen, H. Chen, Z.-Y. Hao, M. Wille, Z.-C. Zhou, J. Wu, F. Li, H.-W. Wang, W.-D. Yang, Q.-Y. Xu, W. Wang, W.-H. Gao, E. C. Holmes, Y.-Z. Zhang, RNA viromes from terrestrial sites across China expand environmental viral diversity. Nat. Microbiol. 7, 1312–1323 (2022).

13. S. R. Krishnamurthy, A. B. Janowski, G. Zhao, D. Barouch, D. Wang, Hyperexpansion of RNA bacteriophage diversity. PLoS Biol. 14, e1002409 (2016).

14. J. K. Jansson, R. Wu, Soil viral diversity, ecology and climate change. Nat. Rev. Microbiol. 21, 296–311 (2022).

15. M. Liao, Y. Xie, M. Shi, J. Cui, Over two decades of research on the marine RNA virosphere. iMeta 1, e59 (2022).

16. Y. I. Wolf, D. Kazlauskas, J. Iranzo, A. Lucía-Sanz, J. H. Kuhn, M. Krupovic, V. V. Dolja, E. V. Koonin, Origins and Evolution of the Global RNA Virome. mBio 9, e02329–02318 (2018).

17. J. M. Labonté, K. L. Campbell, There’s more to RNA viruses than diseases. Science 376, 138–139 (2022).

18. R. Wu, E. M. Bottos, V. G. Danna, J. C. Stegen, J. K. Jansson, M. R. Davison, RNA viruses linked to eukaryotic hosts in thawed permafrost. mSystems 7, e00582–00522 (2022).

19. E. P. Starr, E. E. Nuccio, J. Pett-Ridge, J. F. Banfield, M. K. Firestone, Metatranscriptomic reconstruction reveals RNA viruses with the potential to shape carbon cycling in soil. Proc. Natl. Acad. Sci. U. S. A 116, 25900–25908 (2019).

20. Guillermo Dominguez-Huerta, J. M. Wainaina, A. A. Zayed, A. I. Culley, J. H. Kuhn, M. B. Sullivan, The RNA virosphere: How big and diverse is it? Environ. Microbiol. 25, 209–215 (2023).

21. M. Vila-Nistal, L. Maestre-Carballa, F. Martinez-Hernández, M. Martinez-Garcia, Novel RNA viruses from the Atlantic Ocean: Ecogenomics, biogeography, and total virioplankton mass contribution from surface to the deep ocean. Environ. Microbiol. 25, 3151–3160 (2023).

22. D. Qin, Y. Ding, C. Xiao, S. Kang, J. Ren, J. Yang, S. Zhang, Cryospheric science: research framework and disciplinary system. Natl. Sci. Rev. 5, 255–268 (2018).

23. Y. Liu, M. Ji, T. Yu, J. Zaugg, A. M. Anesio, Z. Zhang, S. Hu, P. Hugenholtz, K. Liu, P. Liu, Y. Chen, Y. Luo, T. Yao, A genome and gene catalog of glacier microbiomes. Nat. Biotechnol. 40, 1341–1348 (2022).

24. R. Wu, G. Trubl, N. Taş, J. K. Jansson, Permafrost as a potential pathogen reservoir. One Earth 5, 351–360 (2022).

25. A. El-Sayed, M. Kamel, Future threat from the past. Environ. Sci. Pollut. Res. Int. 28, 1287–1291 (2021).

26. T. Yao, T. Bolch, D. Chen, J. Gao, W. Immerzeel, S. Piao, F. Su, L. Thompson, Y. Wada, L. Wang, T. Wang, G. Wu, B. Xu, W. Yang, G. Zhang, P. Zhao, The imbalance of the Asian water tower. Nat. Rev. Earth. Environ. 3, 618–632 (2022).

27. B. C. Canavan, Opening Pandora’s Box at the roof of the world: Landscape, climate and avian influenza (H5N1). Acta Trop. 196, 93–101 (2019).

28. E. V. Koonin, V. V. Dolja, M. Krupovic, A. Varsani, Y. I. Wolf, N. Yutin, F. M. Zerbini, J. H. Kuhn, Global organization and proposed megataxonomy of the virus world. Microbiol. Mol. Biol. Rev. 84, e00061–00019 (2020).

29. S. Venkataraman, B. V. L. S. Prasad, R. Selvarajan, RNA Dependent RNA Polymerases: Insights from Structure, Function and Evolution. Viruses 10, 76 (2018).

30. J. Charon, J. P. Buchmann, S. Sadiq, E. C. Holmes, RdRp-scan: A bioinformatic resource to identify and annotate divergent RNA viruses in metagenomic sequence data. Virus Evol. 8, veac082 (2022).

31. A. Babaian, R. Edgar, Ribovirus classification by a polymerase barcode sequence. PeerJ 10, e14055 (2022).

32. S. Roux, A. P. Camargo, F. H. Coutinho, S. M. Dabdoub, B. E. Dutilh, S. Nayfach, A. Tritt, iPHoP: An integrated machine learning framework to maximize host prediction for metagenome-derived viruses of archaea and bacteria. PLoS Biol. 21, e3002083 (2023).

33. B. Ezraty, A. Gennaris, F. Barras, J.-F. Collet, Oxidative stress, protein damage and repair in bacteria. Nat. Rev. Microbiol. 15, 385–396 (2017).

34. L. S. Hillary, E. M. Adriaenssens, D. L. Jones, J. E. McDonald, RNA-viromics reveals diverse communities of soil RNA viruses with the potential to affect grassland ecosystems across multiple trophic levels. ISME Commun. 2, 34 (2022).

35. S.-i. Urayama, A. Fukudome, M. Hirai, T. Okumura, Y. Nishimura, Y. Takaki, N. Kurosawa, E. V. Koonin, M. Krupovic, T. Nunoura, Double-stranded RNA sequencing reveals distinct riboviruses associated with thermoacidophilic bacteria from hot springs in Japan. Nat. Microbiol. 9, 514–523 (2024).

36. S. Sadiq, E. C. Holmes, J. E. Mahar, Genomic and phylogenetic features of the *Picobirnaviridae* suggest microbial rather than animal hosts. bioRxiv, 2024.2002.2004.578841 (2024).

37. T. Gan, D. Wang, Picobirnaviruses encode proteins that are functional bacterial lysins. Proc. Natl. Acad. Sci. U. S. A 120, e2309647120 (2023).

38. G. Dominguez-Huerta, J. M. Wainaina, A. A. Zayed, A. I. Culley, J. H. Kuhn, M. B. Sullivan, The RNA virosphere: How big and diverse is it? Environ. Microbiol. 25, 209–215 (2023).

39. C. Le Lay, J. N. Hamm, T. J. Williams, M. Shi, R. Cavicchioli, E. C. Holmes, Viral community composition of hypersaline lakes. Virus Evol. 9, vead057 (2023).

40. B. Bolduc, D. P. Shaughnessy, Y. I. Wolf, E. V. Koonin, F. F. Roberto, M. Young, Identification of novel positive-strand RNA viruses by metagenomic analysis of archaea-dominated Yellowstone hot springs. J. Virol. 86, 5562–5573 (2012).

41. A. Latifi, M. Ruiz, C.-C. Zhang, Oxidative stress in cyanobacteria. FEMS Microbiol. Rev. 33, 258–278 (2009).

42. S. Xue, Y. Zang, J. Chen, S. Shang, L. Gao, X. Tang, Ultraviolet-B radiation stress triggers reactive oxygen species and regulates the antioxidant defense and photosynthesis systems of intertidal red algae Neoporphyra haitanensis. Front. Mar. Sci. 9, 1043462 (2022).

43. A. Boetius, A. M. Anesio, J. W. Deming, J. A. Mikucki, J. Z. Rapp, Microbial ecology of the cryosphere: sea ice and glacial habitats. Nat. Rev. Microbiol. 13, 677–690 (2015).

44. M. Bourquin, S. B. Busi, S. Fodelianakis, H. Peter, A. Washburne, T. J. Kohler, L. Ezzat, G. Michoud, P. Wilmes, T. J. Battin, The microbiome of cryospheric ecosystems. Nat. Commun. 13, 3087 (2022).

45. A. Li, T. You, X. Pang, Y. Wang, L. Tian, X. Li, Z. Liu, Structural basis for an early stage of the photosystem II repair cycle in Chlamydomonas reinhardtii. Nat. Commun. 15, 5211 (2024).

46. D. J. Vinyard, G. M. Ananyev, G. Charles Dismukes, Photosystem II: The reaction center of oxygenic photosynthesis. Annu. Rev. Biochem. 82, 577–606 (2013).

47. E. T. Sieradzki, J. C. Ignacio-Espinoza, D. M. Needham, E. B. Fichot, J. A. Fuhrman, Dynamic marine viral infections and major contribution to photosynthetic processes shown by spatiotemporal picoplankton metatranscriptomes. Nat. Commun. 10, 1169 (2019).

48. D. Lindell, J. D. Jaffe, Z. I. Johnson, G. M. Church, S. W. Chisholm, Photosynthesis genes in marine viruses yield proteins during host infection. Nature 438, 86–89 (2005).

49. N. H. Mann, A. Cook, A. Millard, S. Bailey, M. Clokie, Bacterial photosynthesis genes in a virus. Nature 424, 741–741 (2003).

50. R. W. Hoham, D. Remias, Snow and Glacial Algae: A Review. J. Phycol. 56, 264–282 (2020).

51. Y. Q. Liu, T. J. Vick-Majors, J. C. Priscu, T. D. Yao, S. C. Kang, K. S. Liu, Z. Y. Cong, J. B. Xiong, Y. Li, Biogeography of cryoconite bacterial communities on glaciers of the Tibetan Plateau. FEMS Microbiol. Ecol. 93, fix072 (2017).

52. Z. Zhong, F. Tian, S. Roux, M. C. Gazitúa, N. E. Solonenko, Y.-F. Li, M. E. Davis, J. L. Van Etten, E. Mosley-Thompson, V. I. Rich, M. B. Sullivan, L. G. Thompson, Glacier ice archives nearly 15,000-year-old microbes and phages. Microbiome 9, 160 (2021).

53. Q. Liu, H.-C. Liu, J.-L. Zhang, Y.-G. Zhou, Y.-H. Xin, *Sphingomonas psychrolutea* sp. nov., a psychrotolerant bacterium isolated from glacier ice. Int. J. Syst. Evol. Microbiol. 65, 2955–2959 (2015).

54. M. A. Ayllón, M. Turina, J. Xie, L. Nerva, S.-Y. L. Marzano, L. Donaire, D. Jiang, I. R. Consortium, ICTV virus taxonomy profile: Botourmiaviridae. J. Gen. Virol. 101, 454 (2020).

55. M. Sõmera, D. Fargette, E. Hébrard, C. Sarmiento, I. R. Consortium, ICTV virus taxonomy profile: Solemoviridae 2021. J. Gen. Virol. 102, 001707 (2021).

56. D. Gilmer, C. Ratti, I. R. Consortium, ICTV virus taxonomy profile: Benyviridae. J. Gen. Virol. 98, 1571–1572 (2017).

57. J. Matthijnssens, H. Attoui, K. Bányai, C. P. Brussaard, P. Danthi, M. Del Vas, T. S. Dermody, R. Duncan, Q. Fāng, R. Johne, ICTV virus taxonomy profile: Sedoreoviridae 2022. J. Gen. Virol. 103, 001782 (2022).

58. P. Simmonds, P. Becher, J. Bukh, E. A. Gould, G. Meyers, T. Monath, S. Muerhoff, A. Pletnev, R. Rico-Hesse, D. B. Smith, ICTV virus taxonomy profile: Flaviviridae. J. Gen. Virol. 98, 2–3 (2017).

59. M. A. Purdy, J. F. Drexler, X.-J. Meng, H. Norder, H. Okamoto, W. H. Van der Poel, G. Reuter, W. M. de Souza, R. G. Ulrich, D. B. Smith, ICTV virus taxonomy profile: Hepeviridae 2022. J. Gen. Virol. 103, 001778 (2022).

60. S. Payne, “Chapter 14 - Family Astroviridae“ in Viruses, S. Payne, Ed. (Academic Press, 2017), pp. 125–128.

61. R. Wu, E. Zimmerman Amy, S. Hofmockel Kirsten, The direct and indirect drivers shaping RNA viral communities in grassland soils. mSystems 0, e00099–00024 (2024).

62. N. Mollentze, D. G. Streicker, Predicting zoonotic potential of viruses: where are we? Curr. Opin. Virol. 61, 101346 (2023).

63. N. Mollentze, S. A. Babayan, D. G. Streicker, Identifying and prioritizing potential human-infecting viruses from their genome sequences. PLoS Biol. 19, e3001390 (2021).

64. F. Zhang, X. Shi, C. Zeng, L. Wang, X. Xiao, G. Wang, Y. Chen, H. Zhang, X. Lu, W. Immerzeel, Recent stepwise sediment flux increase with climate change in the Tuotuo River in the central Tibetan Plateau. Sci. Bull. 65, 410–418 (2020).

65. A. M. Bolger, M. Lohse, B. Usadel, Trimmomatic: a flexible trimmer for Illumina sequence data. Bioinformatics 30, 2114–2120 (2014).

66. E. Kopylova, L. Noé, H. Touzet, SortMeRNA: fast and accurate filtering of ribosomal RNAs in metatranscriptomic data. Bioinformatics 28, 3211–3217 (2012).

67. D. Li, C.-M. Liu, R. Luo, K. Sadakane, T.-W. Lam, MEGAHIT: an ultra-fast single-node solution for large and complex metagenomics assembly via succinct de Bruijn graph. Bioinformatics 31, 1674–1676 (2015).

68. D. Hyatt, G.-L. Chen, P. F. LoCascio, M. L. Land, F. W. Larimer, L. J. Hauser, Prodigal: prokaryotic gene recognition and translation initiation site identification. BMC Bioinformatics 11, 119 (2010).

69. L. S. Johnson, S. R. Eddy, E. Portugaly, Hidden Markov model speed heuristic and iterative HMM search procedure. BMC Bioinformatics 11, 431 (2010).

70. S. Sakaguchi, S.-i. Urayama, Y. Takaki, K. Hirosuna, H. Wu, Y. Suzuki, T. Nunoura, T. Nakano, S. Nakagawa, NeoRdRp: A Comprehensive dataset for identifying RNA-dependent RNA polymerases of various RNA Viruses from metatranscriptomic data. Microbes Environ. 37, ME22001 (2022).

71. S. Nayfach, A. P. Camargo, F. Schulz, E. Eloe-Fadrosh, S. Roux, N. C. Kyrpides, CheckV assesses the quality and completeness of metagenome-assembled viral genomes. Nat. Biotechnol. 39, 578–585 (2021).

72. L. Fu, B. Niu, Z. Zhu, S. Wu, W. Li, CD-HIT: accelerated for clustering the next-generation sequencing data. Bioinformatics 28, 3150–3152 (2012).

73. A. P. Camargo, S. Nayfach, I. M. A. Chen, K. Palaniappan, A. Ratner, K. Chu, Stephan J. Ritter, T. B. K. Reddy, S. Mukherjee, F. Schulz, L. Call, Russell Y. Neches, T. Woyke, Natalia N. Ivanova, Emiley A. Eloe-Fadrosh, Nikos C. Kyrpides, S. Roux, IMG/VR v4: an expanded database of uncultivated virus genomes within a framework of extensive functional, taxonomic, and ecological metadata. Nucleic Acids Res. 51, D733–D743 (2023).

74. M. Shi, X.-D. Lin, J.-H. Tian, L.-J. Chen, X. Chen, C.-X. Li, X.-C. Qin, J. Li, J.-P. Cao, J.-S. Eden, J. Buchmann, W. Wang, J. Xu, E. C. Holmes, Y.-Z. Zhang, Redefining the invertebrate RNA virosphere. Nature 540, 539–543 (2016).

75. P. Jones, D. Binns, H.-Y. Chang, M. Fraser, W. Li, C. McAnulla, H. McWilliam, J. Maslen, A. Mitchell, G. Nuka, S. Pesseat, A. F. Quinn, A. Sangrador-Vegas, M. Scheremetjew, S.-Y. Yong, R. Lopez, S. Hunter, InterProScan 5: genome-scale protein function classification. Bioinformatics 30, 1236–1240 (2014).

76. E. Potenza, T. D. Domenico, I. Walsh, S. C. E. Tosatto, MobiDB 2.0: an improved database of intrinsically disordered and mobile proteins. Nucleic Acids Res. 43, D315–D320 (2014).

77. T. K. Attwood, A. Coletta, G. Muirhead, A. Pavlopoulou, P. B. Philippou, I. Popov, C. Romá-Mateo, A. Theodosiou, A. L. Mitchell, The PRINTS database: a fine-grained protein sequence annotation and analysis resource—its status in 2012. Database 2012, (2012).

78. L. Käll, A. Krogh, E. L. L. Sonnhammer, Advantages of combined transmembrane topology and signal peptide prediction—the Phobius web server. Nucleic Acids Res. 35, W429–W432 (2007).

79. A. Krogh, B. Larsson, G. von Heijne, E. L. L. Sonnhammer, Predicting transmembrane protein topology with a hidden markov model: application to complete genomes. J. Mol. Biol. 305, 567–580 (2001).

80. M. Steinegger, M. Meier, M. Mirdita, H. Vöhringer, S. J. Haunsberger, J. Söding, HH-suite3 for fast remote homology detection and deep protein annotation. BMC Bioinformatics 20, 473 (2019).

81. M. Mirdita, L. von den Driesch, C. Galiez, M. J. Martin, J. Söding, M. Steinegger, Uniclust databases of clustered and deeply annotated protein sequences and alignments. Nucleic Acids Res. 45, D170–D176 (2017).

82. R. D. Finn, J. Clements, S. R. Eddy, HMMER web server: interactive sequence similarity searching. Nucleic Acids Res. 39, W29–W37 (2011).

83. J. Mistry, S. Chuguransky, L. Williams, M. Qureshi, Gustavo A. Salazar, E. L. L. Sonnhammer, S. C. E. Tosatto, L. Paladin, S. Raj, L. J. Richardson, R. D. Finn, A. Bateman, Pfam: The protein families database in 2021. Nucleic Acids Res. 49, D412–D419 (2020).

84. S. Lu, J. Wang, F. Chitsaz, M. K. Derbyshire, R. C. Geer, N. R. Gonzales, M. Gwadz, D. I. Hurwitz, G. H. Marchler, J. S. Song, N. Thanki, R. A. Yamashita, M. Yang, D. Zhang, C. Zheng, C. J. Lanczycki, A. Marchler-Bauer, CDD/SPARCLE: the conserved domain database in 2020. Nucleic Acids Res. 48, D265–D268 (2019).

85. I. Sillitoe, N. Bordin, N. Dawson, V. P. Waman, P. Ashford, H. M. Scholes, C. S. M. Pang, L. Woodridge, C. Rauer, N. Sen, M. Abbasian, S. Le Cornu, S. D. Lam, K. Berka, Ivana H. Varekova, R. Svobodova, J. Lees, C. A. Orengo, CATH: increased structural coverage of functional space. Nucleic Acids Res. 49, D266–D273 (2020).

86. H. Cheng, Y. Liao, R. D. Schaeffer, N. V. Grishin, Manual classification strategies in the ECOD database. *Proteins. Struct., Funct.*, Genet. 83, 1238–1251 (2015).

87. A. A. Pratama, B. Bolduc, A. A. Zayed, Z.-P. Zhong, J. Guo, D. R. Vik, M. C. Gazitúa, J. M. Wainaina, S. Roux, M. B. Sullivan, Expanding standards in viromics: in silico evaluation of dsDNA viral genome identification, classification, and auxiliary metabolic gene curation. PeerJ 9, e11447 (2021).

88. M. Shaffer, M. A. Borton, B. B. McGivern, A. A. Zayed, Sabina L. La Rosa, L. M. Solden, P. Liu, A. B. Narrowe, J. Rodríguez-Ramos, B. Bolduc, M. C. Gazitúa, R. A. Daly, G. J. Smith, D. R. Vik, P. B. Pope, M. B. Sullivan, S. Roux, Kelly C. Wrighton, DRAM for distilling microbial metabolism to automate the curation of microbiome function. Nucleic Acids Res. 48, 8883–8900 (2020).

89. Y. Liu, N. Jiao, K. Xu Zhong, L. Zang, R. Zhang, X. Xiao, Y. Shi, Z. Zhang, Y. Tao, L. Bai, B. Gao, Y. Yang, X. Huang, M. Ji, J. Liu, P. Liu, T. Yao, Diversity and function of mountain and polar supraglacial DNA viruses. Sci. Bull. 68, 2418–2433 (2023).

90. C. T. Roots, A. Lukasiewicz, J. E. Barrick, OSTIR: open source translation initiation rate prediction. J. Open. Source Softw. 6, 3362 (2021).

91. S. R. Krishnamurthy, D. Wang, Extensive conservation of prokaryotic ribosomal binding sites in known and novel picobirnaviruses. Virology 516, 108–114 (2018).

92. Á. Boros, B. Polgár, P. Pankovics, H. Fenyvesi, P. Engelmann, T. G. Phan, E. Delwart, G. Reuter, Multiple divergent picobirnaviruses with functional prokaryotic Shine-Dalgarno ribosome binding sites present in cloacal sample of a diarrheic chicken. Virology 525, 62–72 (2018).

