## Supplementary Materialss for "Diversity and ecological roles of RNA viral communities in the cryosphere of the Tibetan Plateau"

Yongqin Liu et al.

**This PDF file includes:**

Supplementary Text
References
Supplementary figs. S1 to S22

**Other Supplementary Materials for this manuscript include the following:**

Supplementary table S1 to S10 in one Excel file.

### Supplementary Text

### Supplementary Results

#### *Shared vOTUs between different habitats*

In addition to the RNA viral diversity and community structure, we further investigated the common RNA viral groups that shared among various habitats or ecosystems of the TPC (fig. S19). Most vOTUs (4,750; 89.0% of 5,333) only occurred in one type of ecosystem or habitat, and a small proportion (583, 11.0%) were shared among multiple ecosystems or habitats. This is consistent with our other result which shows the number of shared vOTUs across samples follows a power-law function (Fig. S20). Moreover, the distribution of shared RNA viruses at genus to family levels showed similar patterns (fig. S21). The shared vOTUs from the glacier ecosystems represented the largest group of shared vOTUs. The number of RNA vOTUs shared among the snow, ice and/or cryoconite was 411, which accounted for 70.5% of all the shared vOTUs (fig. S19A). The second and third most shared vOTUs were between the wetland and the upland (120, 20.5% of 583) and between the proglacial lake sediments and the glacier habitats (9, 1.5%), respectively (fig. S19A).

Although the number of the shared vOTUs accounted for a small proportion of the total vOTUs, they possessed a high relative abundance in the TPC RNA viral communities, especially in the habitats of snow, ice, and cryoconite (fig. S19B). The relative abundance of the shared vOTUs accounted for more than 64% in the snow, ice and cryoconite and well over 25% in other ecosystems except for the proglacial lake sediments. The shared vOTUs were dominated by the eight established RNA viral families including the viral families *Botourmiaviridae* (with average relative abundance >6% in snow and cryoconite), f.0102.base-Curvula (ice), f.0219 (wetland), f.0385 (ice), *Fiersviridae* (wetland), *Steitzviridae* (cryoconite), *Tombusviridae* (ice and wetland), and *Atkinsviridae* (cryoconite and upland) (fig. S19B).

#### *Genome architecture of putative archaeal RNA viruses*

Five vContigs were putatively linked to the AMGs of archaea through CRISPR-spacers. There of them contain only one ORF, which contains the functional domain for

RdRp (fig. S11). Two of them contains multiple ORFs. vContigs belonging to *Fiersviridae* and *Steitzviridae* contains RBS motifs for all predicted ORFs while one ORFs of *Picobirnaviridae* (KQGRC1_k141_677647) also contains RBS motif. Interestingly, *Steitzviridae* vContigs (TGL_TT3_k141_238474) contained a putative long un-transcribed region (5064 of the full length 8,671bp). This region contains no functional domain according to all our functional annotation pipelines.

***ORF and domain analysis of putative AMGs***

In addition to DRAM and functional annotations of ORF by using hhsuit etc (see Materials and Methods in the Main text), we further checked the completeness of the AMGs functional domain through amino acid sequence alignment of putative AMGs with their closest cellular homologs found in NCBI-NR database (figs. S14 and S22). Detailed alignment procedures for each AMG were described in Supplementary Methods as for PsbA. Eleven of 39 AMGs (28.2%) were only partial of the corresponding cellular homolog ORFs, including phycobilisome protein (2006QYC2_k141_333847_2), inner membrane component of T3SS cytoplasmic domain (AW1-1_k141_425352_4), glucose-6-phosphate dehydrogenase (AJ-7_k141_1444729_1), ribosomal protein L3 (GLYSC2_k141_244580_1), ribosomal protein L4 (GLYSC2_k141_244580_2), acyl-CoA dehydrogenase (KQGRC1_k141_320723_2), Rho termination factor (QY215SS_k141_188611_2), ricin-type beta-trefoil lectin (QY215SS_k141_188611_6), putative tricarboxylic transport membrane protein (TGL_TJ3_k141_162580_4), ATP-dependent Clp protease (TGL_TJ3_k141_464371_3), IMP dehydrogenase / GMP reductase (TGL5300down_k141_394333_3). These AMGs contain partial domain of a functional ORFs with a single full domain or multiple domains. Therefore, the functional roles of theses AMGs in cellular metabolism pathways deserve further verification.

### Supplementary Methods

#### *Samples collection and preparation*

The 79 samples were collected from 4 types of ecosystems, including glaciers (28), proglacial lake sediments (6), upland soil (16) and wetland soil (29) in permafrost regions. These samples, including snow (5), ice (5), and cryoconite (18) from 8 glaciers, upland soil (16, with 13 from surface and 3 from subsurface) and wetland soil (29), and proglacial lake sediments (6), were collected in from November 2020 to July 2023 (Fig. 1; table S1).

Snow and ice samples (~5 cm deep) were collected within a square meter area using a pre-cleaned steel scoop and ice axe and then placed into 2 L sterile Whirl-Pak bags (Nasco, Fort Atkinson, USA) as described in the previous study(*1*). For surface snow and ice, 2 mm of the sample was scraped with a sterile scalpel for decontamination. The cryoconite samples were collected from holes using a stainless-steel scoop and placed in pre-cleaned 1-L polycarbonate bottles (Nalgene, Thermo Fisher, USA). Proglacial lake sediments were collected with grabbers from the surface of the sediments (~15 cm) in Qiangyong Lake that was fed by Qiangyong glacier and then transferred into 500 mL sterile Whirl-Pak bags (Nasco, Fort Atkinson, USA). Wetland soil samples were collected in the lakeside of Aerjin (AJ), the Altun Mountains National Nature Reserve and permafrost regions of Longxiazai with pre-clean spades. The surface soil of each site was stored in three 500 mL sterile Whirl-Pak bags (Nasco, Fort Atkinson, USA). Upland soil samples were collected in permafrost regions of the Tuotuohe (TTH), the source region of the Yangtze River(*2*). Fourteen soil cores were sampled with a motor-driven soil column cylinder. Thirteen surface (0–20 cm) soil samples and three sub-surface (50–120 cm) soil layers of one soil column were collected and placed into 500 mL sterile Whirl-Pak bags (Nasco, Fort Atkinson, USA). All samples were stored at -20 °C during the transportation from the field to the laboratory in Lhasa City (Tibet, China). Snow and ice samples were de-frozen at 4 °C at dark and then filtered through 0.2 μm polycarbonate filters (Millipore) immediately. All samples were stored at –80 °C or liquid nitrogen until total RNA extraction.

***RNA virus identification and verification***

The protein sequence containing domains with an E-value ≤0.01 was kept for RdRp verification:

First, all protein sequences with putative RdRp domains were subjected to the Palmscan algorithm(*3*) screening based on PALMdb. Sequences with a palmprint (a segment of the palm sub-domain robustly delineated by well-conserved catalytic motifs) score ≥20 with treated as positive RdRp sequences. Second, we applied a metagenomic reads mapping strategy as described by Hou et al.(*4*) to remove DNA remnants since most of the metatranscriptomes were accompanied by metagenome from the same sample (table S9; see below for metagenome sequencing). In addition, for metatranscriptome without accompanied metagenome, sequences from the metagenome of similar samples (e.g., cryoconite) were used (table S9). Metagenome reads were mapped against the RdRp nucleotide sequences from prodigal with Bowtie2 v2.4.255 with the "end-to-end" setting to check whether there was a DNA counterpart. Sequences with a metagenome reads coverage ≥75% and average sequences depth ≥1× were removed. RdRp sequences passed the two verification processes were confirmed as true positive RdRps and contigs containing these RdRps were considered as *bona fide* RNA viral genomes. In total, we obtained 8,799 RNA viral genomes (vContigs) from all samples, with an average of ~111 RNA vContigs per sample.

***Taxonomy classification of TPC RNA viruses***

To assign taxonomic information for RNA viruses, we combined different public reference databases, including the global RNA virome dataset (RVMT)(*5*), the *Tara* Oceans datasets (TO)(*6*), the Serratus palmprint database (Palmdb)(*3*) and IMG/VR-v4 database(*7*) to improve taxonomy classification of TPC RNA viruses.

First, the completeness of RdRp domain sequences of the RVMT dataset(*5*) was evaluated as described above, only "Complete" RdRp domain sequences (n=106,561) were kept. Second, the "complete" (n=6,238) centroids RdRp domain sequences used by Zayed et al.(*6*) were downloaded for Markov Cluster Algorithm (MCL) clustering. Then, "complete" RdRp domain sequences (n=3,206) from the TPC dataset were combined with the two "complete" RdRp domain sequences dataset mentioned above and subject to clustering by using USEARCH v10.0.240(*8*) with the following parameters: usearch --cluster_fast -id 0.50 -sort length. The centroids of the resultant 21,763 clusters (638 from TPC datasets) were then used for MCL network analysis as previously described(*6*). In brief, all centroid sequences were cross-compared running all-against-all pairwise blastp v2.13 with parameters (-gapopen 9 -gapextend 1 -word_size 3). E-values for each pair were extracted and negative-log10-transformed in MCL v14.137(*9*) (--stream-mirror --stream-neg-log10 -stream-tf 'ceil(200)'). Transformed e-values were used in an MCL network for iterative clustering, changing the granularity parameter at each iteration (range 1.1–2). Inflation values of 1.1 were used to delineate Phylum level clusters. With Inflation values of 1.1, a total of 237 phylum-level clusters were formed. However, all those TPC clusters contained RdRp domain sequences from RVMT, indicating full-length RdRps of TPC were covered by the RVMT dataset at the phylum level.

In addition, we determine the taxonomy of each vContig based on the sequence identity against reference datasets (the RVMT, TO) or databases (NCBI RefSeq, IMG/VR-v4) with dedicated and detailed RNA virus taxonomic classification. According to the identity between query sequence with reference database, different taxonomy level of reference database was assigned to TPC viral genomes as following:

Level 1, Species level classification based on RdRp AAI and vContigs ANI as Neri et al(*5*).consists of vContigs encoding RdRps with exceptionally high amino acid identity to RdRps from the RVMT and TO datasets (via best BLASTp match with Identity 90%, Query-Coverage 75%, and E-value <1e-3) and contigs with high nucleic similarity to vContigs from RVMT and TO dataset (95% ANI over 95% AF or Nident R 900 nt and E-value <1e-3, established using CheckV anicalc.py and aniclust.py scripts(*10*)). The taxonomy of Level A was consistent with the taxonomy of reference databases at the "species" level; consists of contigs sharing high nucleic similarity to those vContigs from RVMT, TO and IMG-VR4 database (via best dc-MEGABLAST hit at Identity 90%, Query-Coverage 75% OR Nident 900nt and E-value <1e-3).

Level 2, Family to genus level classification based on RdRp AAI consists of contigs encoding RdRp domain sharing high amino acid similarity to those RdRps from the reference database (Identity 50%, Query-Coverage 50%).

Level 3, phylum-level classification based on complete RdRps and MCL clustering. MCL inflation values of 1.1 were used to delineate phylum level clusters. The confidence of RNA virus classification was Level 1> Level 2 > Level 3, and the taxonomic information from the approach with higher confidence were firstly used. Finally, for all RNA viruses within each established phylum, a phylogenetic tree of RdRps was constructed to verify and refine the taxonomic classification. For RNA viruses without a taxonomic classification from Level 1 to Level 3, we also carried out palmprint similarity analysis with palmDB and BLASTp to all reference databases on RdRps to get putative phylum-class level classification or the most closely associated sequences in the reference database (table S10).

***Phylogenetic analysis of the RdRp amino acid sequences***

We constructed phylogenetic trees for all RdRp amino acid (AA) sequences classified into 6 established RNA viral phyla. Reference sequences were obtained from RVMT C90, TO, NCBI Refseq, IMG/VR-V4 during the classification step (see above). In brief, RdRps AA sequence from each dataset or database that affiliated or clustered with our RdRps at each level were kept as references. Reference RdRps and TPC RdRps that from the same phylum were pooled together and aligned using MAFFT v7.508(*11*) with default parameters. The multiple sequence alignment was filtered using trimAl v1.4.rev15(*12*) with -gappyout parameters and custom scripts (fasta_drop.py) that remove sequences with >70% gaps. The final multiple sequence alignments were used for tree construction. The maximum-likelihood tree was constructed by using the FastTree software v2.1.11 with the parameters "-gamma -lg -boot 1000". The tree was visualized with iTOL(*13*).

***Phylogenetic analysis of PsbA amino acid sequences***

Reference sequences were obtained from NCBI IPG database by searching with "photosystem II protein D1" and filtered out none PsbA domain using hmmsearch with TIGR01151.1.hmm and PF00124.hmm. The gathering score 27 and length of 180 AA residue were used for filtering. After filtering, all reference PsbA AA sequences were dereplicated using CD-HIT with 0.99 as the cutoff. PsbA of snow RNA viruses and non-redundant reference PsbA were combined for muscle v5 alignment, trimal v1.4.rev15 (with -gappyout) and gaps removal (drop sequences with >70% gaps). The final multiple sequence alignments were used for maximum-likelihood tree construction using the FastTree v2.1.11 with the parameters "-gamma -lg -boot 1000".

*Estimation of shared RNA viruses among different habitats*

To establish the linkage of RNA virus between different habitats, we applied two strategies:

i) The mRNA reads recruitment strategy. We define that a virus presents in one sample if it has a vContigs TPM value ≥1000 within that sample(*4*). Then it is straightforward that two habitats share the same RNA virus if vContigs of the species were present in the two samples of different habitats.

ii) Genome-based vOTUs sharing strategies. Fifty percent and 90% of RdRp AAI (via UCLUST clustering) represent shared vOTUs between different samples at "family-genus"- and "species"- levels, respectively. If one vOTU was shared between samples from different habitats, then this vOTU was defined as shared vOTUs between the two habitats.

**Reference**


1. Y. Liu, M. Ji, T. Yu, J. Zaugg, A. M. Anesio, Z. Zhang, S. Hu, P. Hugenholtz, K. Liu, P. Liu, Y. Chen, Y. Luo, T. Yao, A genome and gene catalog of glacier microbiomes. *Nat. Biotechnol.* **40**, 1341-1348 (2022).
2. F. Zhang, X. Shi, C. Zeng, L. Wang, X. Xiao, G. Wang, Y. Chen, H. Zhang, X. Lu, W. Immerzeel, Recent stepwise sediment flux increase with climate change in the Tuotuo River in the central Tibetan Plateau. *Sci. Bull.* **65**, 410-418 (2020).
3. A. Babaian, R. Edgar, Ribovirus classification by a polymerase barcode sequence. *PeerJ* **10**, e14055 (2022).
4. X. Hou, Y. He, P. Fang, S.-Q. Mei, Z. Xu, W.-C. Wu, J.-H. Tian, S. Zhang, Z.-Y. Zeng,

Q.-Y. Gou, G.-Y. Xin, S.-J. Le, Y.-Y. Xia, Y.-L. Zhou, F.-M. Hui, Y.-F. Pan, J.-S. Eden, Z.-H. Yang, C. Han, Y.-L. Shu, D. Guo, J. Li, E. C. Holmes, Z.-R. Li, M. Shi, Artificial intelligence redefines RNA virus discovery. *bioRxiv*, 2023.2004.2018.537342 (2023).

5. U. Neri, Y. I. Wolf, S. Roux, A. P. Camargo, B. Lee, D. Kazlauskas, I. M. Chen, N. Ivanova, L. Zeigler Allen, D. Paez-Espino, D. A. Bryant, D. Bhaya, A. B. Narrowe, A. J. Probst, A. Sczyrba, A. Kohler, A. Séguin, A. Shade, B. J. Campbell, B. D. Lindahl, B. K. Reese, B. M. Roque, C. DeRito, C. Averill, D. Cullen, D. A. C. Beck, D. A. Walsh, D. M. Ward, D. Wu, E. Eloe-Fadrosh, E. L. Brodie, E. B. Young, E. A. Lilleskov, F. J. Castillo, F. M. Martin, G. R. LeCleir, G. T. Attwood, H. Cadillo-Quiroz, H. M. Simon, I. Hewson, I. V. Grigoriev, J. M. Tiedje, J. K. Jansson, J. Lee, J. S. VanderGheynst, J. Dangl, J. S. Bowman, J. L. Blanchard, J. L. Bowen, J. Xu, J. F. Banfield, J. W. Deming, J. E. Kostka, J. M. Gladden, J. Z. Rapp, J. Sharpe, K. D. McMahon, K. K. Treseder, K. D. Bidle, K. C. Wrighton, K. Thamatrakoln, K. Nusslein, L. K. Meredith, L. Ramirez, M. Buee, M. Huntemann, M. G. Kalyuzhnaya, M. P. Waldrop, M. B. Sullivan, M. O. Schrenk, M. Hess, M. A. Vega, M. A. O'Malley, M. Medina, N. E. Gilbert, N. Delherbe, O. U. Mason, P. Dijkstra, P. F. Chuckran, P. Baldrian, P. Constant, R. Stepanauskas, R. A. Daly, R. Lamendella, R. J. Gruninger, R. M. McKay, S. Hylander, S. L. Lebeis, S. P. Esser, S. G. Acinas, S. S. Wilhelm, S. W. Singer, S. S. Tringe, T. Woyke, T. B. K. Reddy, T. H. Bell, T. Mock, T. McAllister, V. Thiel, V. J. Denef, W.-T. Liu, W. Martens-Habbena, X.-J. Allen Liu, Z. S. Cooper, Z. Wang, M. Krupovic, V. V. Dolja, N. C. Kyrpides, E. V. Koonin, U. Gophna, Expansion of the global RNA virome reveals diverse clades of bacteriophages. *Cell* **185**, 4023-4037 (2022).

6. A. A. Zayed, J. M. Wainaina, G. Dominguez-Huerta, E. Pelletier, J. Guo, M. Mohssen, F. Tian, A. A. Pratama, B. Bolduc, O. Zablocki, D. Cronin, L. Solden, E. Delage, A. Alberti, J.-M. Aury, Q. Carradec, C. d. Silva, K. Labadie, J. Poulain, H.-J. Ruscheweyh, G. Salazar, E. Shatoff, R. Bundschuh, K. Fredrick, L. S. Kubatko, S. Chaffron, A. I. Culley, S. Sunagawa, J. H. Kuhn, P. Wincker, M. B. Sullivan, S. G. Acinas, M. Babin, P. Bork, E. Boss, C. Bowler, G. Cochrane, C. d. Vargas, G. Gorsky, L. Guidi, N. Grimsley, P. Hingamp, D. Iudicone, O. Jaillon, S. Kandels, L. Karp-Boss, E. Karsenti, F. Not, H. Ogata, N. Poulton, S. Pesant, C. Sardet, S. Speich, L. Stemmann, M. B.

Sullivan, S. Sungawa, P. Wincker, Cryptic and abundant marine viruses at the evolutionary origins of Earth's RNA virome. *Science* **376**, 156-162 (2022).

7. A. P. Camargo, S. Nayfach, I. M. A. Chen, K. Palaniappan, A. Ratner, K. Chu, Stephan J. Ritter, T. B. K. Reddy, S. Mukherjee, F. Schulz, L. Call, Russell Y. Neches, T. Woyke, Natalia N. Ivanova, Emiley A. Eloe-Fadrosh, Nikos C. Kyrpides, S. Roux, IMG/VR v4: an expanded database of uncultivated virus genomes within a framework of extensive functional, taxonomic, and ecological metadata. *Nucleic Acids Res.* **51**, D733-D743 (2023).
8. R. C. Edgar, Search and clustering orders of magnitude faster than BLAST. *Bioinformatics* **26**, 2460-2461 (2010).
9. A. J. Enright, S. Van Dongen, C. A. Ouzounis, An efficient algorithm for large-scale detection of protein families. *Nucleic Acids Res.* **30**, 1575-1584 (2002).
10. S. Nayfach, A. P. Camargo, F. Schulz, E. Eloe-Fadrosh, S. Roux, N. C. Kyrpides, CheckV assesses the quality and completeness of metagenome-assembled viral genomes. *Nat. Biotechnol.* **39**, 578-585 (2021).
11. K. Katoh, K. Misawa, K. i. Kuma, T. Miyata, MAFFT: a novel method for rapid multiple sequence alignment based on fast Fourier transform. *Nucleic Acids Res.* **30**, 3059-3066 (2002).
12. S. Capella-Gutiérrez, J. M. Silla-Martínez, T. Gabaldón, trimAl: a tool for automated alignment trimming in large-scale phylogenetic analyses. *Bioinformatics* **25**, 1972-1973 (2009).
13. I. Letunic, P. Bork, Interactive Tree Of Life (iTOL) v5: an online tool for phylogenetic tree display and annotation. *Nucleic Acids Res.* **49**, W293-W296 (2021).

### Supplementary figures

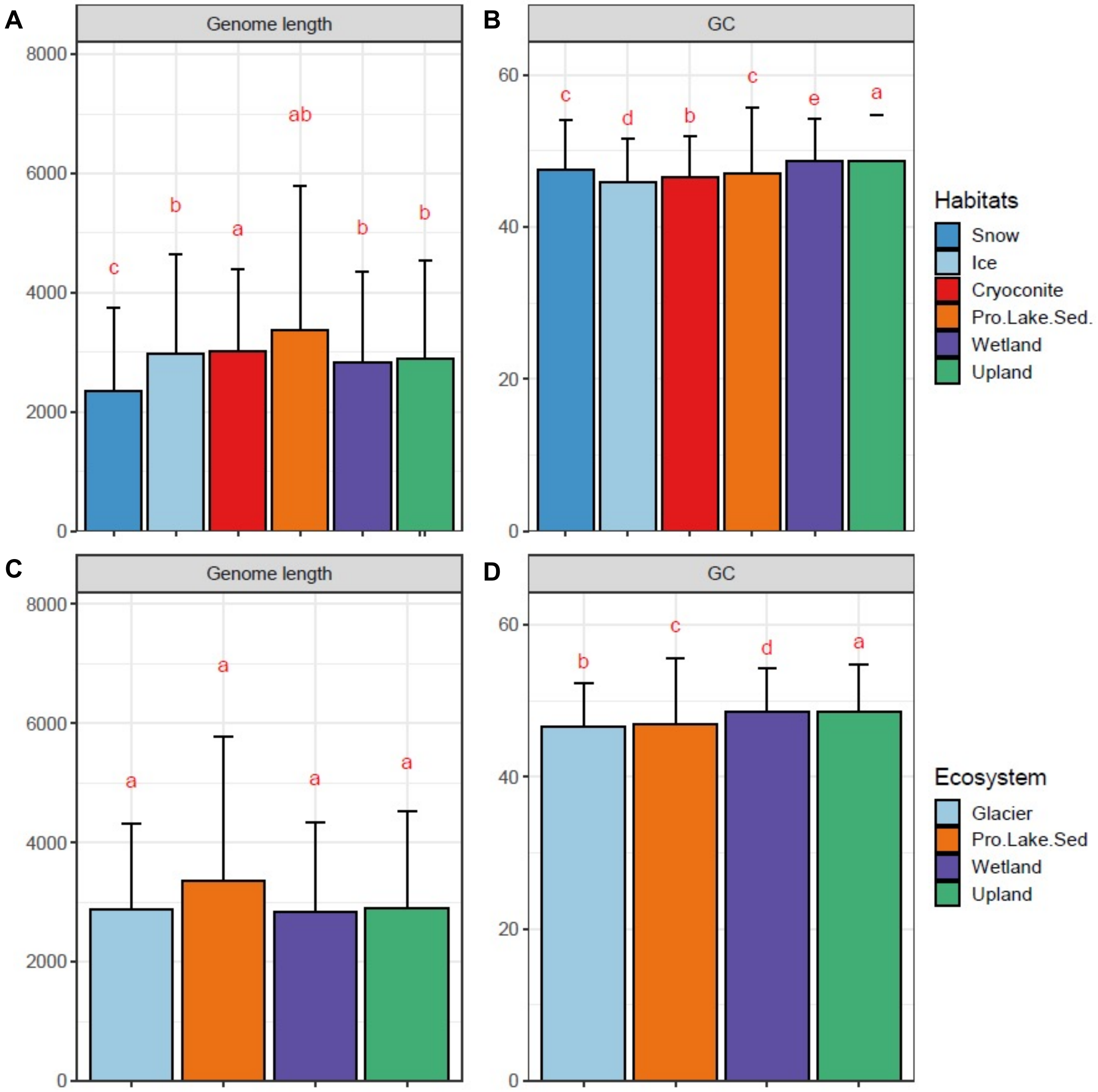


**Fig. S1. Genomic feature of TPC RNA viruses. (A)** Average length of vContigs of glacier habitats and other ecosystems. **(B)** Average (G+C)% content of vContigs of glacier habitats and other ecosystems. **(C)** Average length of vContigs of each ecosystem. **(D)** Average (G+C)% content of vContigs of of each ecosystem. In panel **(C)** and **(D)**, the values of snow, ice and cryoconites were grouped as a single category of glacier.

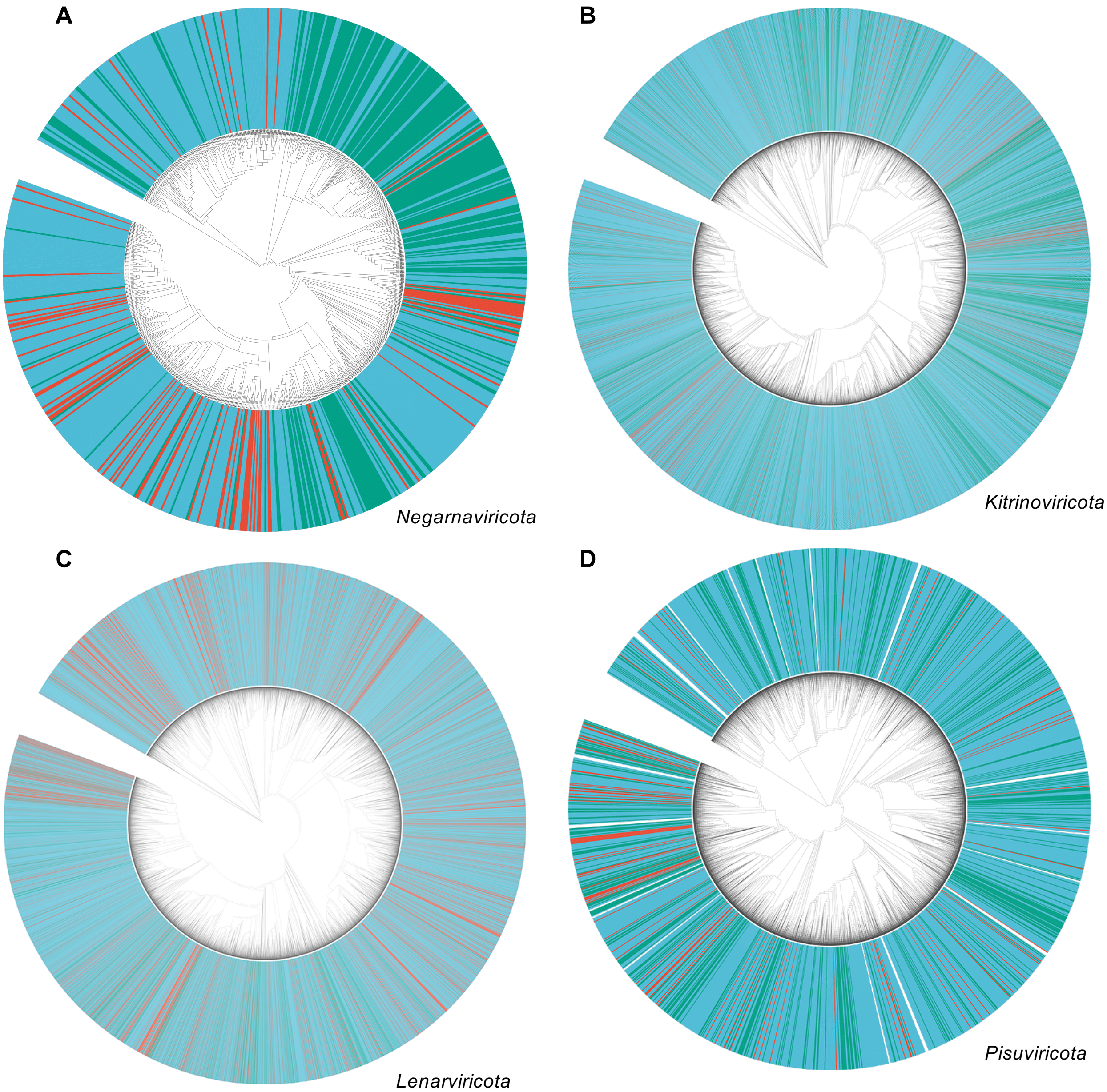


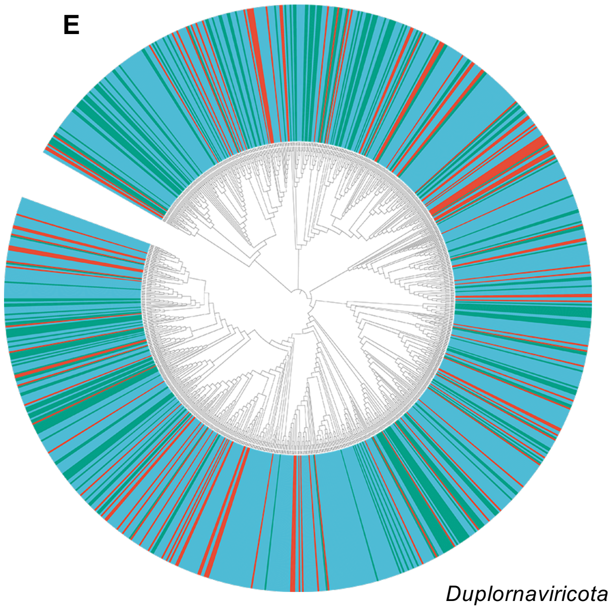


**Fig. S2. Phylogenetic tree of RdRPs for each established Phylum, highlighting the novelty of TPC RNA viral species. (A)** Negarnaviricota. **(B)** Kitrinoviricota. **(C)** Lenarviricota. **(D)** Pisuviricota. (**E**) Duplornaviricota. Within each tree, RdRps from TPC are highlighted in red, reference RdRps from RMVT and *Tara* Oceans were colored in light blue and green, respectively.

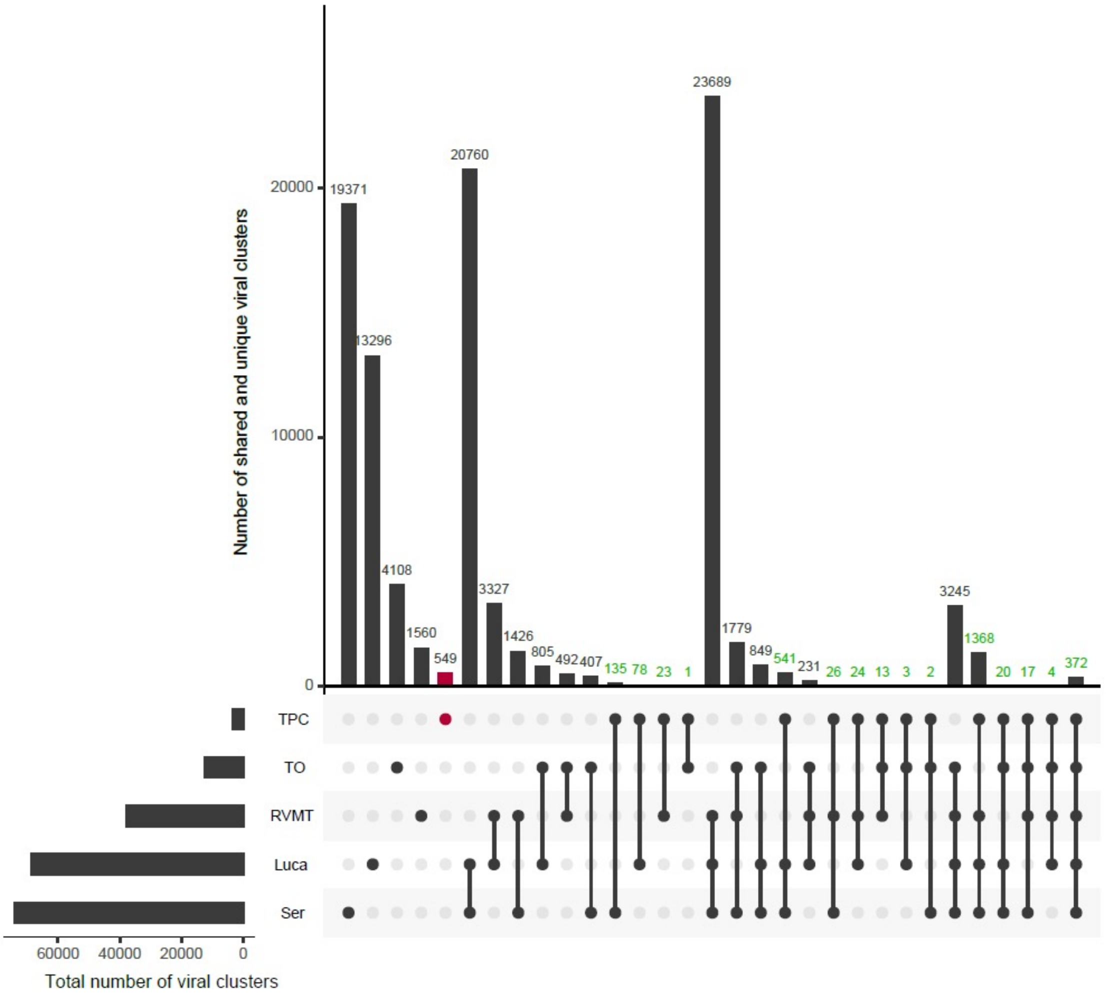


**Fig. S3. Shared viral clusters among TPC, *Tara* Oceans, RVMT, LucaProt and serratus viral datasets.** RNA virus RdRps were clustered using UCLUST with a cutoff of 0.5. The number of unique RNA viral clusters of TPC is highlighted in red, while numbers of shared viral clusters between TPC and other dataset s are colored in green.

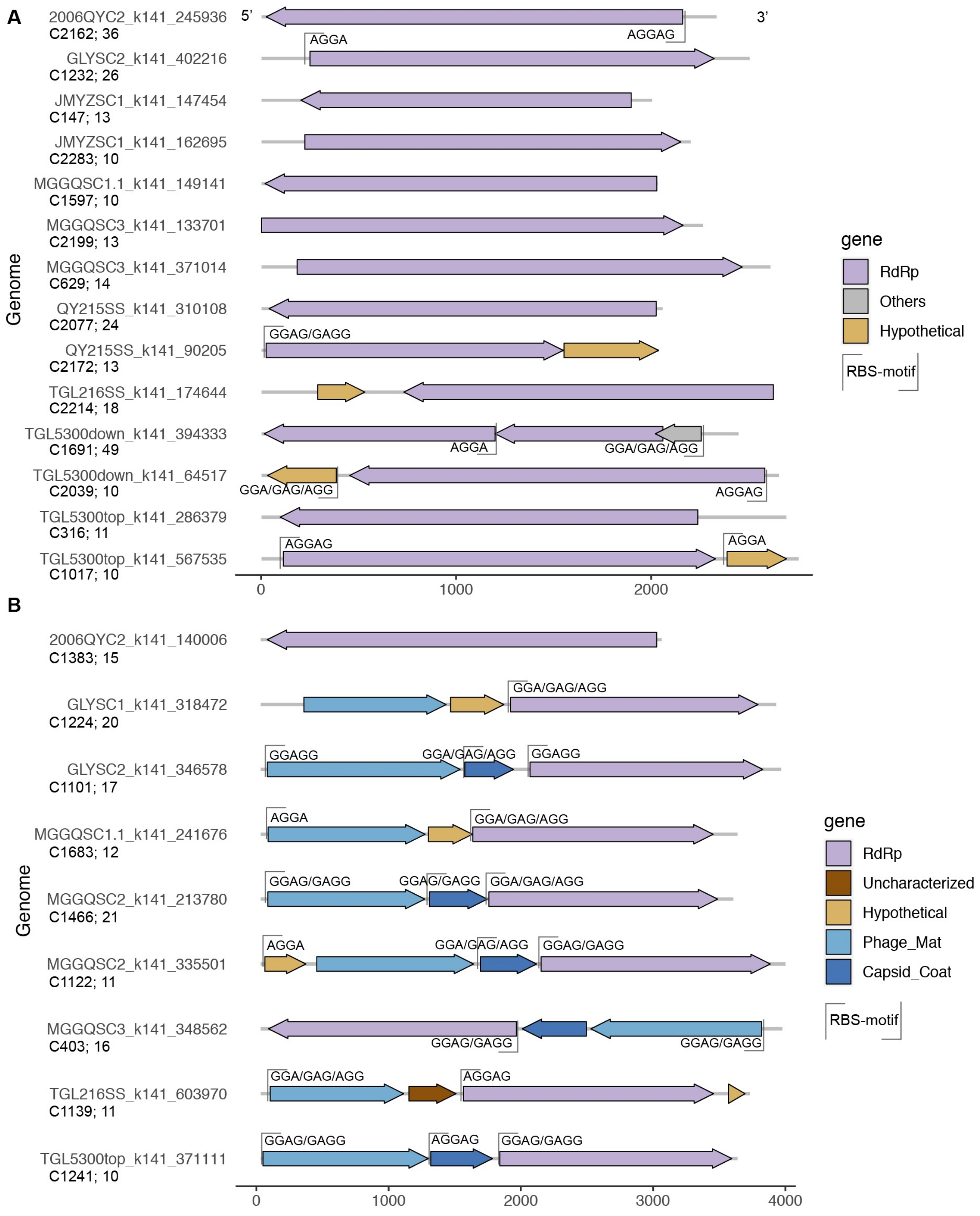


**Fig. S4. Genome structure of representative RNA viruses belonging to novel clades.** For each clade, one genome with the longest genome length were selected for visulization. vContigs were organized according to their length. (**A**) representative vContigs with length between 2 kb - 3 kb. **(B)** representative vContigs with length between 3 kb - 4 kb. For longer representative vContigs, see next page.

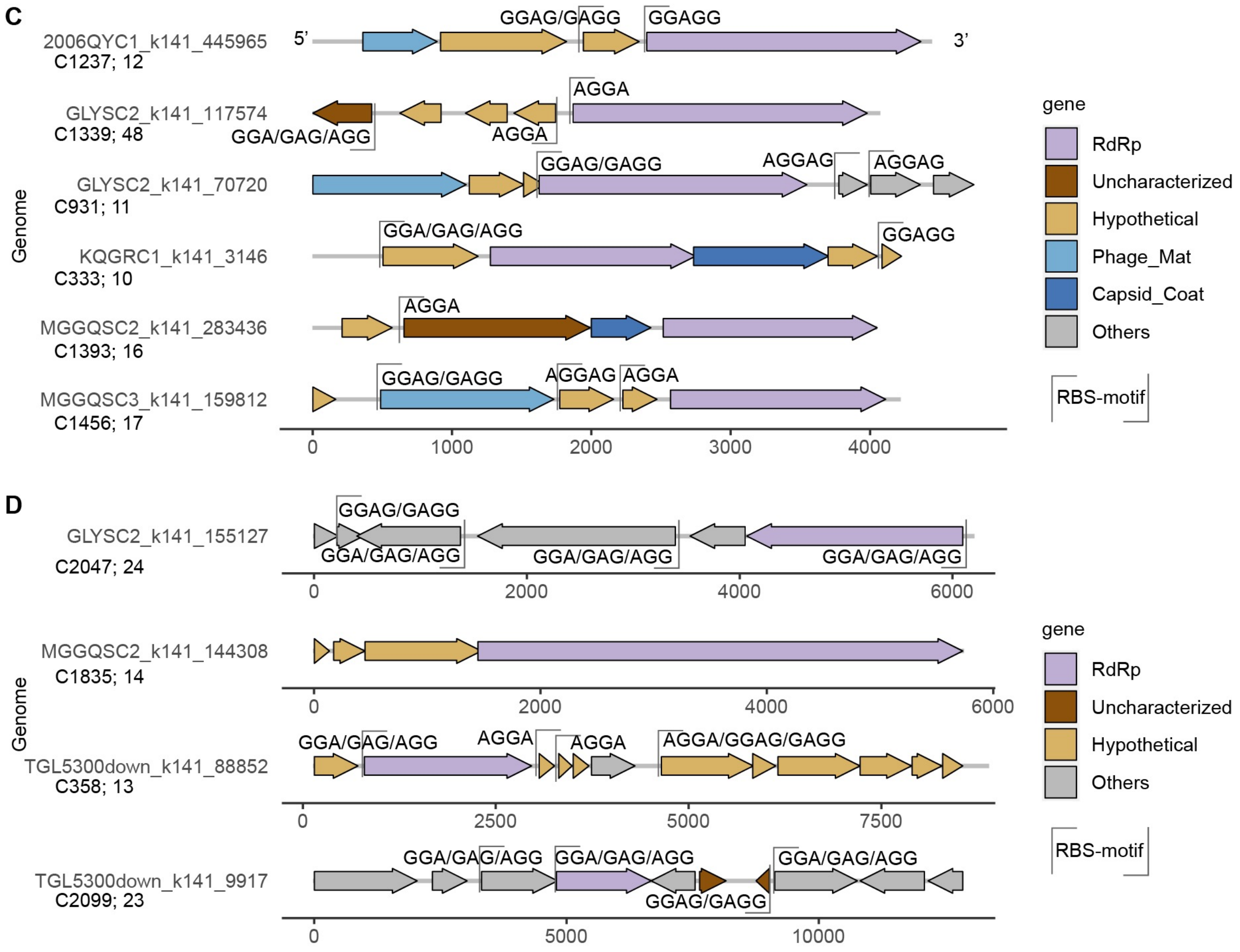


**Fig. S4. Continue. (C)** representative vContigs with length between 4 kb – 5 kb. **(D)** representative vContigs with a length between > 5kb. For **(D),** separate x-axis is used for each vContig.

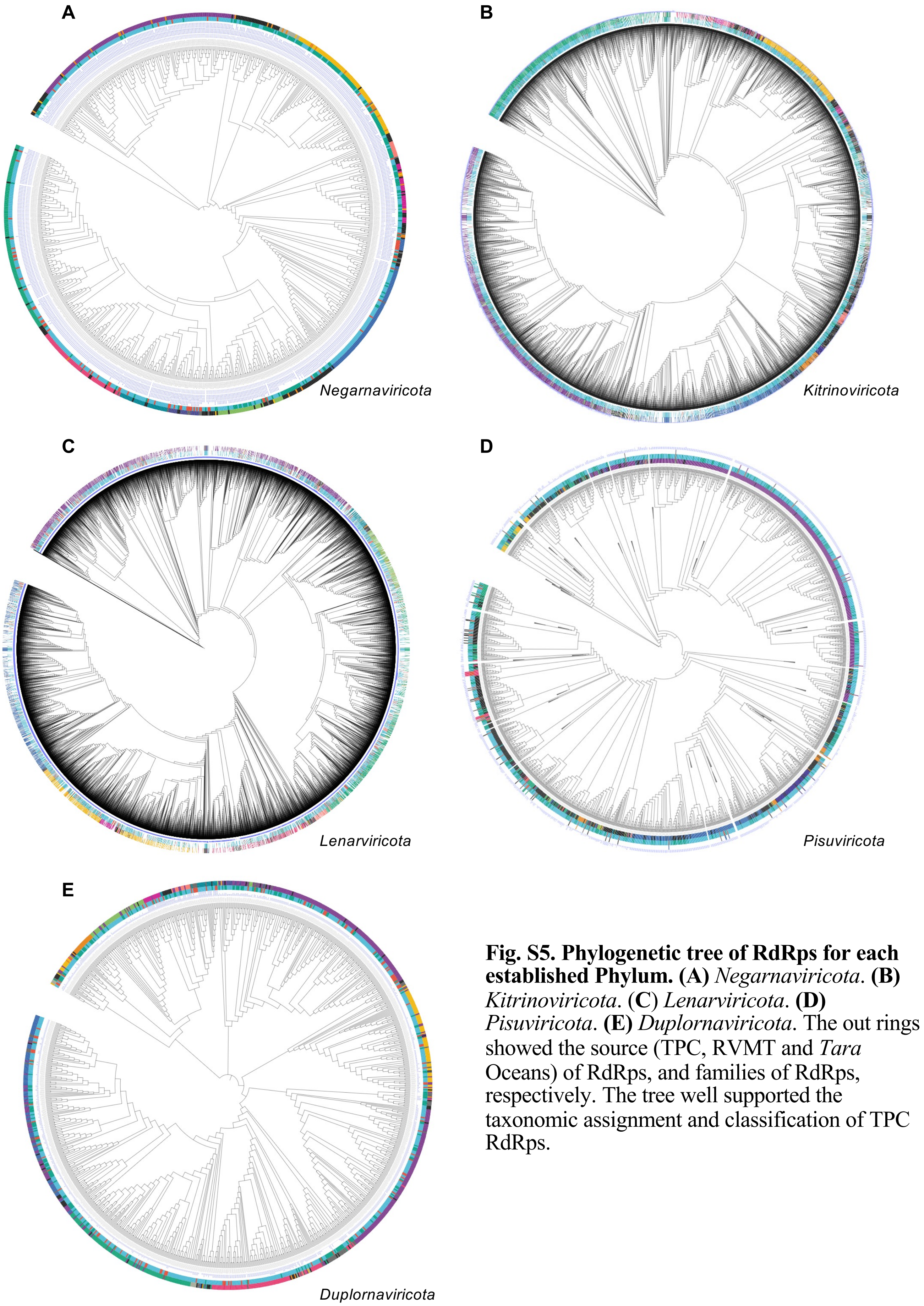


**Fig. S5. Phylogenetic tree of RdRps for each established Phylum. (A)** *Negarnaviricota*. **(B)** *Kitrinoviricota*. **(C)** *Lenarviricota*. **(D)** *Pisuviricota*. **(E)** *Duplornaviricota*. The out rings showed the source (TPC, RVMT and *Tara* Oceans) of RdRps, and families of RdRps, respectively. The tree well supported the taxonomic assignment and classification of TPC RdRps.

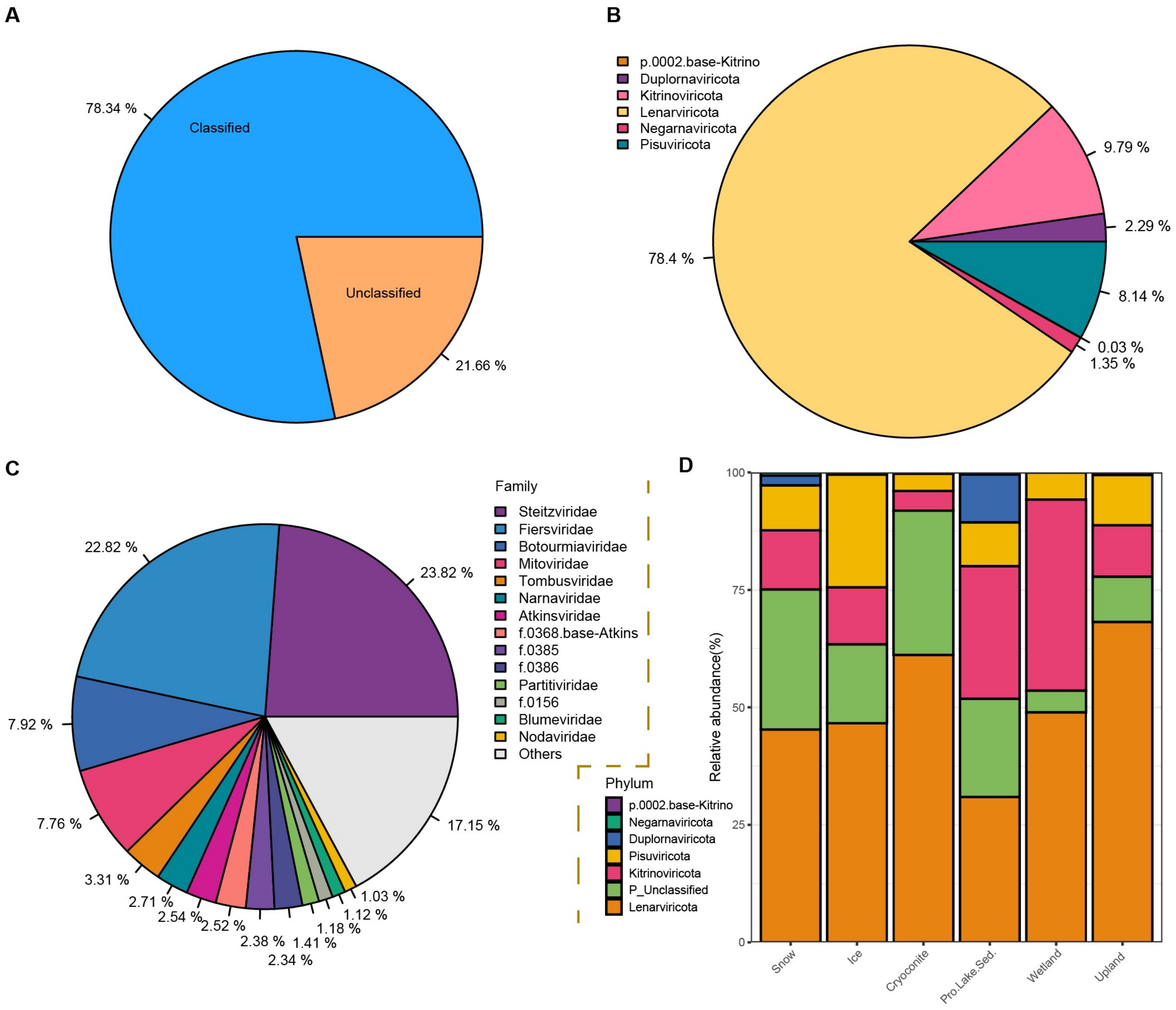


**Fig. S6: Proportion of classified and unclassified RNA viral genomes and the relative abundance of vOTUs at Phylum-level. (A)** Proportion of classified and unclassified vContig. (**B**) proportion of vContigs belonging to established phylum, with the proportion of all classified vContigs scaled up to 100% . **(C)** proportion of vContigs belonging to established dominant families, with the proportion of all classified vContigs scaled up to 100%. **(D)** the average relative abundance of each phylum across habitats and ecosystems.

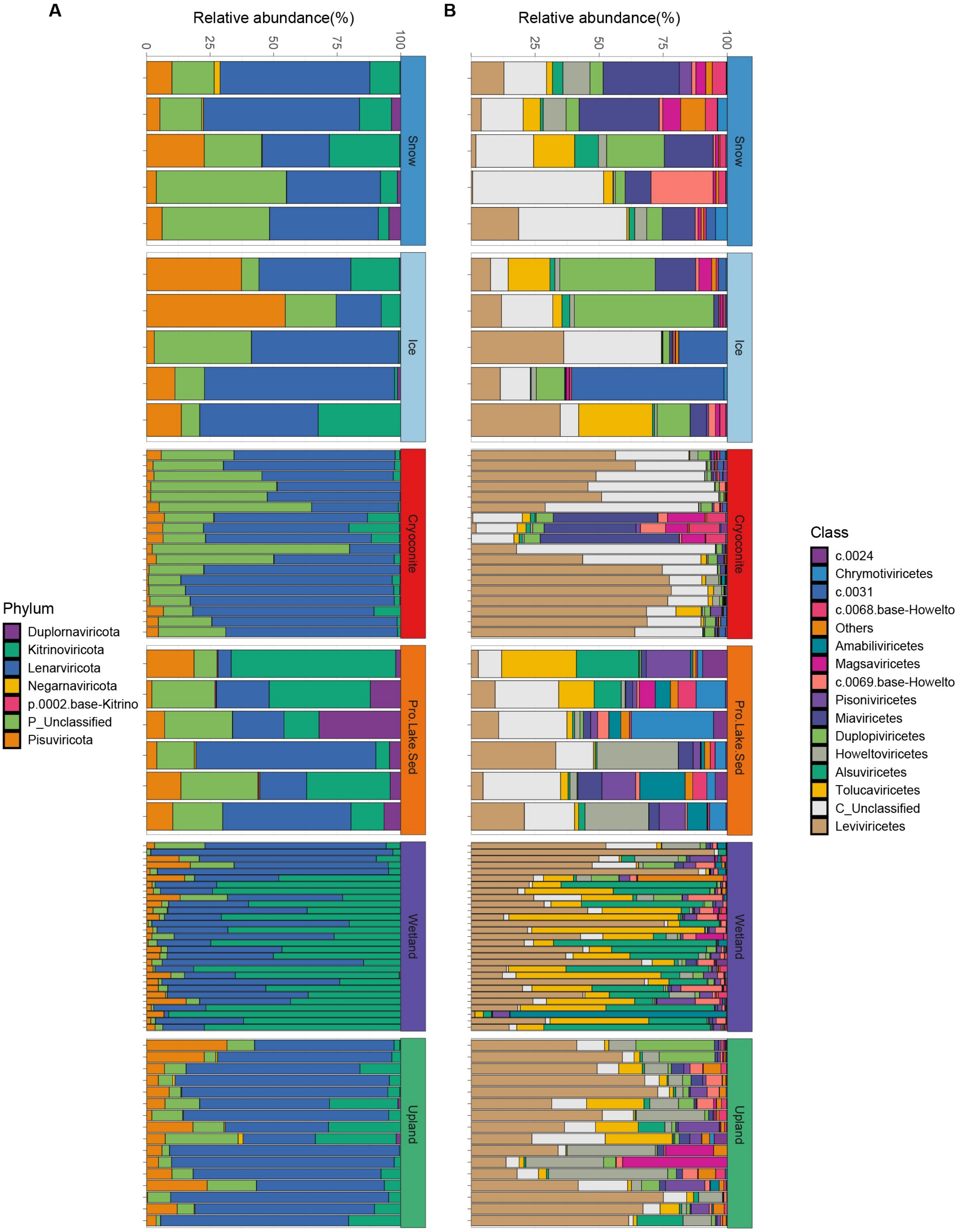


**Fig. S7. The relative abundance of RNA viral Phylum and Class across samples.** The relative abundance of dominant RNA viral Phylum (**A**) and Class (**B**) across samples, respectively.

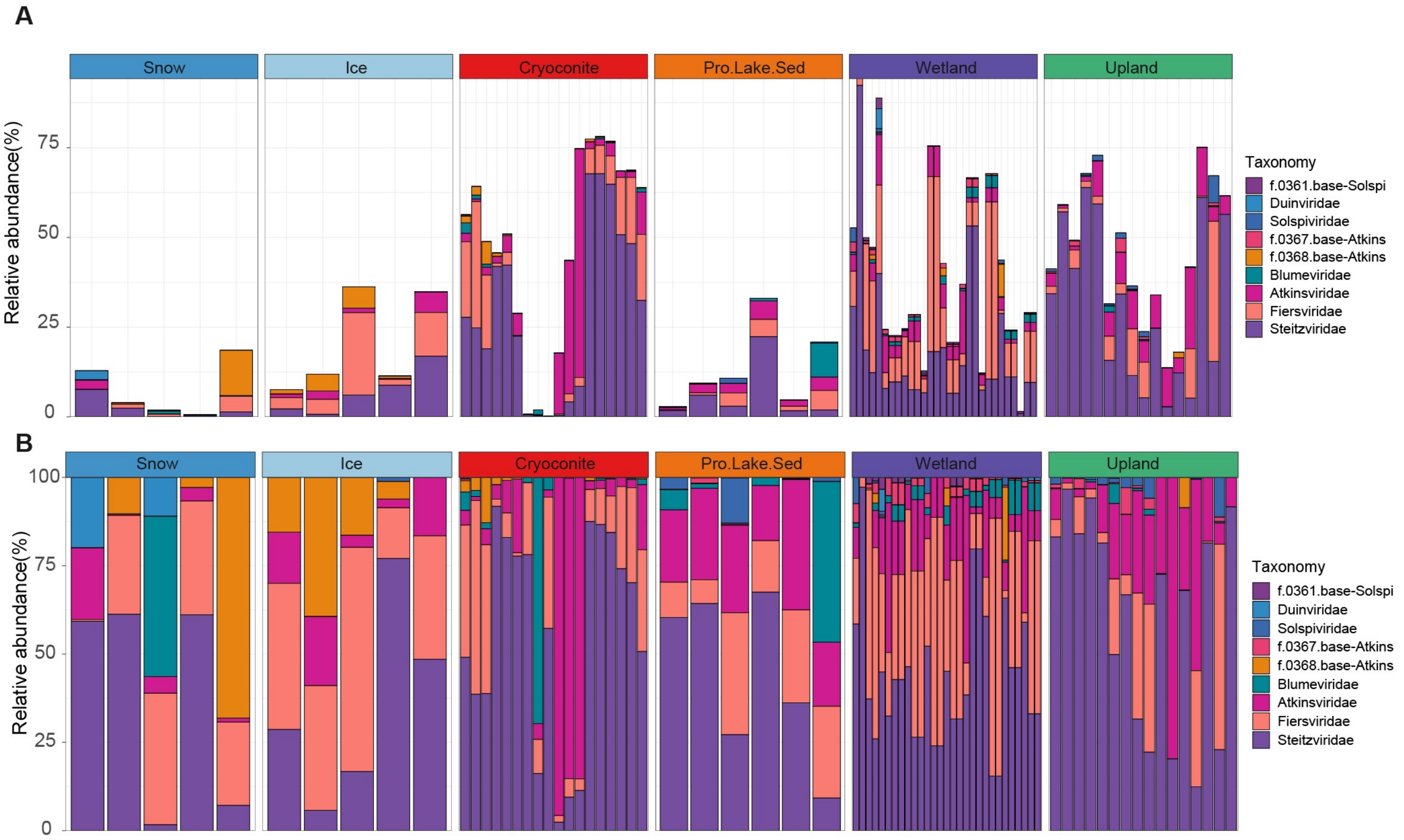


**Fig. S8. The relative abundance of vOTUs belonging to the RNA viral class *Leviviricetes*. (A)** The relative abundance of *Leviviricetes* families among all families (related to main **Fig. 2B**). **(B)** The relative proportion of *Leviviricetes* families abundance among all *Leviviricetes* families scaled up to 100%.

**A**

| | Snow | Ice | Cryoconite | Pro.Lake.Sed. | Wetland | Upland |
|---|---|---|---|---|---|---|
| Snow | | | | | | |
| Ice | 0.154 | | | | | |
| Cryoconite | 0.13 | 0.137 | | | | |
| Pro.Lake.Sed. | 0.216 | 0.252 | 0.197 | | | |
| Wetland | 0.099 | 0.103 | 0.192 | 0.117 | | |
| Upland | 0.085 | 0.1 | 0.135 | 0.116 | 0.095 | |

**B**

| | Glacier | Pro.Lake.Sed. | Wetland | Upland |
|---|---|---|---|---|
| Glacier | | | | |
| Pro.Lake.Sed. | 0.114 | | | |
| Wetland | 0.147 | 0.117 | | |
| Upland | 0.084 | 0.116 | 0.095 | |

**Fig. S9. $R^2$ value of pair-wise adonis analysis of TPC RNA viral communities. (A)** pair-wise of glacier habitats and ecosystems. (**B**) pair-wise analysis of ecosystems. red, $P<0.001$, black, $P<0.05$, dark blue, $0.05<P<0.1$. Total $R^2$ equals to 0.23 and 0.174 for habitats and ecosystems, respectively, with both $P<0.001$.

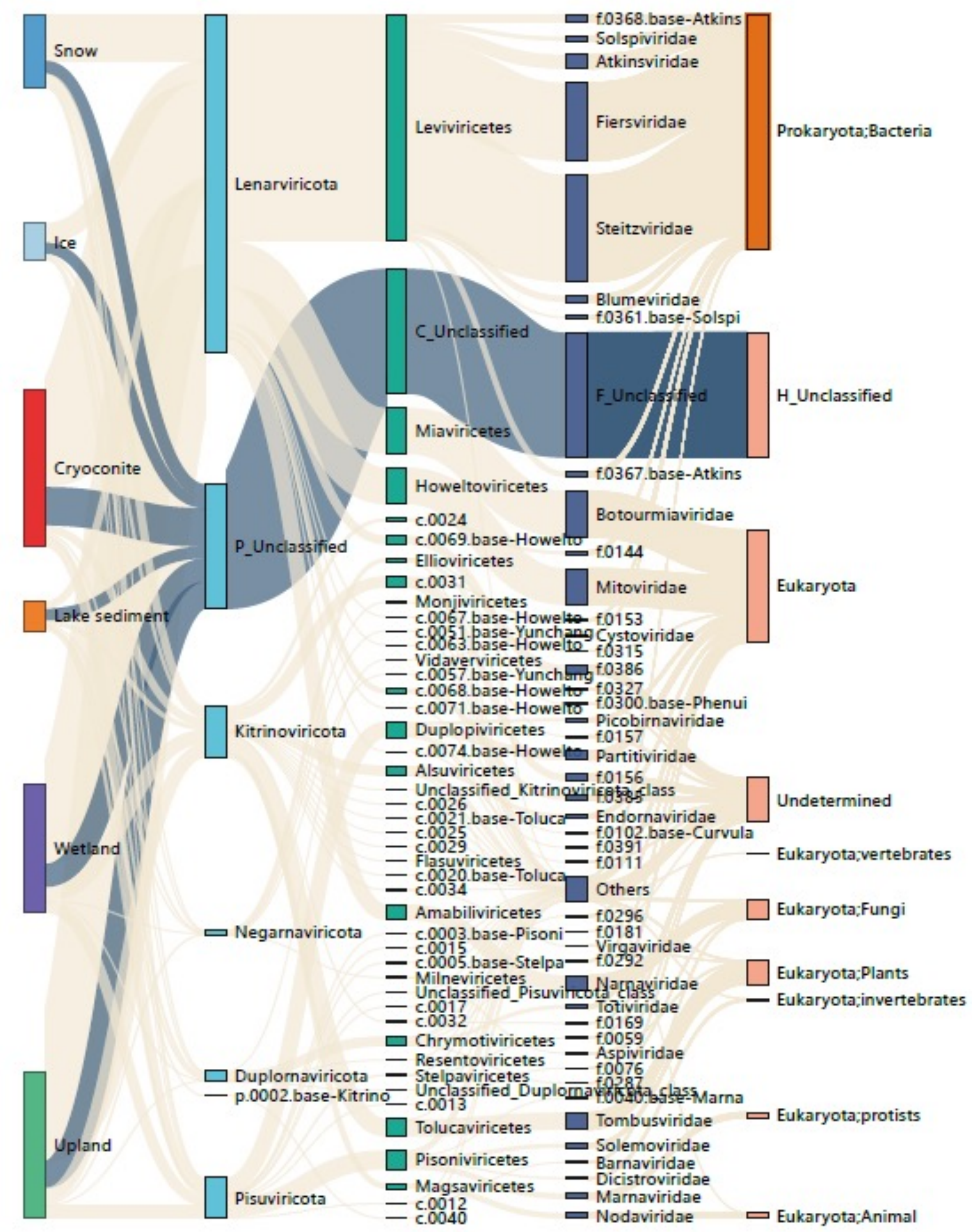


**Fig. S10. Sankey plot showing the distribution of vContigs in habitats, phyla, classes, families and the types of their corresponding hosts.**

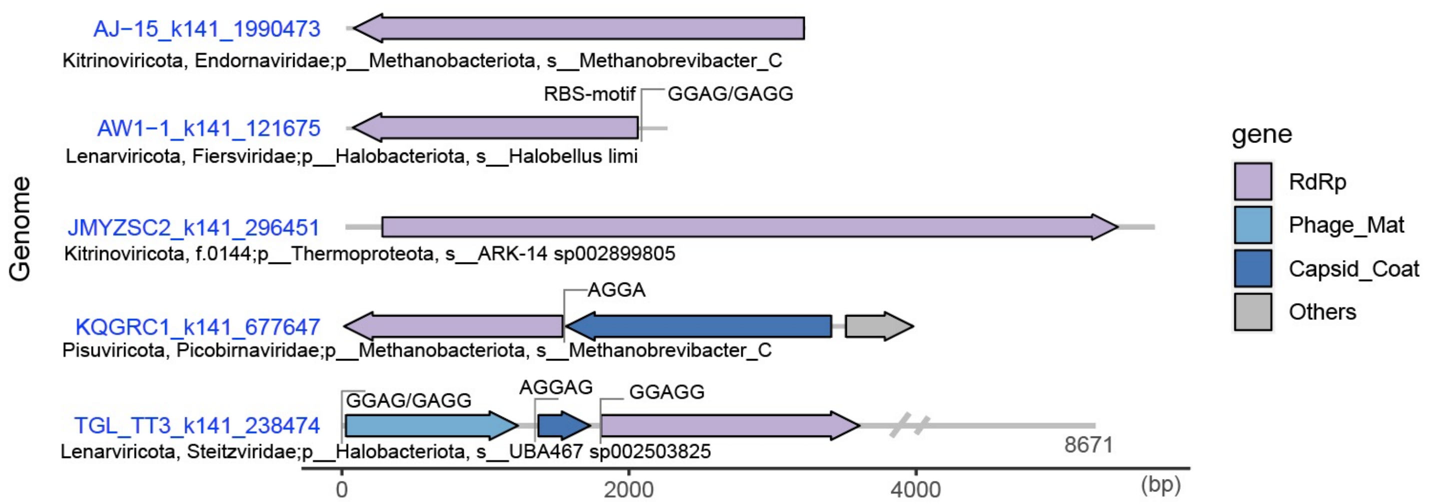


**Fig. S11. Genome architecture of putative archaeal RNA viruses.**

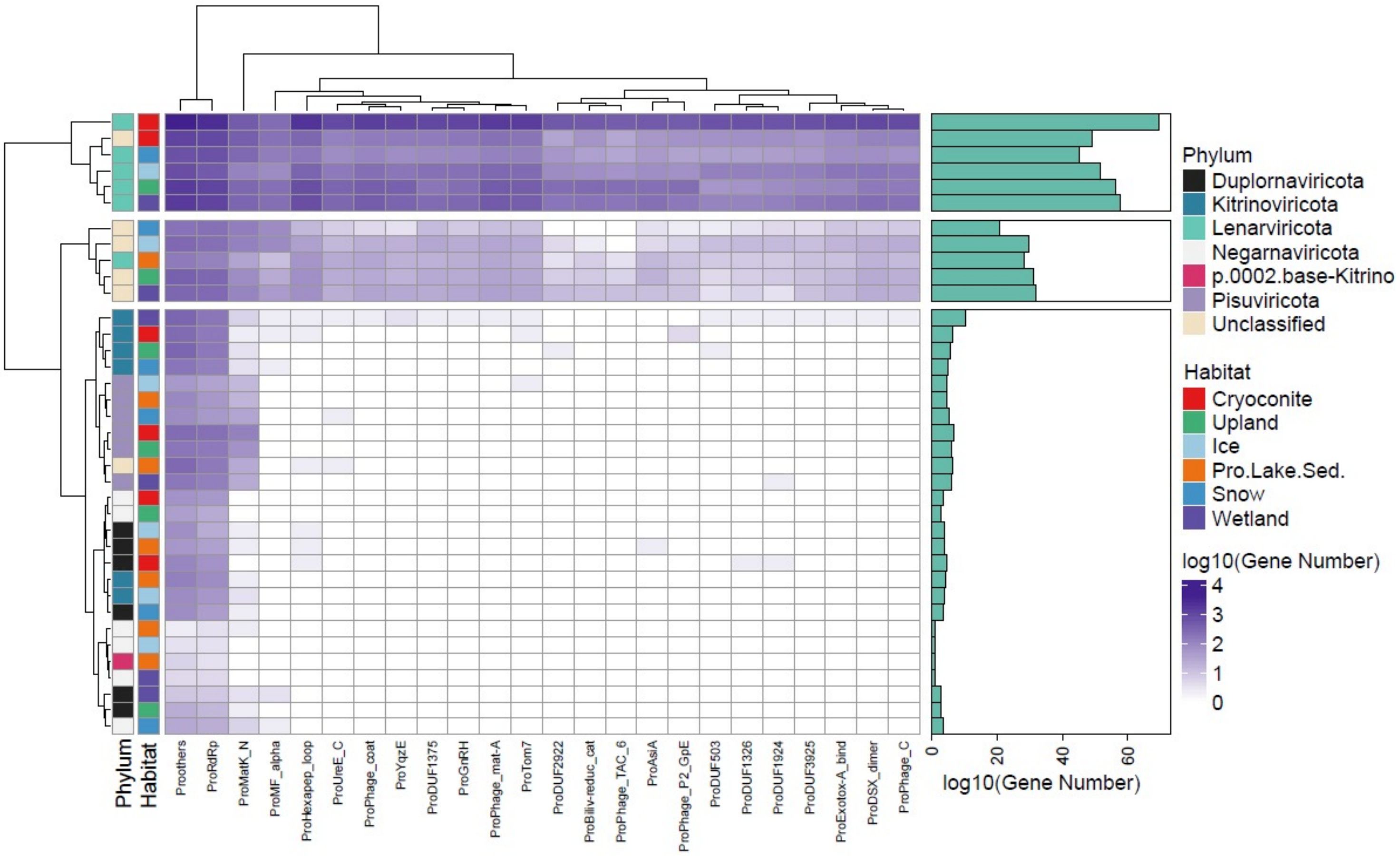


**Fig. S12. Functional profile of TPC RNA viral genomes**

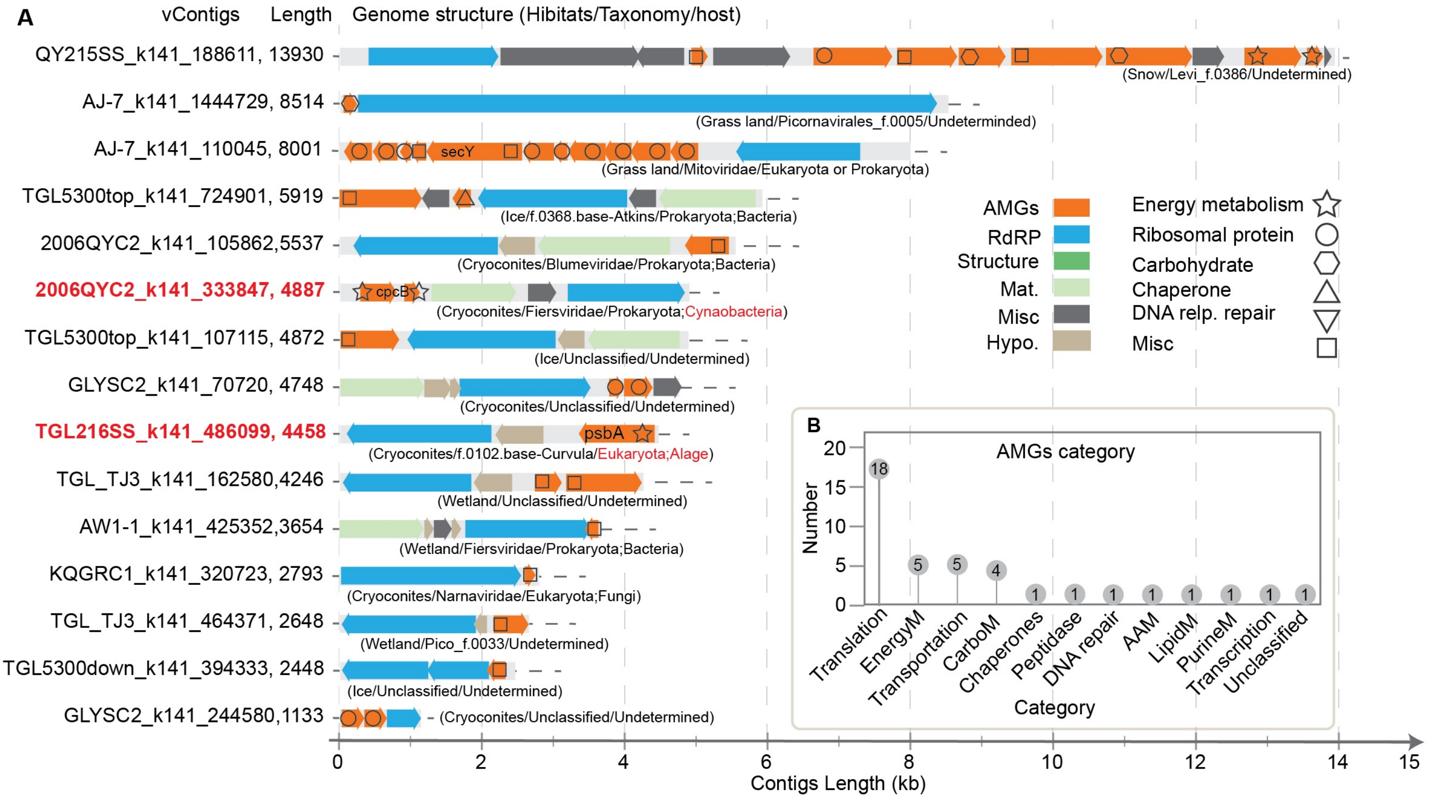


**Fig. S13. Genome structure of vContigs with AMGs and number of AMGs belonging to different functional categories. (A)** Genome structure of vContigs with AMGs. **(B)** number of AMGs belonging to different functional categories.

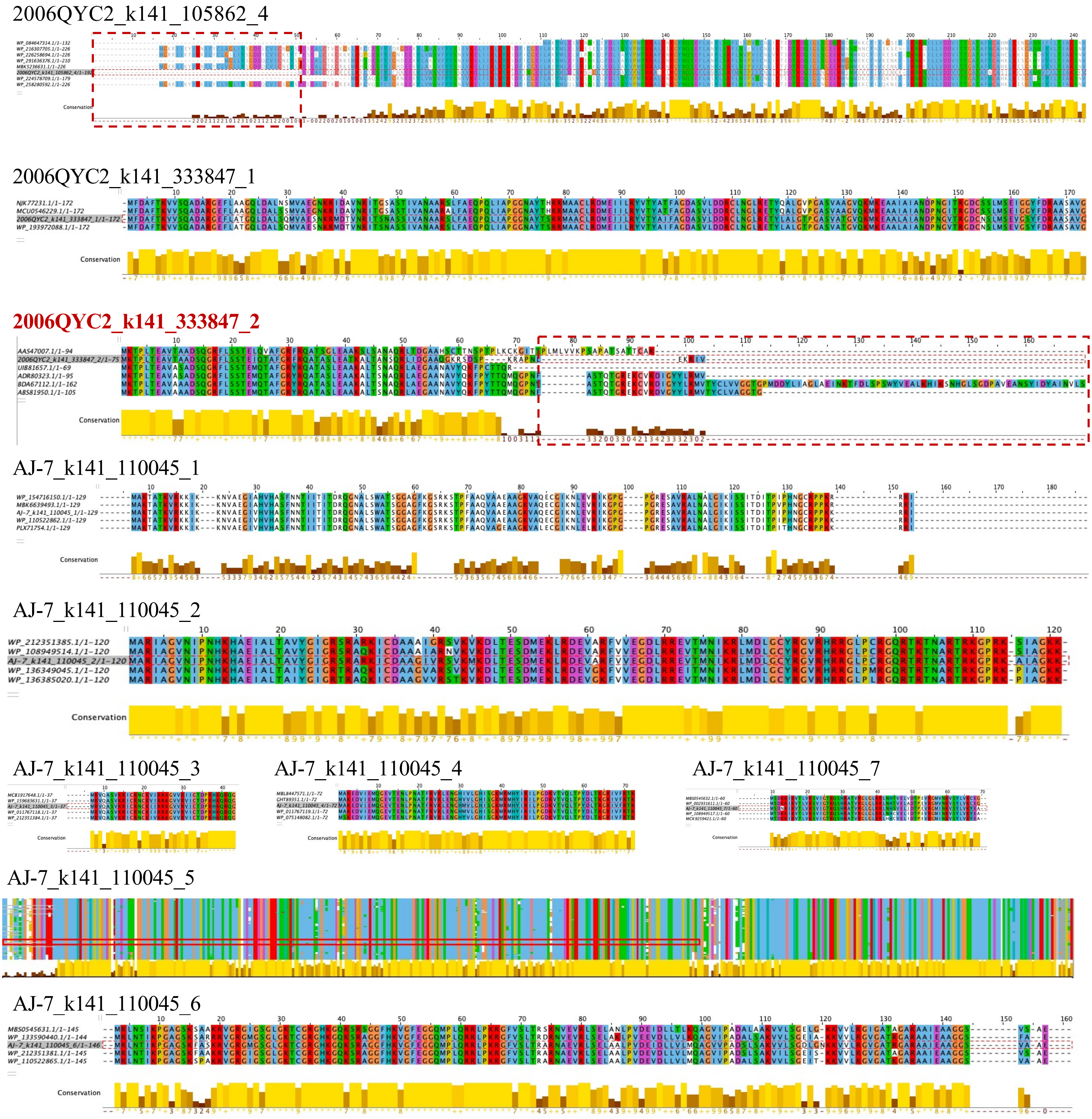


**Fig. S14. Sequences alignments of putative AMGs amino acid sequences with their closest cellular homologs found in NR database .** AMGs amino acid sequences that were only partial of the full ORF compared to the references are highlighted in red.

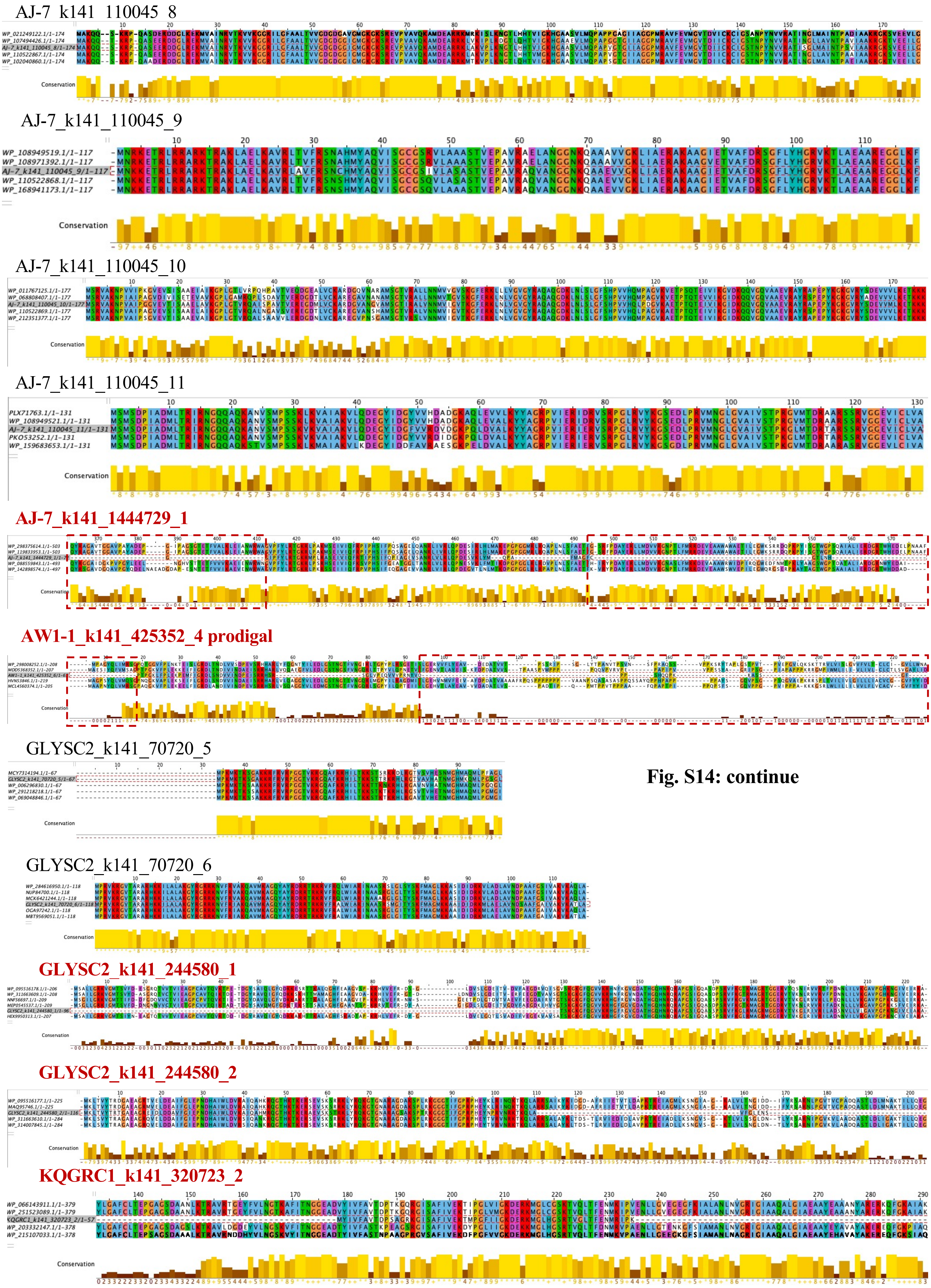


**Fig. S14: continue**

## QY215SS_k141_188611_2

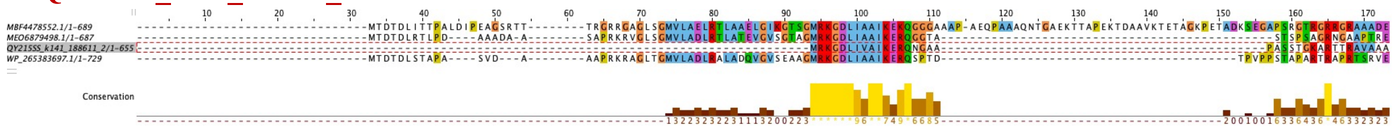

## QY215SS_k141_188611_4

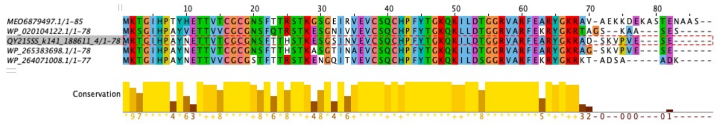

## QY215SS_k141_188611_5

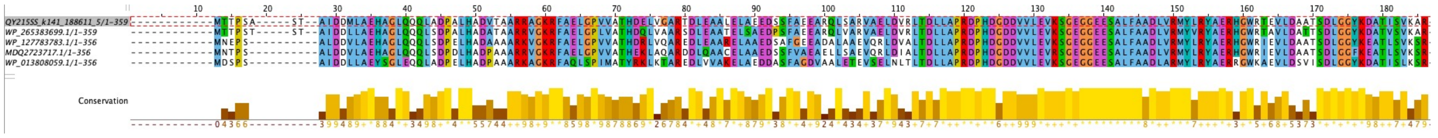

## QY215SS_k141_188611_6

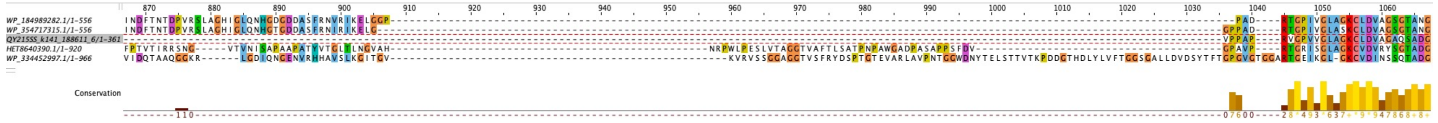

## QY215SS_k141_188611_7

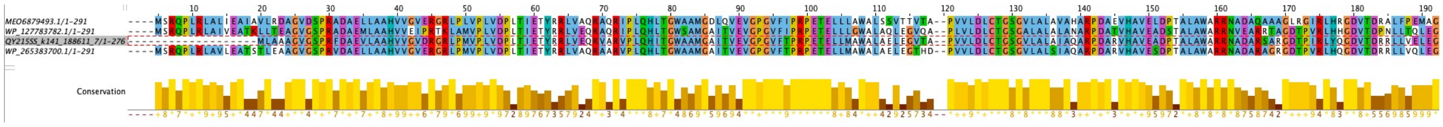

## QY215SS_k141_188611_8

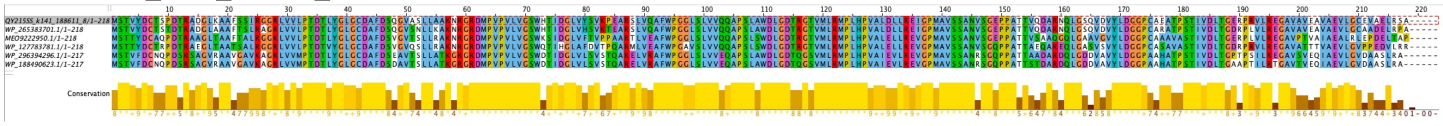

## QY215SS_k141_188611_9

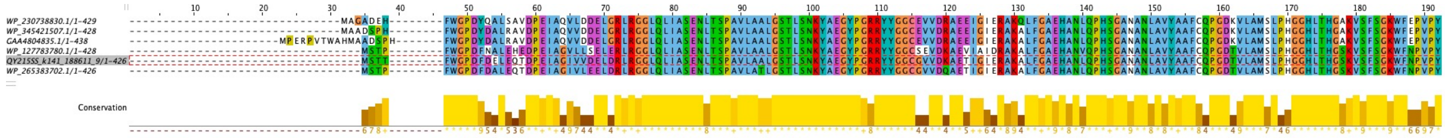

## QY215SS_k141_188611_10

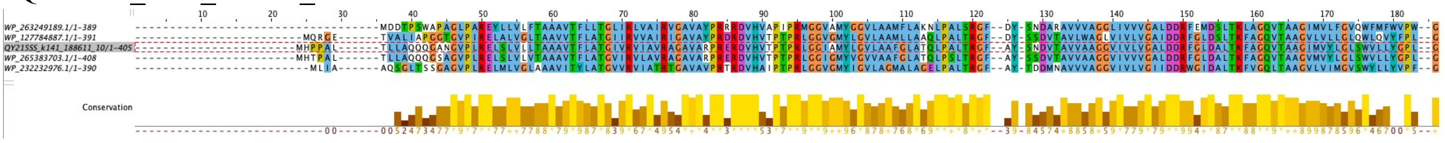

## QY215SS_k141_188611_12

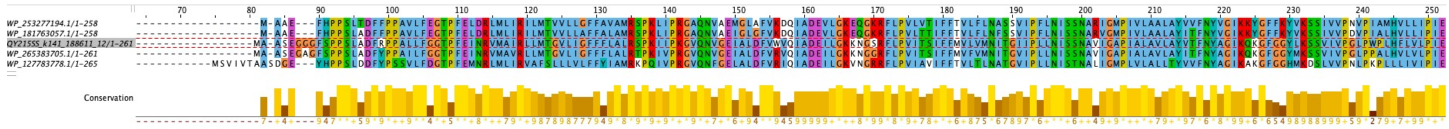

## QY215SS_k141_188611_13

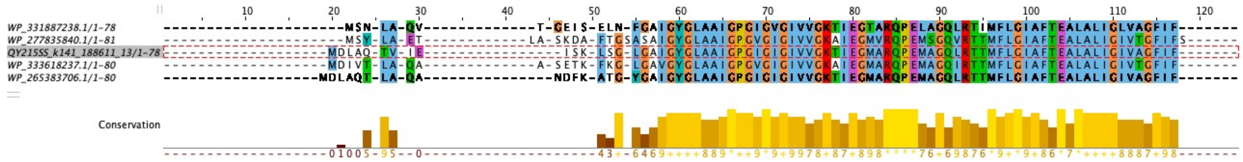

**Fig. S14: continue**

TGL_TJ3_k141_162580_3

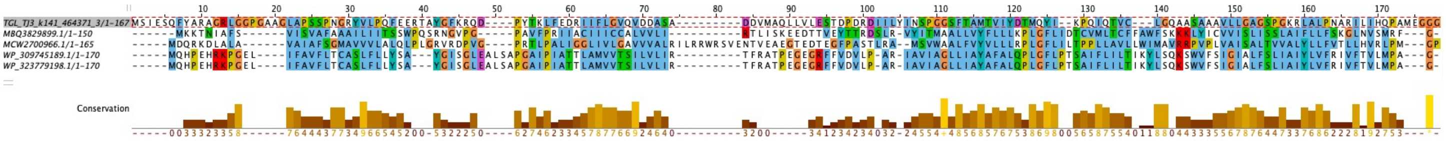

TGL_TJ3_k141_162580_4

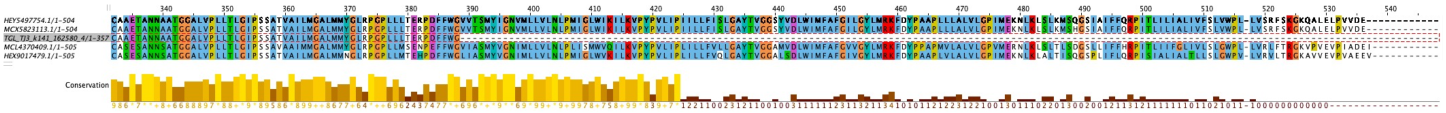

TGL_TJ3_k141_464371_3

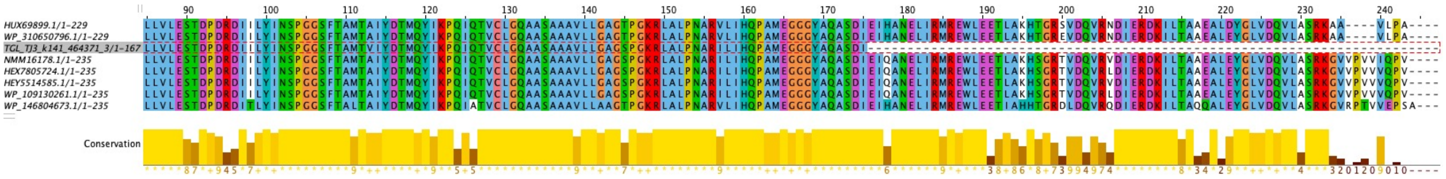

TGL216SS_k141_486099_3

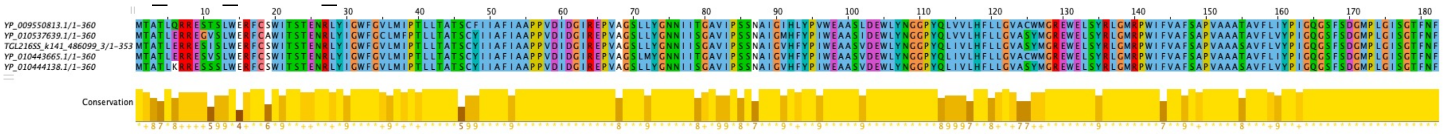

TGL5300down_k141_394333_3

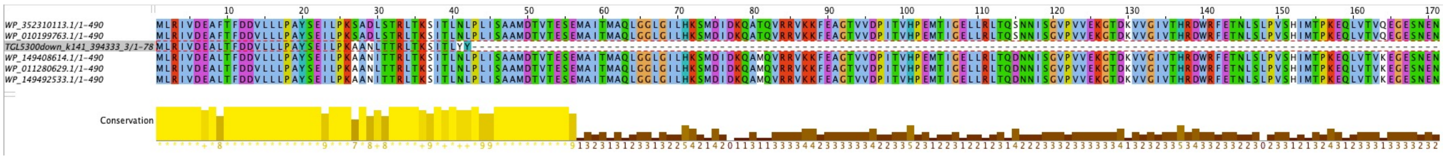

TGL5300top_k141_107115_1

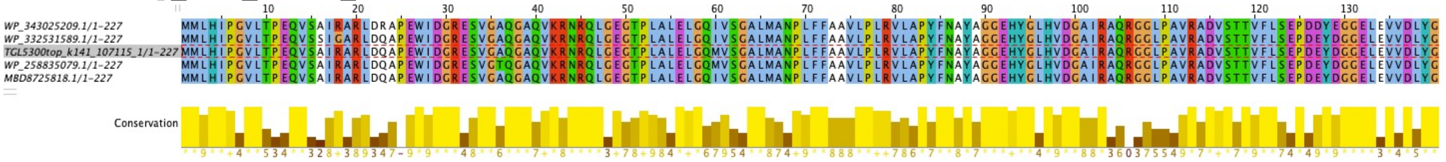

TGL5300top_k141_724901_1

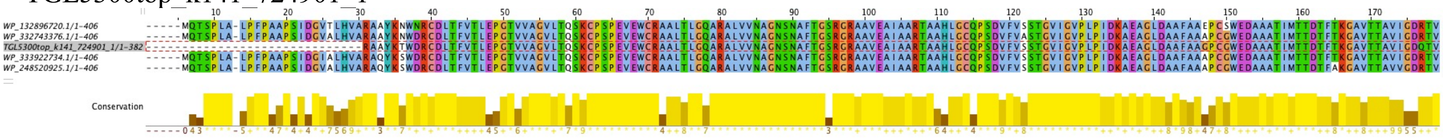

TGL5300top_k141_724901_3

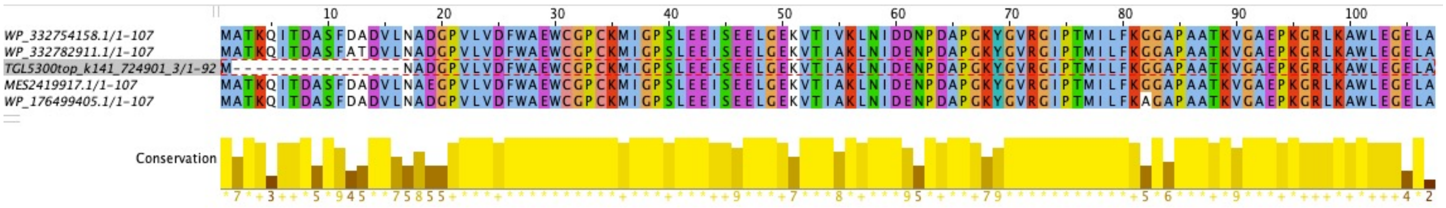

**Fig. S14: continue**

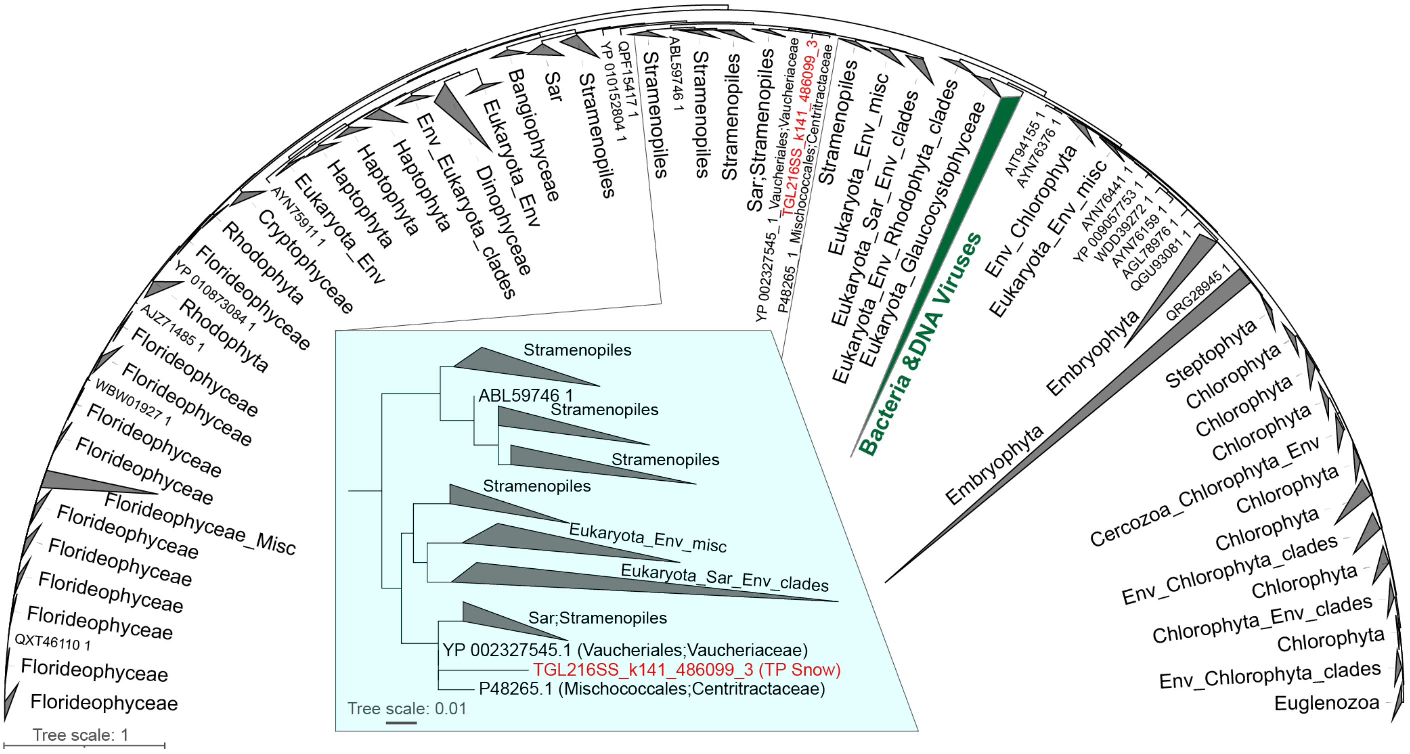


**Fig. S15. Maximum-likelihood tree of the protein sequences of PsbA1.** PsbA1 from the vContigs is highlight in red and the sub-clade with this sequences is showed in the inserted panel. The clade of PsbA1 protein encoded by Bacteria and DNA viruses is colored in deep green. The RNA viruses coded by vContigs from TP snow is closely related to Stramenopiles (Sar supergroup), Vaucheriaceae (a family in Chromista, brown algae and allies, YP_002327545.1) and Centritractaceae (a family in Xanthophyceae, yellow-green algae, P48265.1). The maximum-likelihood tree was constructed by using the FastTree software with the parameters "-gamma -lg -boot 1000". Reference sequences were obtained from NCBI with IPG database by searching with "photosystem II protein D1" and filtered out none PsbA1 domain using hmmsearch with TIGR01151.1.hmm and PF00124.hmm. The gathering score 27 and length of 180 aa were used for filtering. After filtering, all reference PsbA1 aa sequences were dereplicated using CD-HIT with 0.99 as cutoff. PsbA1 and non-redundant reference PsbA1 were combined for muscle alignment, trimal (with -gappyout) and gaps removal (drop sequences with >70% gaps). The final multiple sequence alignments were used for tree construction.

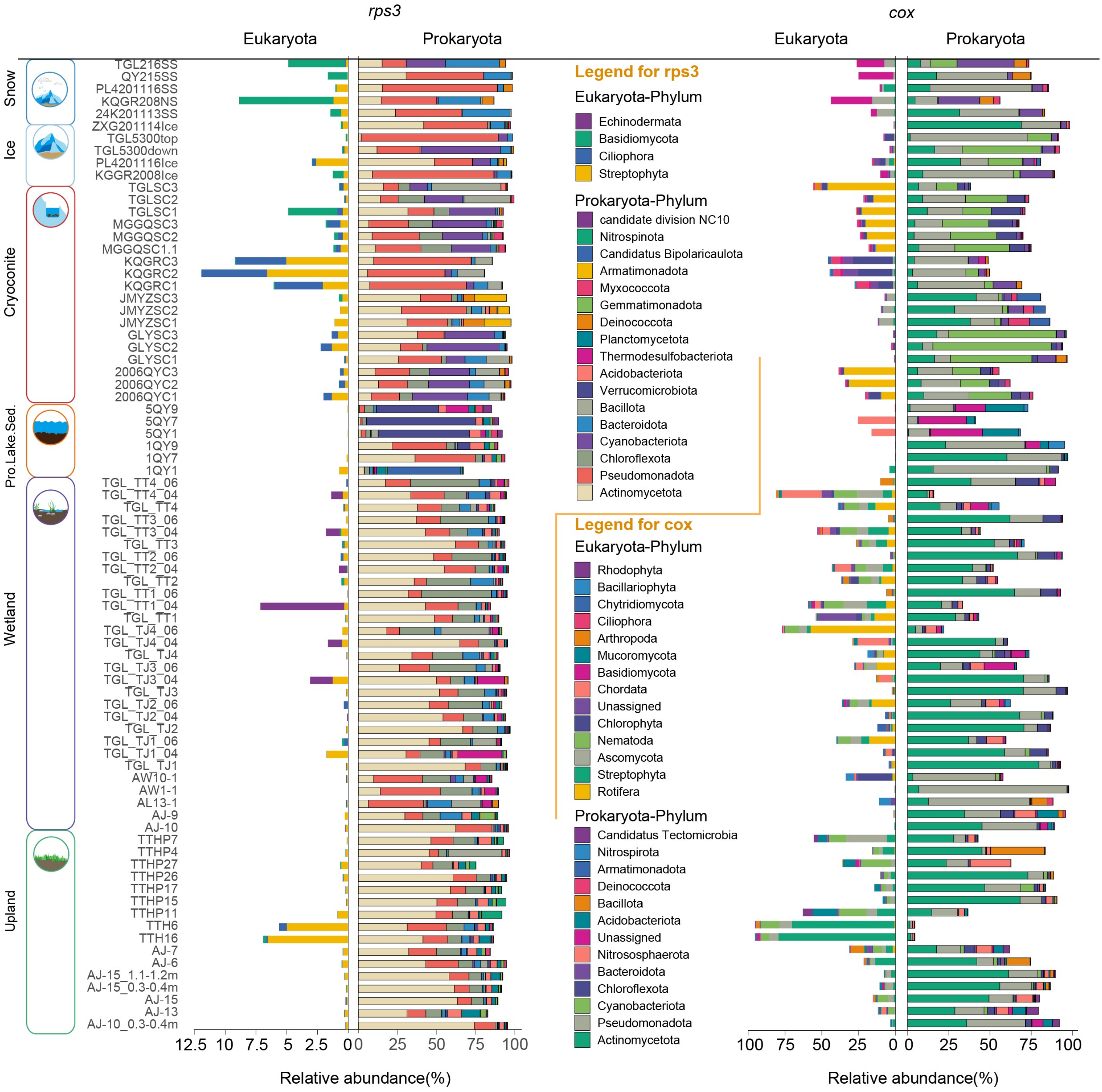


**Fig. S16. The relative abundance of *rps3* gene and *cox1* gene transcripts across habitats, ecosystems and samples.**

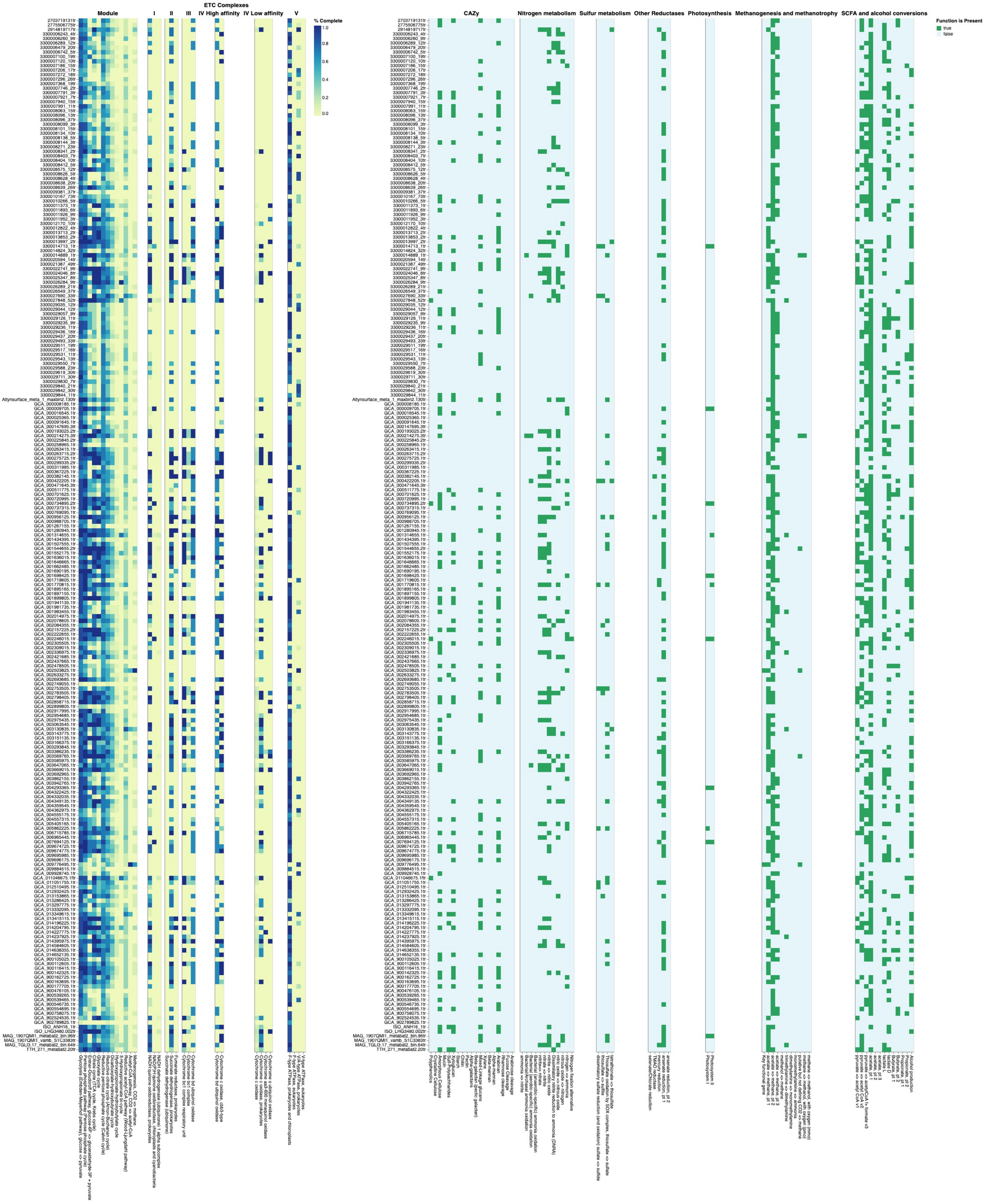

**Fig. S17. Genome annotation of MAGs with at least one CRISPR-linkage to RNA vContigs.**

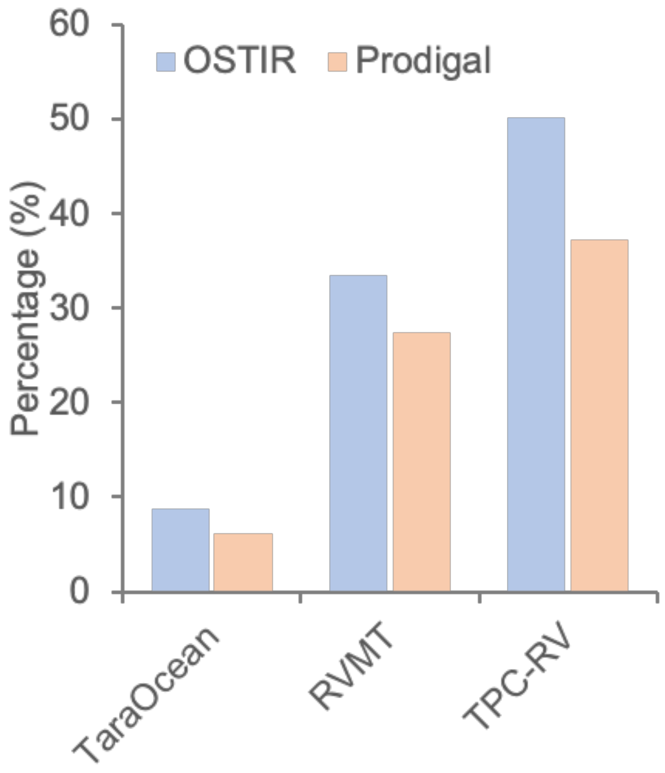


**Fig. S18. Percentage of RNA viral genomes with cd ratio ≥ 50%.** Two pipelines were used to identify the RBS motifs within each vContigs, including OSTIR (Roots, C. T. et al., 2021, J Open Source Softw, 10.21105/joss.03362) and prodigal (Hyatt et al., 2010, BMC Bioinformatics, 10.1186/1471-2105-11-119).

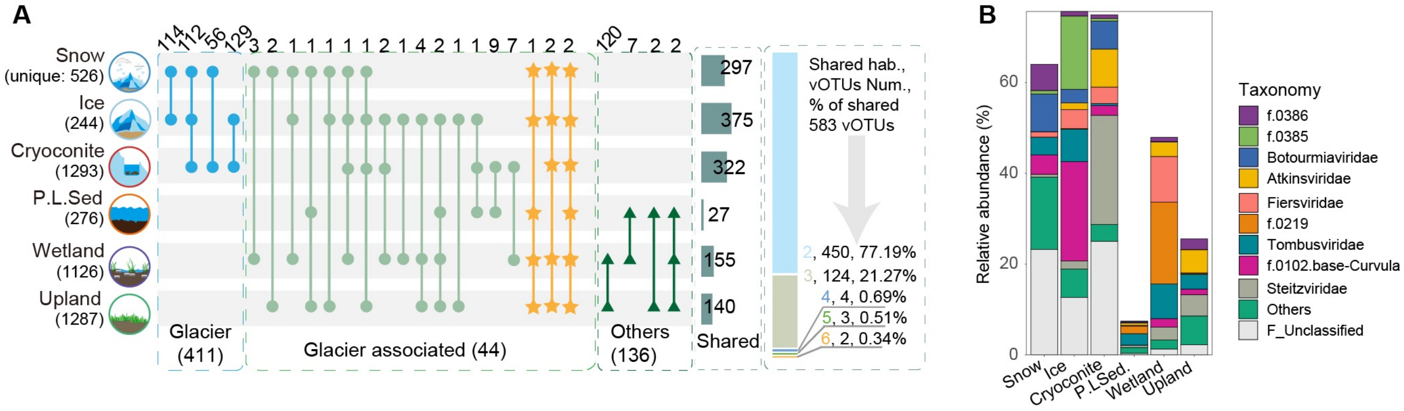


**Fig. S19. Shared number of vOTUs across different habitats.**

(**A**). Upset plot showing the number of shared and unique RNA vOTUs across Ecosystems and Habitats. (**B**) Relative abundance of shared vOTUs. The relative abundance of those shared vOTUs was grouped at the family levels.

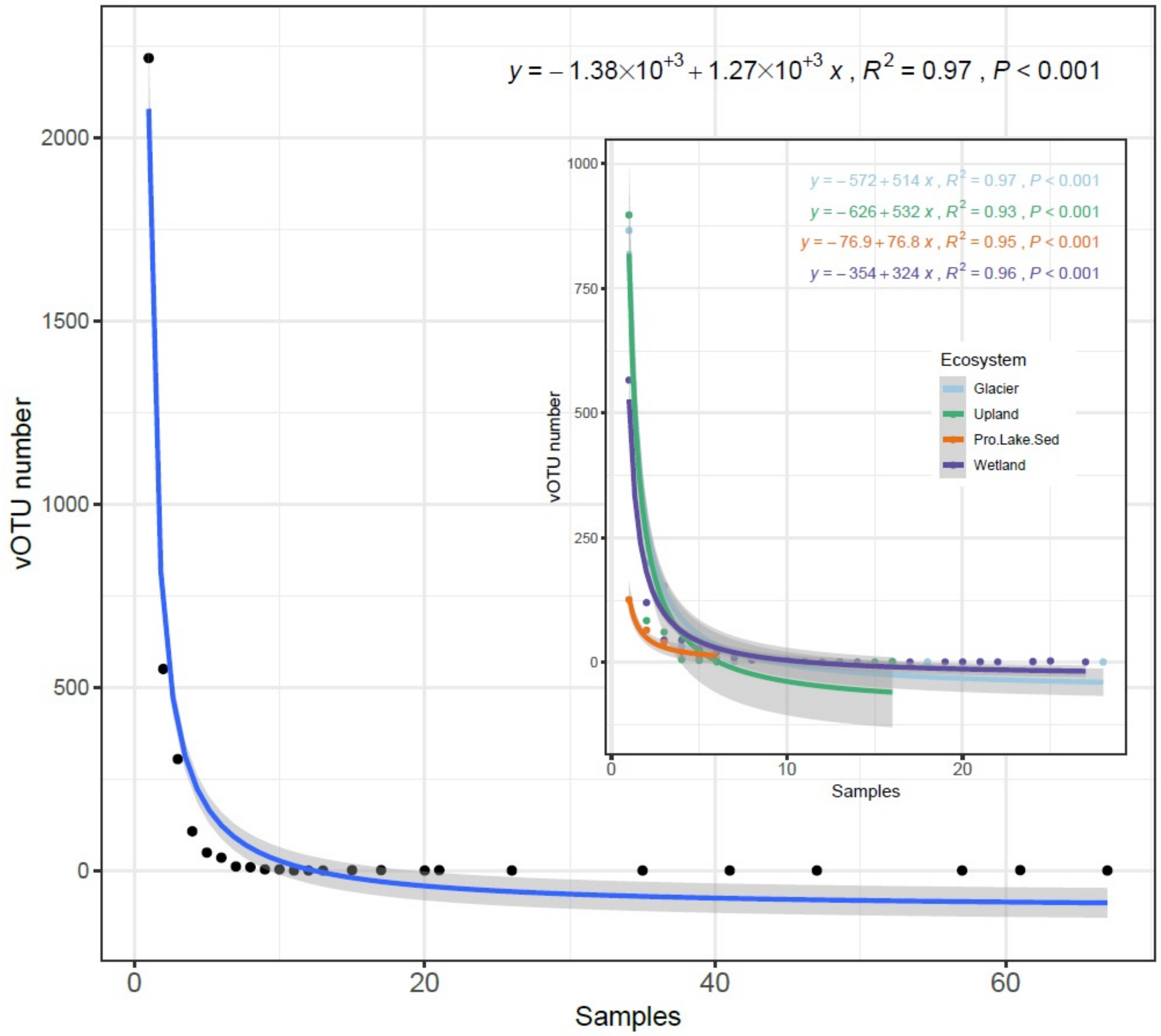


**Fig. S20. Shared number of vOTUs as a function of samples.**

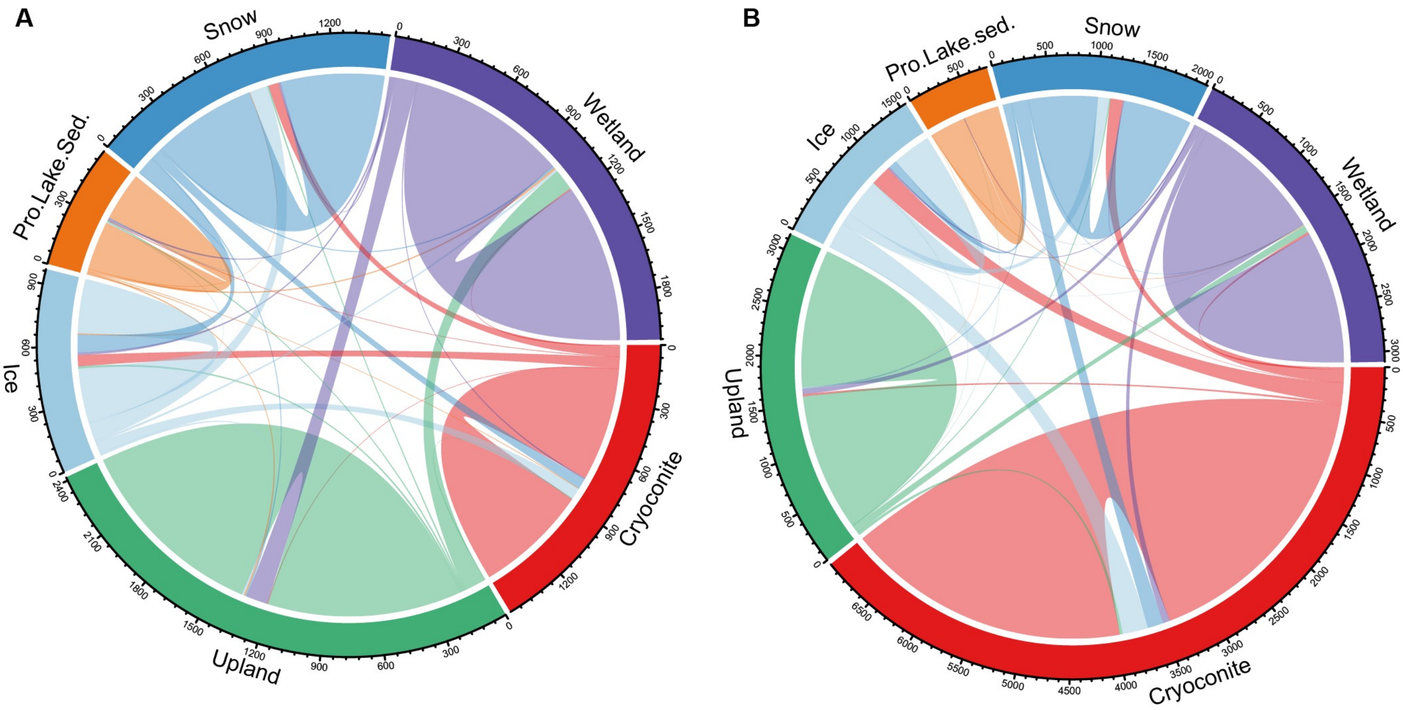


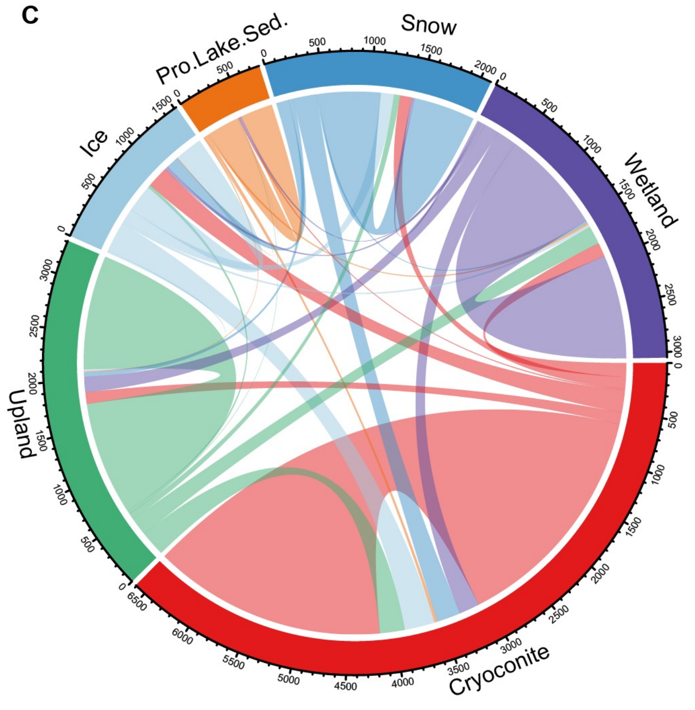


**Fig. S21. Sharing of vOTUs among different habitats and ecosystems at different levels. (A)** based on TPM >=1000 as a threshold for the occurrences of one vOTUs in certain environments. **(B)** based on uclust clustering with cutoff 0.7 (~genus level) and **c** based on UCLUST clustering with cutoff 0.5 (family to subfamily levels)

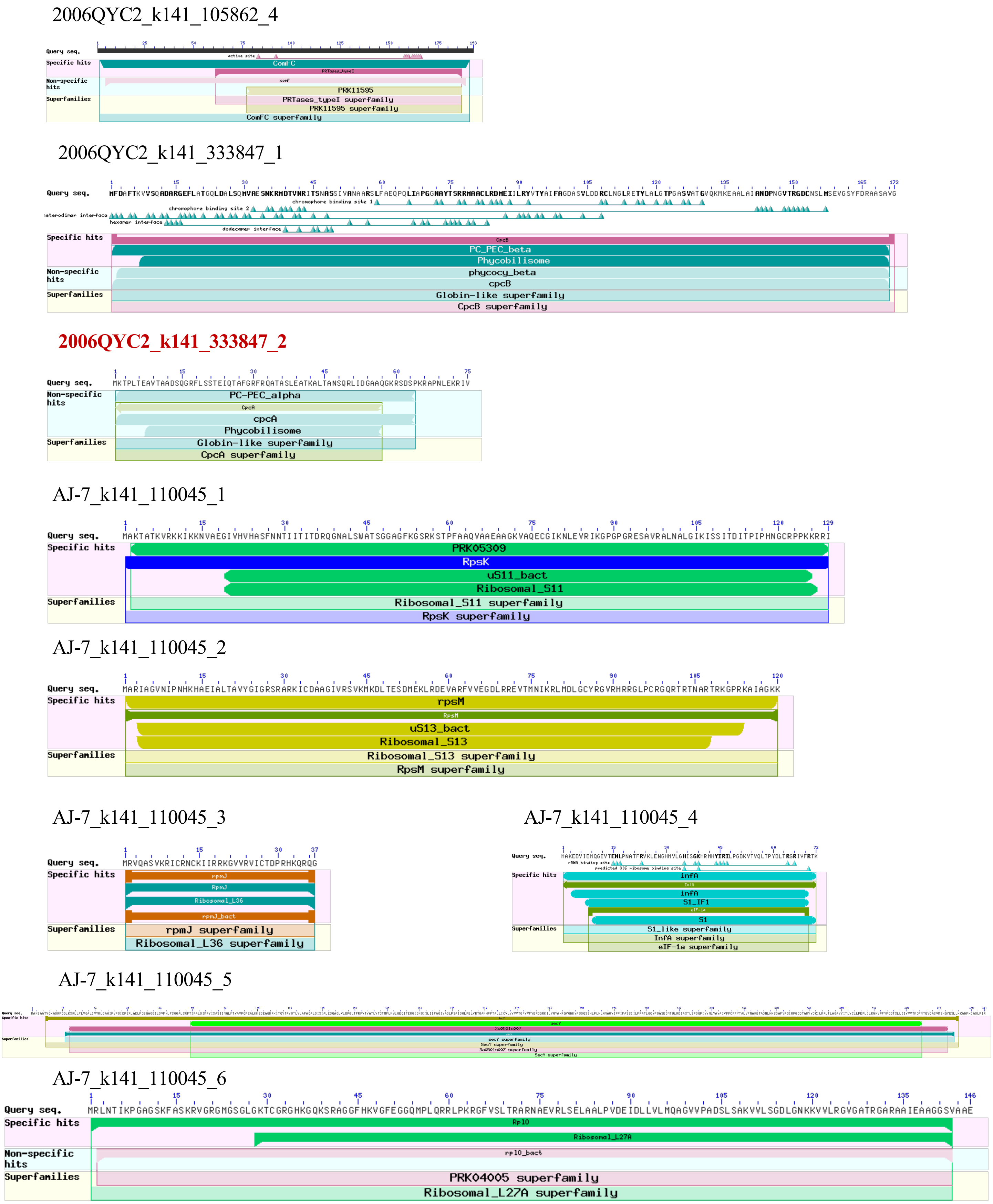


**Fig. S22. Domain analysis of putative AMGs.** AMGs amino acid sequences were searched against the (https://www.ncbi.nlm.nih.gov/Structure/cdd/wrpsb.cgi) for detailed functional domain annotation.

AJ-7_k141_110045_7

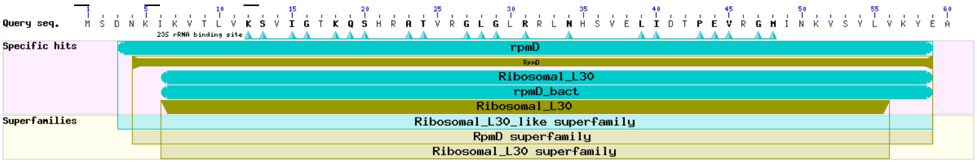


AJ-7_k141_110045_8

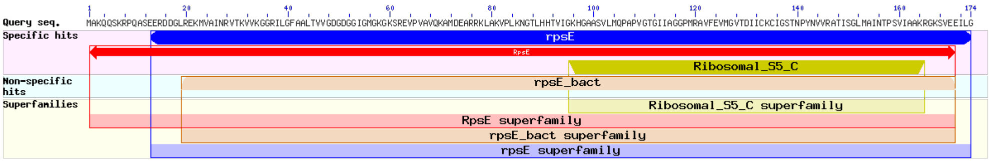


AJ-7_k141_110045_9

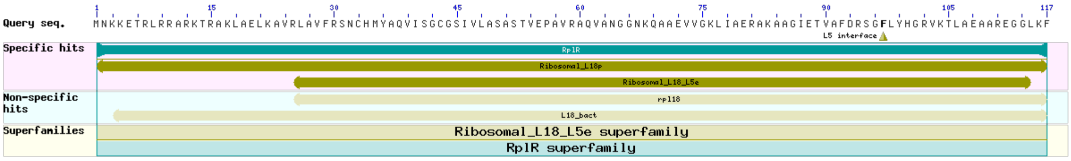


AJ-7_k141_110045_10

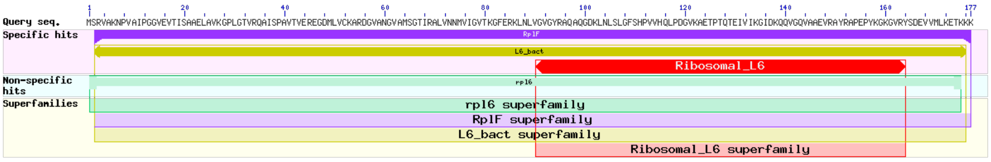


AJ-7_k141_110045_11

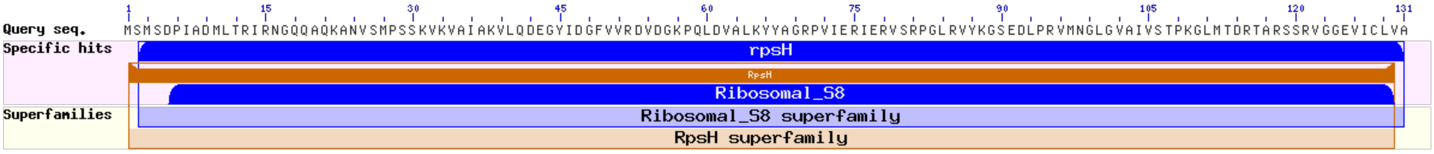


AJ-7_k141_1444729_1

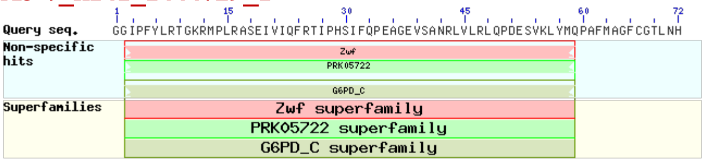


AW1-1_k141_425352_4

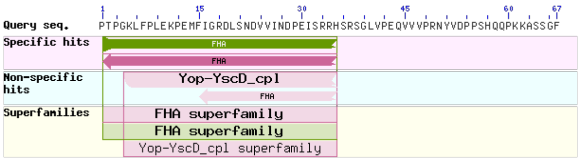


GLYSC2_k141_244580_1

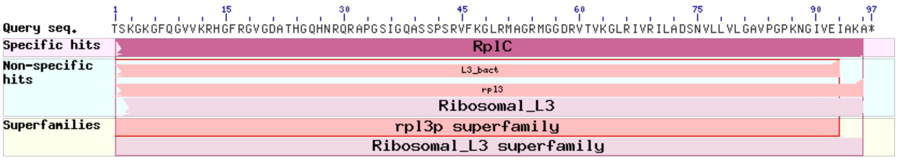


GLYSC2_k141_244580_2

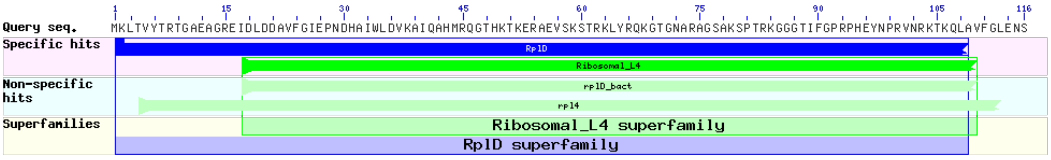


**Fig. S22: Continue.**

GLYSC2_k141_70720_5

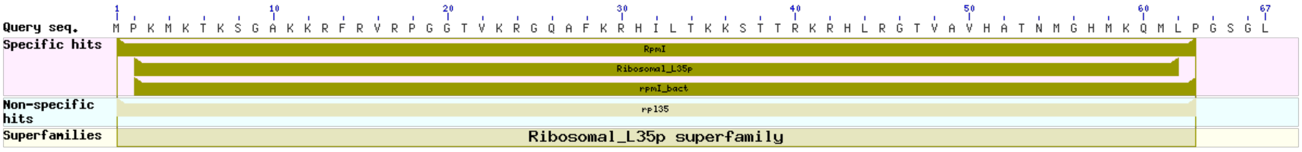


GLYSC2_k141_70720_6

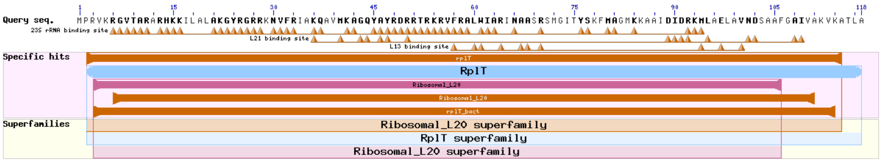


**KQGRC1_k141_320723_2**

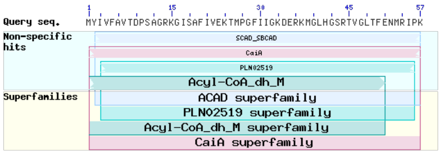


QY215SS_k141_188611_4

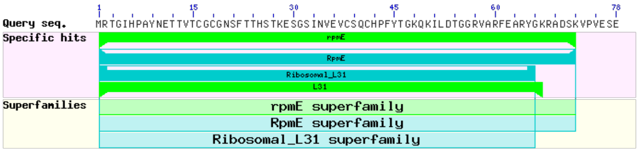


**QY215SS_k141_188611_2**

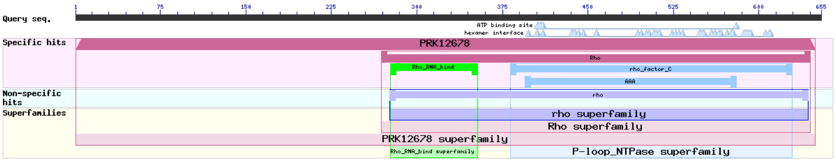


QY215SS_k141_188611_5

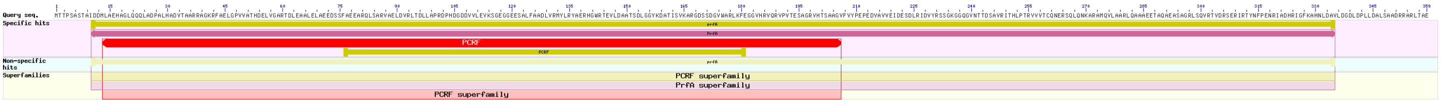


**QY215SS_k141_188611_6**

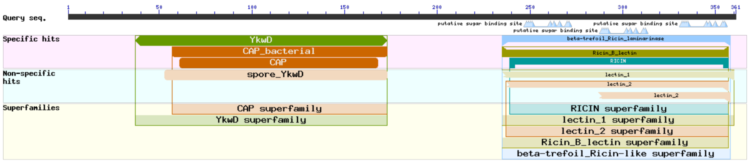


QY215SS_k141_188611_7

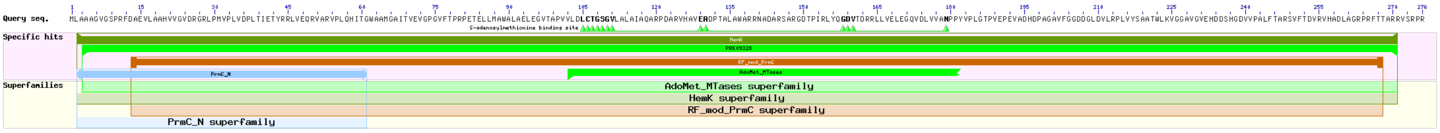


QY215SS_k141_188611_8

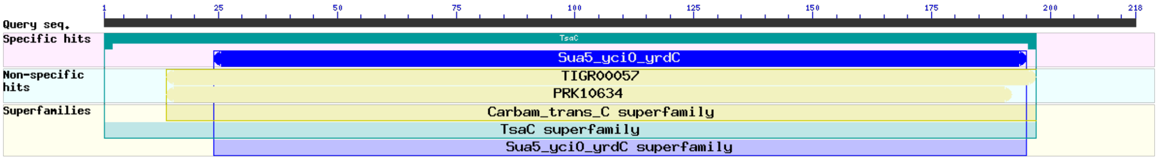


QY215SS_k141_188611_9

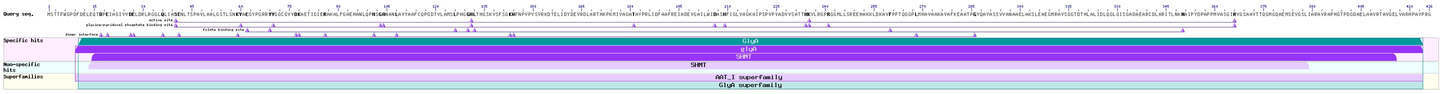


**Fig. S22: Continue.**

## QY215SS_k141_188611_10

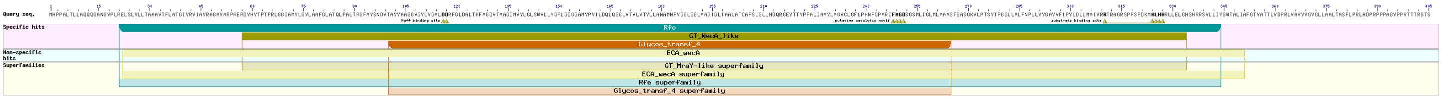


## QY215SS_k141_188611_12

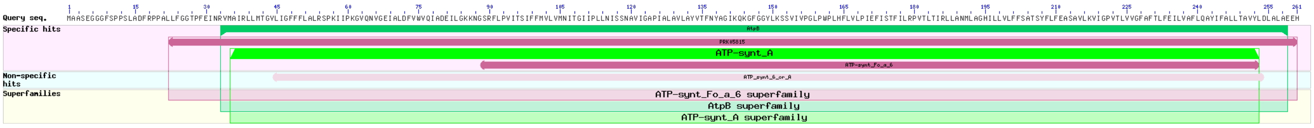


## QY215SS_k141_188611_13

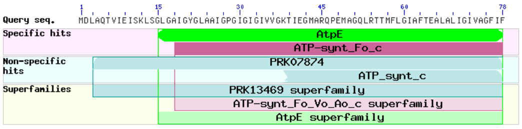


### TGL_TJ3_k141_162580_3

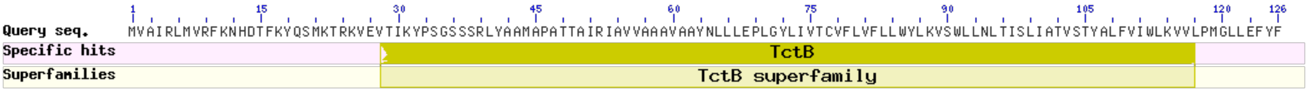


### TGL_TJ3_k141_162580_4

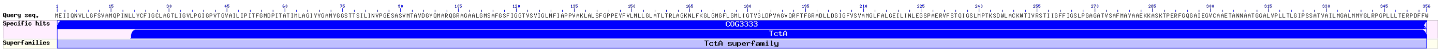


### TGL_TJ3_k141_464371_3

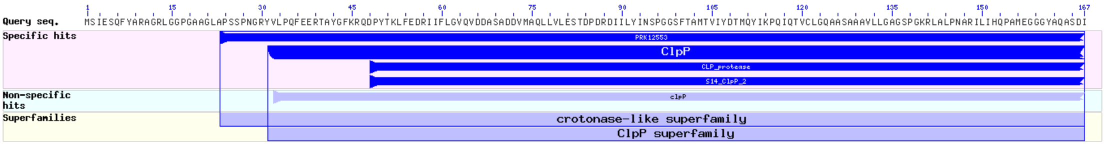


### TGL216SS_k141_486099_3

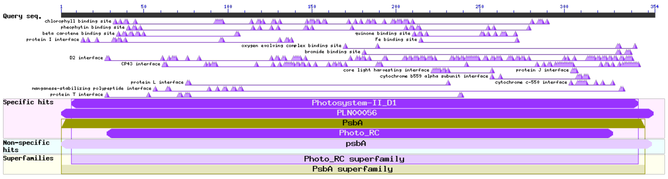


### TGL5300down_k141_394333_3

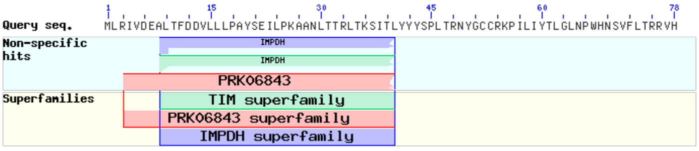


### TGL5300top_k141_107115_1

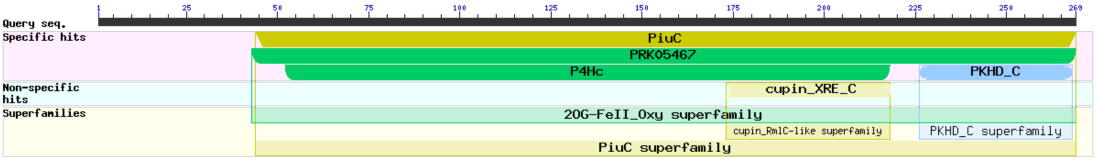


### TGL5300top_k141_724901_1

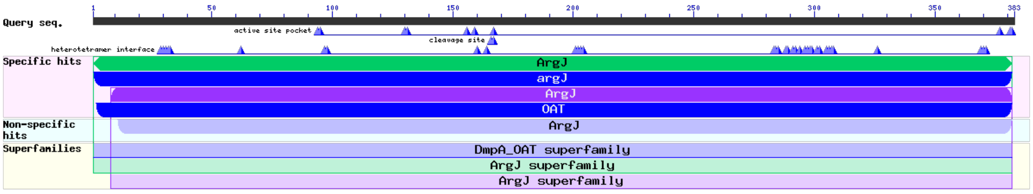


### TGL5300top_k141_724901_3

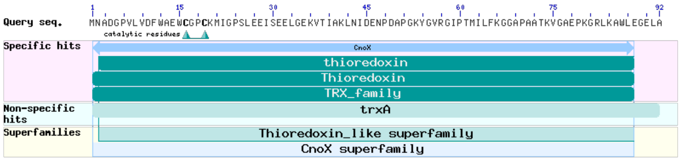


**Fig. S22: Continue.**
